# Cooperativity in *E. coli* Aspartate Transcarbamoylase is Tuned by Allosteric Breathing

**DOI:** 10.1101/2024.11.19.624407

**Authors:** Robert C. Miller, Michael G. Patterson, Neti Bhatt, Xiaokun Pei, Nozomi Ando

**Author notes:** These authors contributed equally (Robert C. Miller, Michael G. Patterson).

## Abstract

Aspartate transcarbamoylase (ATCase) from *Escherichia coli* catalyzes a key step in pyrimidine nucleotide biosynthesis and has long served as a model for allosteric regulation. Despite decades of study, how nucleotide binding at distant regulatory sites controls cooperativity between active sites remained unresolved. Here we show that ATCase does not simply interconvert between two conformations, as traditionally depicted, but instead samples a continuum of conformations that tune enzyme cooperativity. Using complementary cryo-electron microscopy, small-angle X-ray scattering, and crystallography under conditions that ensure full assembly of the allosteric sites, we show that ATCase behaves like a flexible balloon whose global “breathing” motions directly regulate activity: compression enforces high cooperativity, inhibiting the enzyme, whereas expansion relieves this cooperativity and activates the enzyme. We further show that all four ribonucleoside triphosphates act in symmetric pairs to tune this motion, with the pyrimidines CTP and UTP compressing the enzyme to limit further pyrimidine production, and the purines ATP and GTP expanding it to balance pyrimidine and purine pools. Together, these findings uncover a dynamic breathing mechanism for long-range allosteric communication in ATCase.

## INTRODUCTION

Discovered nearly 70 years ago as a key regulatory point in pyrimidine biosynthesis (Figure 1A), *Escherichia coli* aspartate transcarbamoylase (ATCase) has served as a textbook example of allostery.^1–3^ The enzyme is composed of two catalytic trimers (Figure 1B) that are stacked with their active sites facing each other, held together by three regulatory dimers (Figure 1C) with their allosteric sites pointing outward. Each of the six active sites sequentially binds two substrates using two loops – carbamoyl phosphate (CP) via the “80s loop,” followed by L-aspartate (Asp) via the “240s loop” (Figure 1B). The condensation of these substrates initiates the pyrimidine biosynthesis pathway, which ultimately produces uridine 5′-triphosphate (UTP) and cytidine 5′-triphosphate (CTP) (Figure 1A). Following the early finding that CTP inhibits ATCase while the purine nucleotide adenosine 5′-triphosphate (ATP) stimulates it,^4^ the enzyme has captivated generations of scientists with a central question: How might the binding of pyrimidines or purines at the allosteric sites control the active sites located ∼60 Å away (Figure 1D-E, dashed paths)? Despite decades of study, this mechanism has remained the subject of intense debate.

**Figure 1.**
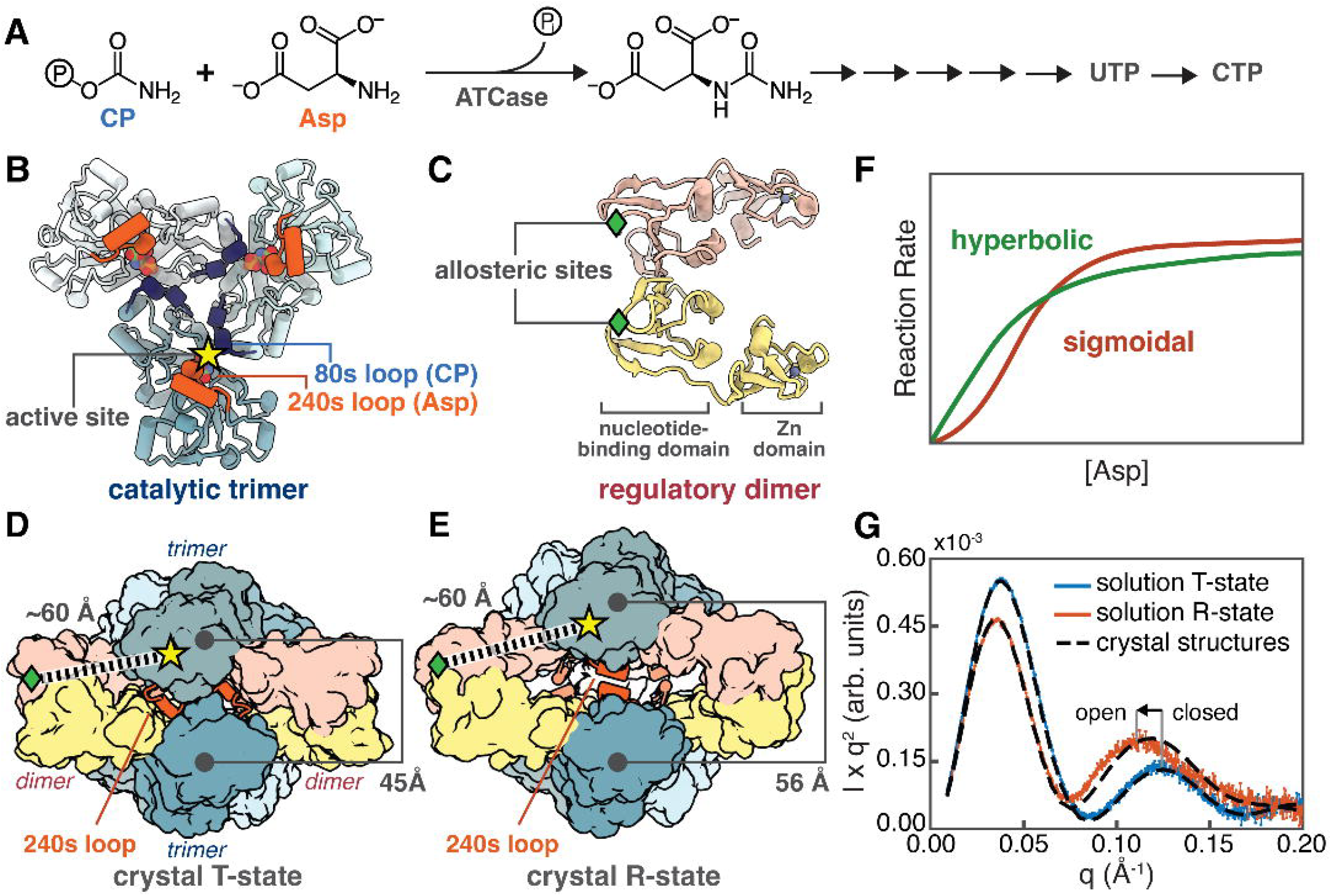
*Escherichia coli* ATCase is a complex allosteric enzyme. **(A)** ATCase catalyzes the condensation of carbamoyl phosphate (CP) and L-aspartate (Asp) in pyrimidine biosynthesis. **(B-C)** The enzyme is composed of two catalytic trimers (blue) facing each other, held together by three regulatory dimers (pink/yellow). **(B)** Each catalytic subunit houses an active site (star) that binds CP with an “80s loop” (residues 74-88, deep blue) from a neighboring subunit, then binds Asp with a “240s loop” (residues 229-247, orange) from the same subunit. **(C)** Each regulatory subunit contains a nucleotide-binding domain housing an allosteric site (diamond) and a Zn domain that contacts a catalytic subunit. Crystal structures depict two major conformations (shown are side views with one regulatory dimer in the back and not in view) with differing environments for the 240s loops: **(D)** a closed T-state and **(E)** an open R-state with center-of-mass trimer-trimer distances of ∼45 Å and ∼56 Å, respectively. Allosteric communication spans ∼60 Å. **(F)** Inhibition of ATCase produces a sigmoidal response, while activation produces a more hyperbolic curve,^4^ shown as illustrations here. **(G)** SAXS profiles shown in Kratky representation (*Iq*^2^ vs *q*) in arbitrary units (arb. units). Experimental curves are shown in color with error bars representing standard deviations in intensity *I* propagated during data reduction prior to *q*-binning; simulated curves from crystal structures are shown as dashed lines. For ATCase, the position of the second Kratky peak inversely correlates with the degree of expansion. The crystallographic T-state (PDB: 6at1)^15^ agrees with ligand-free ATCase (*blue*), whereas the crystallographic R-state (PDB: 1d09)^9^ deviates from the SAXS profile obtained with 50 µM PALA (*red*), indicating that the solution R-state is more expanded than in the crystal. The structure in panel D is 6at1, while all other structural images are 1d09.

To begin understanding this mechanism, we return to the original biochemical observations that first sparked the debate:^4^ CTP inhibited ATCase by inducing a sigmoidal dependence on the concentration of Asp, while ATP activated the enzyme by inducing a more hyperbolic dependence, as seen in conventional ligand binding (Figure 1F). Monod, Wyman, and Changeux (MWC) offered an elegant explanation for this behavior by proposing an equilibrium between two conformations with contrasting affinities for Asp: a low-affinity tense (T) state and a high-affinity relaxed (R) state.^5^ In this framework, the Asp dependence becomes more sigmoidal when (a) the intrinsic conformational equilibrium favors the T-state and (b) the affinity gap between T and R is large. Under these conditions, turnover remains low until sufficient Asp accumulates to drive a cooperative T-R transition, producing a sigmoidal relationship between activity (reaction rate) and Asp concentration. In contrast, when the enzyme exists predominantly in a single conformational state, or when the affinity difference between states is negligible, the MWC model collapses to standard Michaelis-Menten behavior with a hyperbolic dependence. Consistent with an MWC-like model, crystal structures have depicted two major conformations with distinct configurations of the Asp-binding 240s loops:^6–19^ a closed conformation consistent with a low-affinity T-state where the two trimers are held close, burying the 240s loops within subunit interfaces (Figure 1D), and an open conformation consistent with a high-affinity R-state where the trimers are separated and the 240s loops are engaged in substrate binding (Figure 1E).

However, unlike substrate binding, which induces a dramatic conformational change consistent with the MWC model, the reported effects of nucleotide binding have been far more subtle and difficult to directly relate to enzyme cooperativity.^14–16,20–25^ One contributing factor was the limitations of available structural tools. Although crystal structures revealed conformational shifts upon nucleotide binding,^14–16^ they were modest – typically sub-angstrom (±0.5 Å) – and ultimately showed no consistent pattern or correlation with enzyme cooperativity (Supplementary Fig. 1A,C). Moreover, a long-standing discrepancy between crystallography and small-angle X-ray scattering (SAXS) raised further questions.^25,26^ While crystal structures consistently showed an R-state with a fixed trimer-trimer distance of ∼56 Å (Figure 1E), SAXS data suggested that the R-state is more expanded in solution, implying that crystal packing may mask conformational changes critical for allosteric regulation.

Compounding these interpretive challenges was the complexity of the allosteric sites in ATCase. For much of the field’s history, it was assumed that only one nucleotide binds per allosteric site (Figure 1C), with CTP and ATP considered the primary effectors. Although it was known that UTP enhances the inhibitory effect of CTP,^27^ it was only more recently demonstrated that each allosteric site binds two nucleotides in the presence of the physiologically relevant counterion, Mg^2+^.^17–19^ Unfortunately, most ATCase studies predated this discovery and thus were conducted under conditions in which the allosteric sites were not fully assembled.

In this study, we leverage the capabilities of single-particle cryo-electron microscopy (cryo-EM) alongside SAXS and crystallography to overcome the limitations of each technique and capture structural changes in *E. coli* ATCase across multiple length scales. By performing structural and biochemical characterizations at physiological pH and in the presence of Mg^2+^, we obtain ten cryo-EM structures and two crystal structures that define a continuum of nucleotide-dependent conformations, validated by solution SAXS and directly correlated with activity assays. These data show that nucleotides modulate the enzyme’s cooperativity by altering the conformation of the R-state. In solution, free of crystal packing effects, the R-state behaves like a flexible balloon that expands and contracts. These motions directly impact enzyme cooperativity because the 240s loops, which are confined within the “balloon,” must rearrange in order to bind Asp. Remarkably, the expansion and contraction of the R-state are governed by symmetric nucleotide pairs. The pyrimidine pair, CTP and UTP, acts as a potent inhibitor by compressing the R-state, thereby imposing highly cooperative behavior among the 240s loops. In contrast, we find that the physiological activator is not ATP alone, as long assumed, but ATP paired with guanosine 5′-triphosphate (GTP), a nucleotide largely overlooked in prior studies. This purine pair uniquely expands ATCase to its fullest extent, allowing the 240s loops to function independently. These findings demonstrate how long-range communication between distal sites can arise from the motions of a flexible quaternary structure and reveal an elegant mechanism for balancing pyrimidine and purine pools.

## RESULTS

In addition to the absence of Mg^2+^ in many ATCase studies, previous structural and biochemical observations have not been consistently made at the same pH. Most solution-based studies, including biochemical assays and SAXS, have been performed at pH 7.0 or 8.3,^4,19,23–26,28–31^ whereas all prior nucleotide-bound crystal structures were obtained from crystals grown at mildly acidic pH (5.7–6.0).^14,15,17– 19,32^ To enable direct comparison between structural and biochemical behaviors under a physiologically relevant condition, we carried out all experiments with *E. coli* ATCase in a common buffer containing 15 mM MgCl_2_ at pH 7.5.

### The R-state is flexible in solution

We began by using SAXS to reexamine the R-state of ATCase in solution at a dodecamer concentration of 8 μM (Supplementary Table 1). To prevent turnover during measurements, we first used *N*-(phosphonoacetyl)-L-aspartate (PALA), a tight-binding bisubstrate analog known to trigger the same quaternary response as native substrates,^33^ to ensure that the transition from the T-state to R-state is complete. It is well known that ATCase scattering displays a prominent secondary peak in a Kratky representation of the SAXS data (*Iq*^2^ vs *q*) that is highly sensitive to changes in quaternary structure.^26^ Consistent with previous observations made at pH 7.0 and 8.3,^23,24,26,28^ we observed a left-shift of this peak (Figure 1G, arrow), which, in combination with singular value decomposition (SVD) of the dataset (Supplementary Fig. 2A), confirmed that PALA induces a two-state transition from a closed to an open conformation. We then verified that the same transition is induced by a more physiological pair of ligands: the native substrate CP paired with succinate, a commonly used analog of Asp^12,23^ (Supplementary Fig. 2B-C). While the closed state observed in solution matches the T-state crystal structure, the secondary Kratky peak for the open state is further left-shifted than that of the R-state crystal structure (Figure 1G), indicating that the solution R-state is more expanded and sufficiently flexible to be compressed within the crystal lattice.

Given the apparent flexibility of the R-state in solution, we next examined whether nucleotides induce structural changes. We prepared the solution R-state using saturating concentrations of CP (500 μM) and succinate (10 mM) and obtained SAXS profiles at varying nucleotide concentrations (Supplementary Tables 2–3): (a) 0–3 mM CTP, (b) 0–5 mM CTP + UTP, (c) 0–4 mM ATP, and (d) 5 mM ATP + 0–3 mM

GTP. Compared to the dramatic changes observed with PALA or CP/succinate, nucleotide-induced changes in scattering were relatively subtle (Supplementary Fig. 3), suggesting that nucleotides do not merely shift the T–R equilibrium. Indeed, SVD analysis of each titration dataset indicated that nucleotides stabilize new conformations distinct from both the T-state and the nucleotide-free R-state (Supplementary Fig. 3).

To confirm these subtle effects, we performed size-exclusion chromatography-coupled SAXS (SEC-SAXS) under conditions representing the titration endpoints (Supplementary Tables 4-6) and extracted high-quality scattering profiles via model-free deconvolution^34^ (Figure 2A, Supplementary Fig. 4-6). These profiles recapitulate the trends observed in the titrations. While the ligand-free T-state displays a Kratky peak at 0.122 Å^-1^, a gradient of peak positions is observed for the R-states (Figures 2A, Supplementary Fig. 7). At a saturating concentration of 1.5 mM, the canonical inhibitor CTP induced a small right-shift of the Kratky peak without reaching the T-state position, indicating compaction of the R-state. Conversely, the canonical activator ATP (5 mM) induced a slight leftward shift, consistent with expansion of the R-state. Notably, these effects were amplified when CTP was combined with UTP (1.5 mM each) and when ATP was paired with GTP (5 mM ATP + 1 mM GTP). Altogether, SAXS experiments confirmed that nucleotides modulate the conformation of the R-state, with the pyrimidine and purine pairs, CTP/UTP and ATP/GTP, exerting the strongest effects.

**Figure 2.**
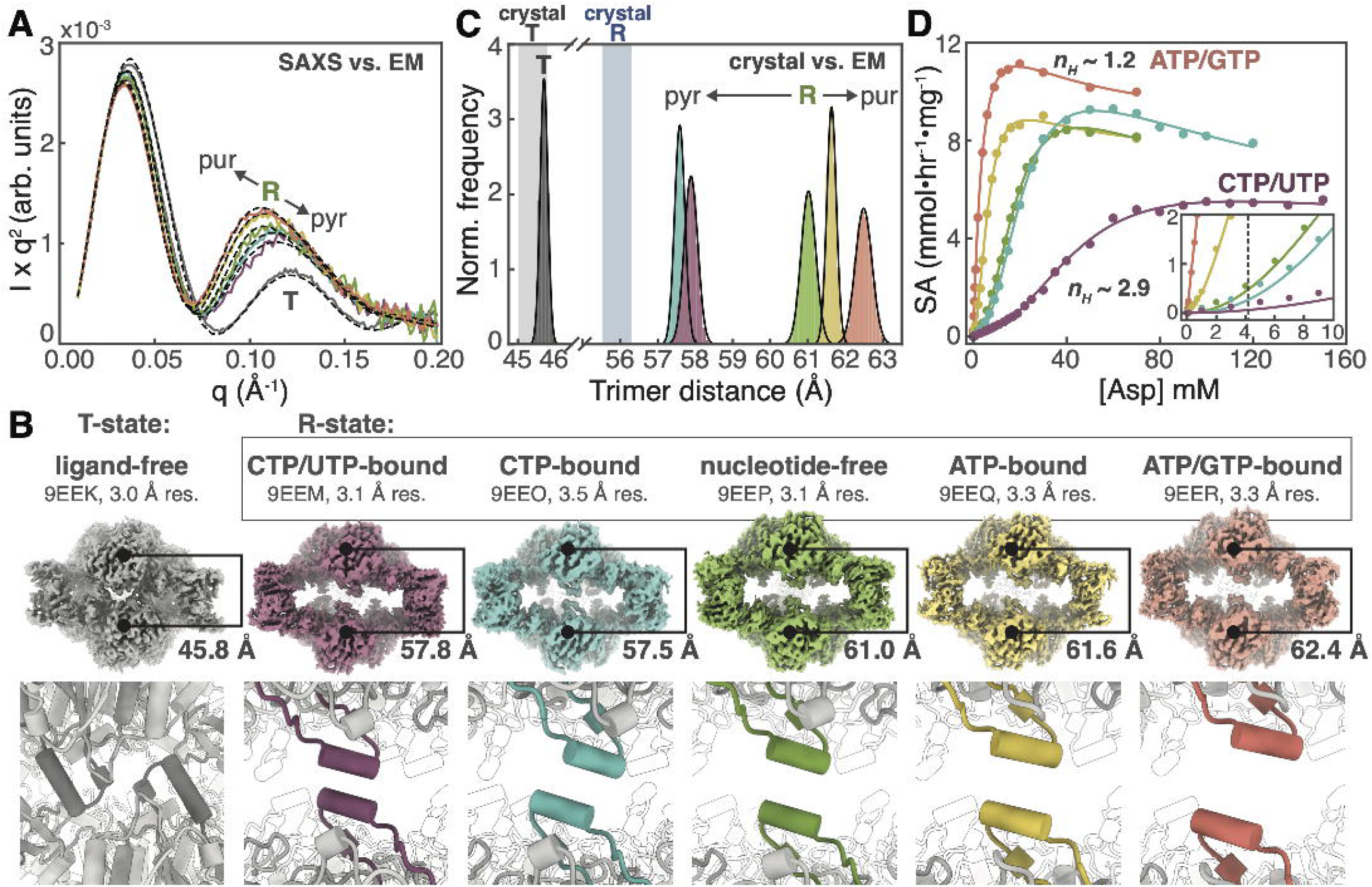
Nucleotides exert control over cooperativity by expanding or contracting the R-state, which behaves like a flexible balloon. SAXS, cryo-EM, and biochemical assays were performed on *E. coli* ATCase in the same buffer (pH 7.5, 15 mM MgCl_2_). The T-state condition (*gray*) contained no ligands. R-state conditions included 500 μM CP, 10 mM succinate, and one of the following nucleotide combinations: 1.5 mM CTP/UTP (*purple*), 1.5 mM CTP (*blue*), no nucleotides (*green*), 5 mM ATP (*yellow*), or 5 mM ATP + 1 mM GTP (*salmon*). **(A)** SEC-SAXS profiles (solid curves) show peak shifts consistent with a flexible R-state that expands in the presence of purines and contracts with pyrimidines. Profiles align well with cryo-EM models shown in panel B (dashed curves). **(B)** Cryo-EM 3D classification resolved only T-state or R-state particles (no intermediates), but nucleotides modulate the openness of the R-state. *Top:* Consensus maps at 3.0–3.5 Å resolution. *Bottom:* Refined models illustrate how nucleotides modulate the distance between catalytic trimers, visualized by the separation of the 240s loops within the central cavity. **(C)** 3D Variability Analysis (3DVA) reveals that in solution, the R-state is more expanded than in crystal structures. Purine- and pyrimidine-bound forms differ by up to ∼5 Å in center-of-mass distance between trimers (mean values indicated in panel B). Shaded bars indicate distances from crystal structures: T-state (45.0–45.8 Å) and R-state (55.5–56.3 Å). **(D)** Specific activity (SA) of ATCase measured in replicates (model fits shown in Supplementary Fig. 18) under the same nucleotide conditions as panels A–C, with 4.8 mM CP and varying [Asp]. CTP/UTP promotes strong cooperativity, as reflected in the Hill coefficient *n*_h_, while ATP/GTP removes this effect. At high [Asp], activities decrease, resembling substrate inhibition.^31^ *Inset:* Activity curves near physiological [Asp] of 4.2 mM.

### Cryo-EM reveals a gradient of R-states

To visualize nucleotide-induced changes at high resolution, we next performed cryo-EM of 2 μM *E. coli* ATCase under the same six conditions used for SEC-SAXS (Supplementary 8-15, Supplementary Tables 7-8). From these conditions, 3D classification and subsequent refinement yielded three T-state and five R-state consensus maps at overall resolutions of 3.0-3.5 Å, as estimated by gold-standard Fourier shell correlation (FSC) at the 0.143 threshold. The enzyme interior, including the active sites, generally aligned to higher resolution (∼2.5-3.0 Å) than the peripherally located allosteric sites (∼2.8-4.5 Å). For all nucleotide-containing conditions, the resulting cryo-EM maps supported the presence of two nucleotides per allosteric site, coordinating a shared Mg^2+^ ion (Supplementary Fig. 16).

For ligand-free ATCase, a single consensus class was obtained corresponding to a closed T-state, and the resulting model agreed well with the scattering profile from SEC-SAXS (Figure 2A-B, gray). In contrast, conditions containing saturating CP and succinate produced a gradient of R-state maps, with pyrimidines or purines inducing compaction or expansion, respectively (Figure 2B). These changes are readily visualized in the refined R-state models, where the 240s loops come nearly into contact in the presence of pyrimidines but are far apart with purines (Figure 2B, bottom row). Importantly, these models agree well with SAXS data under matching conditions (Figure 2A), supporting the interpretation that in solution, the R-state conformation is inherently flexible and modulated by nucleotide binding.

To assess this flexibility, particles used for each consensus refinement were subjected to 3D variability analysis (3DVA)^35^ (Supplementary Figs. 8-13, Supplementary Methods). The conformational variability in each state could be described by a single, breathing-like mode along the axis connecting the catalytic trimers (Supplementary Movies 1-6). By fitting rigid-body models into sub-volumes along the 3DVA trajectory, we generated particle distributions as a function of the center-of-mass distance between catalytic trimers (Figure 2C). The sharpness of the T-state distribution and its overlap with trimer distances from crystal structures suggest that the T-state is relatively rigid (Figure 2C, gray peak vs. gray bar). In contrast, substantial differences were observed for the R-states (Figure 2C, colored peaks vs. blue bar): cryo-EM models were more expanded than crystal structures, and pyrimidine- and purine-bound forms differed by as much as ∼5 Å – an order of magnitude greater than the sub-angstrom shifts observed crystallographically^14–16^ (Supplementary Fig. 1). Together, these findings suggest that the R-state behaves like a flexible balloon – compressed in crystal lattices by packing interactions, and in solution, compressed by pyrimidines or expanded by purines, reaching maximum expansion with the ATP/GTP pair.

Cryo-EM also revealed that nucleotide binding can shift the T-R equilibrium. In datasets containing CP, succinate, and pyrimidines (CTP alone or CTP/UTP), 3D classification identified minor T-state populations (Supplementary Figs. 9-10). The key distinction between refined R- and T-state maps was the presence or absence of the second substrate: R-state maps showed density for both CP and succinate, whereas T-state maps lacked succinate density (Supplementary Fig. 17), consistent with the 240s loops being sequestered in the T-state conformation. Fitting linear combinations of T- and R-state cryo-EM models to SAXS data indicated that cryo-EM can overestimate the T-state population (Supplementary Fig. 17), possibly due to perturbation by the thin-layer sample environment. Nonetheless, the SAXS profile for the CTP/UTP condition was best fit with a 20:80 mixture of the T- and R-states (Supplementary Fig. 17A). The appearance of a T-state population only in cryo-EM datasets with pyrimidines and in SAXS data for the CTP/UTP condition further refine our model: purine-induced expansion of the R-state likely stabilizes it by resisting collapse into the T-state, whereas pyrimidine-induced compression likely destabilizes it, revealing a T-state population under certain conditions.

### Cooperativity tracks with R-state compression

To directly compare structural changes with biochemical behavior, we performed activity assays of 1 nM ATCase at the same nucleotide concentrations used for SEC-SAXS and cryo-EM (Figure 2D). Initial rates were measured at varying Asp concentrations with CP fixed at 4.8 mM. Consistent with prior observations at pH 7.0 and 8.3,^19^ we find that with Mg^2+^, the combination of CTP and UTP is most inhibitory, while the combination of ATP and GTP is most activating at physiological Asp concentrations of ∼4.2 mM^36^ (Figure 2D, inset).

The CTP/UTP pair produced a highly sigmoidal saturation curve, while the ATP/GTP pair produced a near-hyperbolic curve at low [Asp] (at high [Asp], ATCase is known to exhibit behavior reminiscent of substrate inhibition^31^). Fits to the Hill equation in the transition regions (Supplementary Fig. 18, Supplementary Table 9) revealed that the Hill coefficient *n*_*H*_ and the Asp concentration needed to reach half-maximal activity *K*_*1/2*_ are inversely correlated with the openness of the R-state observed by cryo-EM (Supplementary Fig. 1D). Two important conclusions can be drawn: (1) CTP and UTP together inhibit ATCase by compressing the R-state to produce highly cooperative behavior (*n*_*H*_ ∼2.9), rendering the enzyme virtually inactive near physiological [Asp] (*K*_*1/2*_ ∼37 mM), whereas (2) ATP and GTP together produce the most expanded R-state, where cooperative behavior is largely lost (*n*_*H*_ ∼1.2) and the enzyme is highly active at physiological [Asp] (*K*_*1/2*_ ∼ 2.9 mM). Since cellular concentrations of the four nucleotides^36,37^ exceed those needed to saturate the structural effects induced by these mixed-nucleotide pairs (Supplementary Table 10), it is likely that these binding modes are more physiologically relevant than binding to only one type of nucleotide or none at all.

### ATCase is specific for CTP/UTP and ATP/GTP

SAXS, cryo-EM, and biochemical assays collectively indicated that the pyrimidine pair CTP/UTP and the purine pair ATP/GTP exert the clearest effects on ATCase structure and activity. To determine whether the enzyme preferentially binds these pairs, we turned to crystallography to obtain high-resolution insight into nucleotide specificity (Supplementary Table 11).

Prior crystallographic studies showed that when CTP binds the canonical nucleotide-binding site (“site 1” in Figure 3A), a Mg^2+^ ion enables UTP to bind in an adjacent site (“site 2” in Figure 3A), recruiting the flexible N-terminus of the regulatory subunit.^18,19^ We confirmed this result in our own soaking experiments with CTP and brominated UTP.^38^ Prior studies also reported that in the presence of Mg^2+^ and either CTP or ATP, electron density is consistent with two CTP or two ATP molecules per allosteric site,^18,19^ but weak site-2 density suggested that these same-nucleotide pairings are suboptimal. Because ATP was introduced by soaking in the previous study, we solved a 3.0-Å co-crystal structure of ATCase bound to ATP, CP, and succinate at near-neutral pH (Supplementary Figs. 19-20). Weak density was observed for the second-site ATP as well as for the N-termini, which reach across the dimer interface without adopting an ordered conformation (Figures 3D, Supplementary Fig. 20).

**Figure 3.**
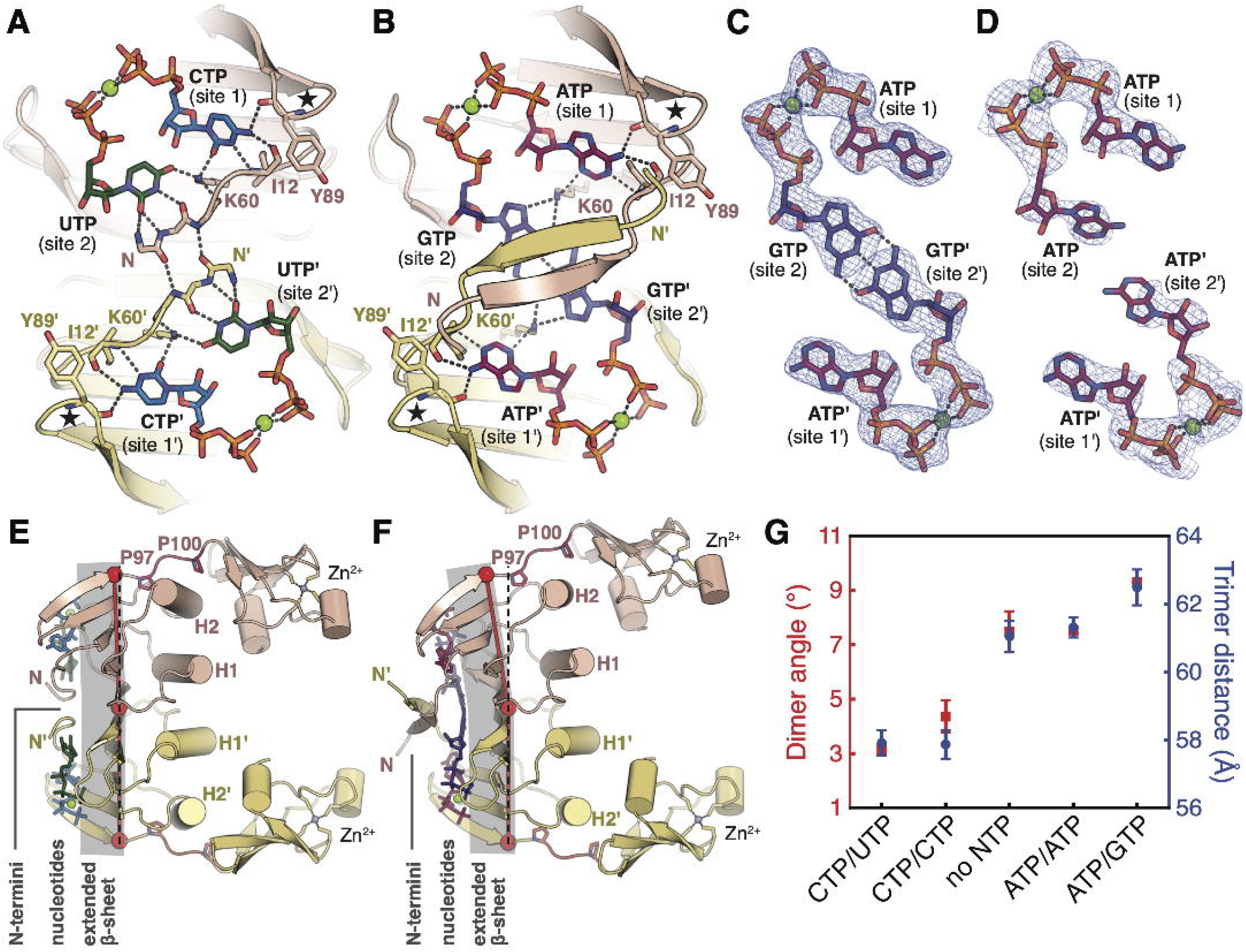
Symmetric pairs of pyrimidines and purines bind specifically to *E. coli* ATCase, inducing structural changes that couple the allosteric and active sites. **(A)** Crystal structure of CTP/UTP-bound R-state^19^ (pdb: 4kh1) shows Y89 and I12 favoring CTP in the first site, while N-terminal residues favor UTP in the second. K60 provides selectivity for both. **(B)** 2.2-Å crystal structure of ATP/GTP-bound R-state at neutral pH (this study). ATP binds preferentially in the first site; K60 selects for GTP in the second. GTPs form a non-canonical base pair, stabilized by K60 and T43 (Supplementary Fig. 22) and a hydrophobic pocket formed by a domain-swapped N-terminal β-sheet. In panels A-B, the outer loop (residues 88-89) is starred, and Mg^2+^ ions are green spheres. **(C)** Polder omit map (*blue mesh*, 6σ) supports nucleotide assignments in panel B. **(D)** 3.0-Å crystal structure of ATP-bound R-state (this study) at near-neutral pH. Polder omit map (*blue mesh*, 6σ) shows disordered ATP in the second site. **(E)** Side view of the regulatory dimer in the cryo-EM CTP/UTP R-state model shows a planar β-sheet (shaded) holding the Zn-domains close. The H2-helix remains angled (as in T-state structures), while H2’ straightens. **(F)** Corresponding view of the ATP/GTP R-state model shows a bent β-sheet (shaded) increasing the Zn-domain separation, with both H2 and H2’ straightening and becoming parallel. In panels E-F, linker residues P97 and P100 are shown as sticks, and the dimer angle is defined by three points on the β-sheet (*red spheres*) and its deviation from planarity (*black dashed*). **(G)** Bending of the regulatory dimer modulates Zn-domain separation and correlates with the center-of-mass distance between catalytic trimers. The dimer angle is 180 ° minus the average of the three regulatory dimers, with error bars representing the standard deviation; trimer distances are the mean with error bars represented as the full-width-at-half-maximum values of the trimer distance distributions from 3DVA in Figure 2C.

We then obtained a previously unreported *P*321 crystal form at pH 7.0-7.5 (Supplementary Fig. 21, Supplementary Table 11), yielding a 2.19-Å resolution structure of ATCase bound to ATP, GTP, and PALA. Strong density supported the assignment of ATP in site 1 and GTP in site 2, coordinating a shared Mg^2+^ ion (Figures 3B-C, Supplementary Fig. 22). Unlike when ATP occupies site 2 (Figure 3D), here, we observe well-ordered GTPs forming a non-canonical base pair across the dimer interface (Figure 3C). For such an interaction, one GTP must be deprotonated at the N1 position at any given time. Although the pKa at this position is 9.2 in free GTP, electronic structure calculations suggest that it can be shifted towards neutrality by a positive charge on the Hoogsteen face of the nucleobase,^39^ which is provided by a conserved^40^ K60 in ATCase (Supplementary Fig. 23). Such a mechanism would explain why a GTP-bound structure had not been obtained under previous acidic conditions as well as why our crystals dissolve at lower pH. Although this crystal structure featured a compressed trimer-trimer distance of 56.2 Å, the same ATP/GTP crystallization condition also produced a second crystal form that diffracts to lower resolution but is elongated by nearly 14 Å along the **c**-axis, which lies parallel to the direction of the quaternary expansion observed in solution by cryo-EM and SAXS (Supplementary Fig. 21C, Supplementary Table 12).

Although resolution at the allosteric sites in our cryo-EM maps is limited by the inherent flexibility of ATCase, the same trend is observed: site-2 density is weaker, and the N-termini are disordered for same-nucleotide pairings (Supplementary Fig. 16). Similarly, our soaking experiments suggested that mixed pyrimidine-purine pairings are also suboptimal.^38^ Altogether, observations from crystallography and cryo-EM indicate that the physiological binding partner for CTP is indeed UTP, and for ATP, it is GTP.

Structural comparison revealed how these nucleotide pairs rearrange the regulatory subunits. Site 1 is specific for the amino group and the adjacent nitrogen of CTP or ATP, which hydrogen-bond with the backbone carbonyls of I12 and Y89 and with the backbone amide of I12 or A11 (Supplementary Fig. 24A). These interactions couple the N-terminus (residues 1-14) with the “outer loop” (residues 88-89, stars in Figure 3A-B). K60 bridges sites 1 and 2, favoring interactions with the carbonyl oxygen of a UTP or GTP in site 2. When CTP occupies site 1, the N-terminus wraps around UTP in site 2, forming additional base-specific interactions that bring the two N-termini together, creating a barrier between the CTP/UTP pairs across the dimer (Figure 3A). Conversely, when ATP occupies site 1, K60 selects for GTP in site 2, stabilizing non-canonical base-pairing across the dimer interface to form a 4-nucleotide unit (Figure 3B-C). Here, the N-termini form a domain-swapped antiparallel β-sheet above the plane of the nucleotides, making a hydrophobic pocket for the GTP bases (Supplementary Fig. 24B-C). Altogether, these features explain the critical role of the N-termini in allosteric regulation^14,41,42^ and the specificity for the pyrimidine pair CTP/UTP and the purine pair ATP/GTP.

### Nucleotides trigger local-to-global changes

We next asked how nucleotide binding at the allosteric sites promotes changes in internal structure that leads to global changes in quaternary expansion. To do so in an unbiased manner, cryo-EM models were refined using non-crystallographic symmetry (NCS) restraints – a strategy adapted from crystallography that detects deviations in symmetry and thereby reveals structural rearrangements (Supplementary Methods, Supplementary Fig. 25).

The refined cryo-EM models reveal a clear nucleotide-dependent path for allosteric communication between the nucleotide-binding site to the active site (Figure 3E-G). Each regulatory dimer forms an extended β-sheet across the dimer interface. The outer face forms polar interactions with nucleotides, while the inner face, which consists almost entirely of isoleucines, makes hydrophobic interactions with two pairs of helices, H1 and H2 (Figure 3E-F). The H2-helices directly pack against the Zn domains, which are responsible for binding catalytic subunits. Remarkably, we find that the nucleotide binding controls the local bending of the regulatory dimers, which directly correlates with the openness of the R-state quaternary structure (Figure 3G).

Binding of ATP/GTP induces the largest bending of the β-sheet (9.3° in Figure 3F), pulling up the outer loops that connect to the Zn domain linkers. This subtle bending of the regulatory dimer is amplified into a substantial separation between the catalytic trimers. Interestingly, the linker residues P97 and P100 form a groove around the H2-helices, which repack in response to dimer bending. In the T-state, the H2-helices are tucked into the “armpits” of the regulatory dimer, angled in opposing directions (Supplementary Fig. 25D). In contrast, in the ATP/GTP-bound R-state, the dimer opens, allowing the H2-helices to slide into a parallel, more relaxed configuration (Figure 3F, Supplementary Fig. 25A).

By comparison, CTP/UTP binding produces a nearly flat β-sheet (3.1° in Figure 3E), resulting in an intermediate Zn-domain separation that is larger than that in the T-state but smaller than that of the ATP/GTP-bound R-state. This partial compaction introduces asymmetry within each regulatory dimer: one H2-helix is angled, as in the T-state (H2 in Figures 3E, Supplementary Fig. 25B) while the other straightens (H2’ in Figure 3E, Supplementary Fig. 25B). Interestingly, this unusual asymmetry is also observed in R-state crystal structures (Supplementary Fig. 25C),^14^ which are more compressed than any of our cryo-EM models (Figure 2C).Thus, we find that this asymmetry is only a feature of collapsed forms of the R-state, which is induced in solution by pyrimidines but not purines.

### ATCase bypasses the T-state with ATP/GTP

Given that binding of the purine pair ATP/GTP leads to an unusually expanded quaternary structure, one not previously visualized, we next asked whether this expansion is driven by the nucleotides or by the substrates. Using SAXS, we first examined ATCase at 5 mM ATP and 1 mM GTP, as in earlier sections except without any substrates. Much to our surprise, we observed a substantial left-shift of the second Kratky peak relative to that of the T-state (Figure 4A-B, green), indicating that ATP/GTP alone can expand the enzyme. When CP was added to this condition, the peak shifted all the way to the position observed with both CP and succinate (Figure 4A-B, orange/red). No other nucleotide combination induced a significant structural change in the absence of substrates or with CP alone (Figure 4A-B).

**Figure 4.**
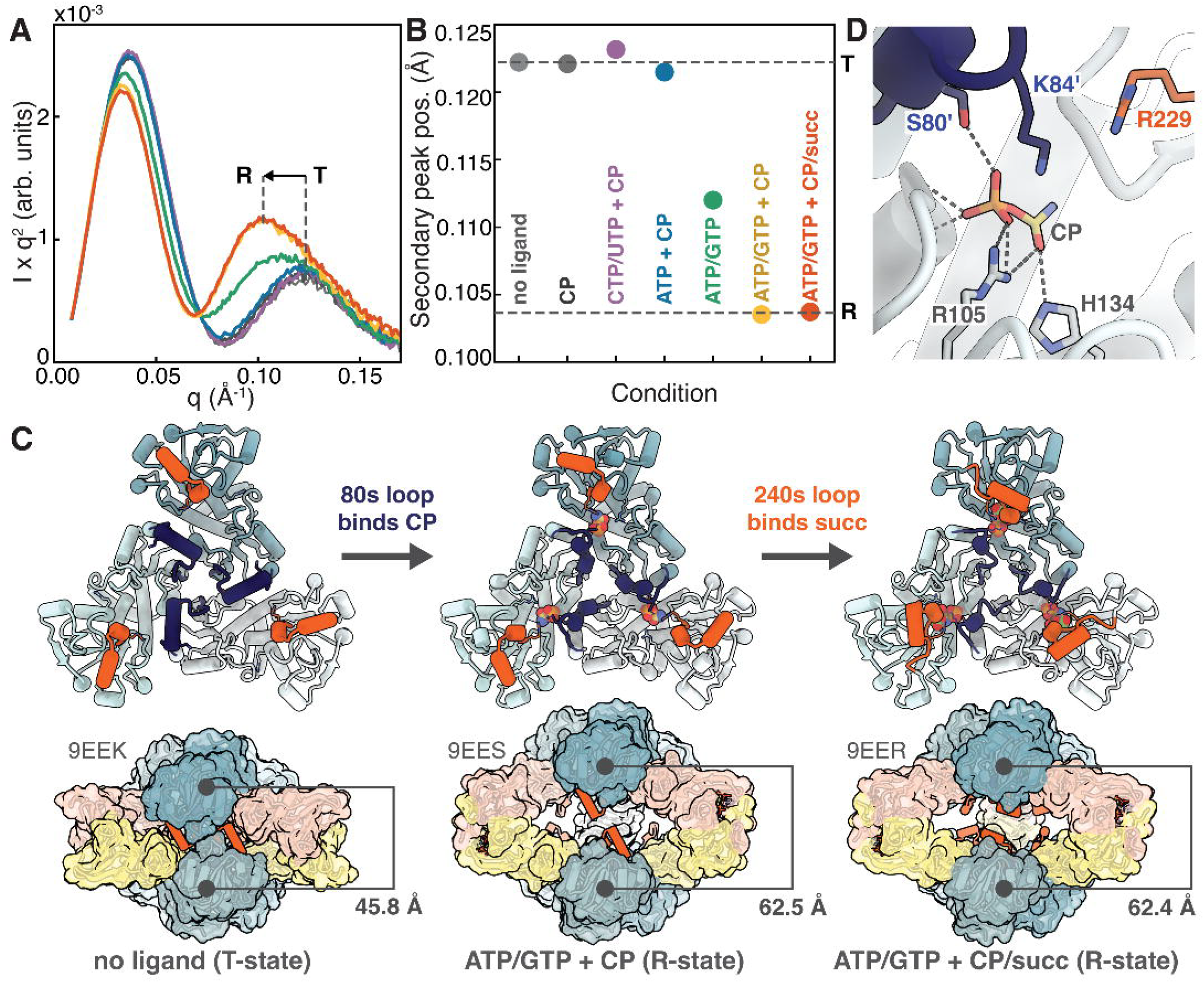
Cooperative behavior is lost with the purine pair, ATP/ GTP, because ATCase bypasses the T-state conformation. **(A)** SAXS profiles of *E. coli* ATCase, shown as a Kratky plot, and **(B)** corresponding positions of secondary peak in the same color scheme. In the absence of ligands, ATCase adopts the T-state (*light gray*); addition of the first substrate alone (500 μM CP) causes little change (*dark gray*), even in the presence of 1.5 mM CTP/UTP (*purple*) or 5 mM ATP (*blue*). Unlike all other nucleotide combinations, the purine pair (5 mM ATP, 1 mM GTP) induces a conformational shift even without substrates (*green*); addition of 500 μM CP completes the transition to the R-state (*yellow*-*orange*), and further expansion is not observed with 500 μM CP, 10 mM succinate (*red*). **(C)** Cryo-EM model of ATCase obtained with 5 mM ATP, 1 mM GTP, 500 μM CP (*middle*) reveals a distinct conformation, where the 80s loops (*deep blue*) engage CP but the 240s loops (*orange*) remain extended. Ligand-free T-state (*left*) and the ATP/GTP R-state (*right*) from Figure 2 shown for comparison. **(D)** Close up of the active site in the distinct conformation shows CP interacting with S80 and K84 of the neighboring 80s loop (*deep blue*). Engagement of the 240s loop (*orange*) with the second substrate involves interactions with both R229 and R234, but only R229 is observed near the empty site.

To confirm this unexpected expansion, we performed cryo-EM on ATCase with just CP, ATP, and GTP (Supplementary Fig. 26, Supplementary Table 7). 3D classification and subsequent refinement resulted in a ∼3.2-Å map of a fully open conformation with an inter–trimer distance of 62.5 Å that fully explained the SAXS profile obtained under the same condition (Supplementary Fig. 27). A defining feature of this distinct conformation is its active-site configuration: the 80s loops are bound to CP (Figure 4D), but the 240s loops extend freely into the central enzyme cavity, remaining available to bind Asp (Figure 4C, middle). Thus, this conformation represents a “pre-Asp conformer” within the R-state ensemble. Notably, when we docked the catalytic subunits from this model into cryo-EM maps obtained under other nucleotide conditions, we observe steric clashes between opposing 240s loops (Supplementary Fig. 28). These observations provide a structural explanation for why cooperative behavior is lost with the purine pair ATP/GTP: once CP binds, ATCase bypasses the T-state entirely, achieving an R-state conformation that is sufficiently expanded that the 240s loops are able to move independently between their “pre-Asp” and “Asp-bound” configurations (Figure 4C).

## Discussion

For nearly 70 years, the mechanism of nucleotide-dependent allosteric regulation in *E. coli* ATCase has been challenging to resolve. A central question has been why the magnitude of Asp-binding cooperativity changes so dramatically in the presence of different nucleotides. In the classical MWC model, such changes arise because allosteric ligands shift the intrinsic T-R equilibrium constant. However, this model implies that ATCase interconverts between only two discrete conformations. As we show in this study, the R-state is highly flexible in solution, and thus, all prior crystal structures of the R-state have captured it in an artificially compressed form. In the crystal lattice, this compression is physically limited by the 240s loops coming into contact (acting like “bumpers”), resulting in R-state crystal structures with a nearly invariant trimer-trimer distance of ∼56 Å (Supplementary Fig. 1A,C). Yet, even within the crystallography field, it has long been recognized that nucleotides must induce structural perturbations,^14,16^ but their magnitude and significance remained unclear due to discrepancies across different experimental techniques^25,26^ and because many earlier studies did not use conditions that support proper assembly of the allosteric sites (e.g., Mg^2+^).^17–19^

By integrating insights from cryo-EM, SAXS, and crystallography, we show that the R-state is not a single conformation but a flexible structure that responds dynamically to allosteric effectors by expanding or contracting (Supplementary Fig. 1B,D). Given the estimated intracellular concentrations of nucleotides in *E. coli* (Supplementary Table 10),^36^ a nucleotide-free form of ATCase is likely rare. Furthermore, although we found that the allosteric site can bind two molecules of CTP or ATP (Supplementary Fig. 16), our data indicate that the most physiologically relevant effectors are the CTP/UTP and ATP/GTP pairs, which exhibit specific binding and elicit the strongest regulatory effects (Figures 2D,3). The pyrimidines CTP and UTP together suppress ATCase activity at physiological Asp concentrations by producing a strongly sigmoidal dependence on Asp (Figure 2D), whereas the purines ATP and GTP together produce a hyperbolic Asp dependence, reaching a substantially higher activity at low Asp concentrations (Figure 2D).

Based on these findings, we propose a dynamic model of how nucleotides tune the cooperativity of *E. coli* ATCase (Figure 5). Unlike the classical MWC model – in which nucleotides simply shift the intrinsic equilibrium between fixed T- and R-state conformations – our results show that nucleotide binding reshapes the conformational landscape by generating a continuum of R-state conformations (R′, R″, …). Substrate binding, particularly by Asp or its analogs, induces a direct rearrangement of the active-site 240s loops, whereas nucleotides bind at peripheral sites to exert an indirect effect on the 240s loops by compressing or expanding the enzyme. The pyrimidine pair CTP/UTP induces a partially collapsed, asymmetric form of the regulatory dimers, resulting in a compressed R-state when substrates are bound. The tendency of this pyrimidine pair to compress the enzyme assures that the closed T-state is favored in the absence of substrates. In the T-state, the CP-binding 80s loops retain mobility, whereas the 240s loops are buried within subunit interfaces and must be released to allow Asp binding (Figure 1D). Because the CTP/UTP-bound R-state confines the 240s loops within a reduced space, high concentrations of Asp are needed to drive the T-to-R transition, accounting for the observed high cooperativity. In contrast, the ATP/GTP pair is unique among all nucleotide combinations in expanding the enzyme to such a degree that the T-state becomes disfavored, and CP binding alone is sufficient to complete the transition to the R-state. Under these conditions, cooperativity is lost because a T-state conformation with low Asp affinity is bypassed. Structurally, this loss of cooperativity can also be explained by the expanded trimer-trimer distance, which allows the 240s loops to move independently without steric interference. Overall, nucleotides regulate Asp cooperativity through two coupled mechanisms: (a) by shifting the relative stabilities of closed and open conformations and (b) by reshaping the conformational landscape itself through changes in the R-state structure.

**Figure 5.**
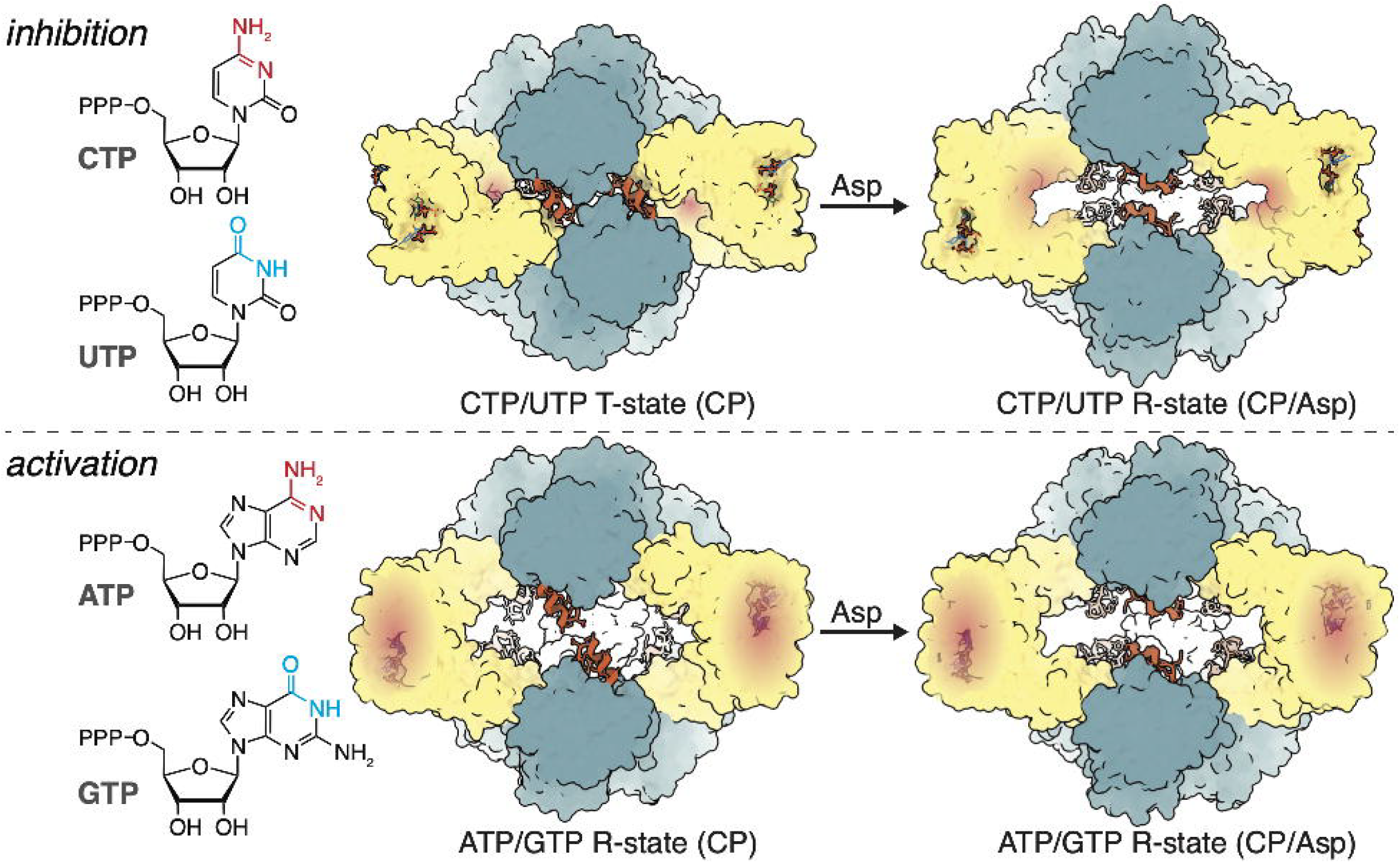
Model for nucleotide-dependent allosteric regulation of *E. coli* ATCase. ATCase is a flexible enzyme whose conformational changes are governed by symmetric nucleotide pairs: CTP/UTP for inhibition and ATP/GTP for activation. Each allosteric site achieves specificity for these pairs by recognizing features of CTP and ATP nucleobases (*red*) in one position, and those of UTP and GTP (*cyan*) in the adjacent site. Affinity for the second substrate, Asp, is determined by the ability of the 240s loop (*orange*) to move within the confined space of the enzyme cavity. *Top:* With CTP/UTP bound, the regulatory dimers (*yellow*) are pinched closed, holding the catalytic trimers (*blue*) close together and imposing cooperativity among the 240s loops in binding Asp. *Bottom:* With ATP/GTP bound, the regulatory dimers open, allowing the enzyme to bypass the T-state. In this expanded conformation, the enzyme behaves like dissociated catalytic trimers, removing cooperativity among active sites by allowing independent movement of the 240s loops.

Intriguingly, not all members of the ATCase family possess regulatory subunits.^43^ Like these ATCases which naturally exist as catalytic trimers,^44^ denaturation studies of *E. coli* ATCase have shown that removal of regulatory subunits leads to high activity and no cooperativity.^4^ Thus, the loss of cooperativity observed with the purine pair ATP/GTP can also be understood as the result of inducing a conformation in which the catalytic trimers and regulatory dimers are nearly dissociated. These observations suggest that the regulatory subunit evolved to enable switching between two functional modes: one in which ATCase is subject to feedback inhibition by end products of the pyrimidine biosynthesis pathway, and another in which it behaves like a constitutively active catalytic trimer in response to elevated levels of end products from the competing pathway. Altogether, our results highlight how the intrinsic flexibility of the R-state enables ATCase to respond to opposing metabolic signals.

## METHODS

### Protein Expression and Purification

Tagless *E. coli* ATCase (*pyrBI*) was expressed in T7 Express BL21 (DE3) *E. coli* cells (New England Biolabs) and purified following previous protocols^18^ with modifications detailed in the Supplementary Methods. In the final size-exclusion step, the protein was exchanged into ATCase buffer (40 mM Tris-HCl pH 7.5, 15 mM MgCl_2_, 1 mM TCEP). Protein concentrations throughout are given in molar concentrations of the holoenzyme (308.4 kDa dodecamer).

### Activity Assays

Enzyme kinetics were measured using a colorimetric assay for carbamoyl aspartate production.^45^ Reaction mixtures contained 1 nM ATCase, 40 mM Tris-HCl pH 7.5, 15 mM MgCl_2_, 4.8 mM CP, 0–150 mM Asp, and nucleotide(s). The assays and kinetic modeling are detailed in the Supplementary Methods.

### SAXS

SAXS was performed at the Cornell High-Energy Synchrotron Source (CHESS) ID7A station as detailed in the Supplementary Methods. Integrated scattering profiles were normalized by transmitted intensities, and scattering from a matched buffer was used to yield the background-subtracted protein scattering profile, *I*(*q*), where *q*= 4π/*λ* sin *θ, λ* is the X-ray wavelength, and 2*θ* is the scattering angle. Titrations were performed with 8 μM ATCase in ATCase buffer with variable ligand concentrations (Supplementary Tables 1-3). SEC-SAXS was performed with 56 μM ATCase in ATCase buffer with six different ligand conditions (Supplementary Tables 4-6): 1) no ligands, 2) 500 μM CP, 10 mM succinate, 3) 500 μM CP, 10 mM succinate + 1.5 mM CTP, 4) 500 μM CP, 10 mM succinate + 1.5 mM CTP, 1.5 mM UTP, 5) 500 μM CP, 10 mM succinate + 5 mM ATP, and 6) 500 μM CP, 10 mM succinate + 5 mM ATP, 1 mM GTP.

### Cryo-EM

Samples for cryo-EM contained 2 μM ATCase in ATCase buffer with seven different ligand conditions (Supplementary Tables 7-8): 1) no ligands, 2) 500 μM CP, 10 mM succinate, 3) 500 μM CP, 10 mM succinate + 1.5 mM CTP, 4) 500 μM CP, 10 mM succinate + 1.5 mM CTP, 1.5 mM UTP, 5) 500 μM CP, 10 mM succinate + 5 mM ATP, 6) 500 μM CP, 10 mM succinate + 5 mM ATP, 1 mM GTP, and 7) 500 μM CP + 5 mM ATP, 1 mM GTP. Data were collected on a Talos Arctica (Thermo Fisher) at the Cornell Center for Materials Research (CCMR), operating at 200 keV with a Gatan K3 detector, and on a Krios G4 (Thermo Fisher) at the National Center for CryoEM Access and Training (NCCAT) operating at 300 keV with a Gatan K3 detector. All steps from sample preparation to data processing in cryoSPARC 4^46^ and RELION 4^47^ (Supplementary Figs. 8-15, 26) are detailed in the Supplementary Methods. Cryo-EM maps are rendered in ChimeraX.^48^

### Crystallography

Co-crystals of ATCase with PALA, ATP, and GTP were obtained via vapor diffusion with a 1:1:2 mixture of precipitant (8% (v/v) Tacsimate pH 7.0 and 9-14% (w/v) PEG 3350), seed solution, and protein solution (16 μM ATCase in ATCase buffer with 10 mM ATP, 2 mM GTP, 2 mM PALA). Room-temperature diffraction was collected at the CHESS ID7B2 station on an Eiger2 16M detector with an X-ray wavelength of 0.9185 Å, 10-ms exposures, and 0.1° oscillation steps.

Co-crystals of ATCase with CP, succinate, and ATP were obtained via vapor diffusion using a 1:1 mixture of protein solution (16 μM ATCase in 40 mM Tris-HCl pH 7.0, 20 mM CP, 20 mM succinate, 30 mM MgCl_2_, and 10 mM ATP) and precipitant (20-25% (w/v) PEG 300 and 0.1 M Bis-Tris pH 6.5). Cryo-protectant solutions matched the crystallization drops adjusted to 30% PEG 300. Data were collected at 100 K at the Advanced Photon Source (APS) Northeastern Collaborative Access Team (NE-CAT) beamline 24-ID-C on a Pilatus 6M detector with a wavelength of 0.9795 Å, 0.2-s exposures, and 0.2° oscillation steps.

Diffraction data were integrated, scaled, and merged using DIALS (Supplementary Table 11).^49^ All steps from PALA synthesis to data processing are detailed in the Supplementary Methods.

### Cryo-EM and Crystallographic Model Building, Refinement, and Analysis

Model building and refinement were performed in Phenix^50^ and Coot^51^ (detailed in Supplementary Methods). The angle of each regulatory dimer was calculated from the dot product of the vectors connecting the center of mass (COM) of the residue-44 Cα atoms at the dimer interface and the Cα atoms of residue 95 on each chain. In Figure 3G, the “dimer angle” is 180 ° minus the average of the three regulatory dimers, with the uncertainty estimated by the standard deviation.

## Data Availability Statement

Coordinates and structure factors for crystal structures have been deposited in the Protein Data Bank with accession codes: 9EEH [http://doi.org/10.2210/pdb9EEH/pdb] (R: ATP/GTP/PALA) and 9EEJ [http://doi.org/10.2210/pdb9EEJ/pdb] (R: ATP/CP/SIN). Cryo-EM maps have been deposited to the Electron Microscopy Data Bank with accession codes: EMD-47956 [https://www.ebi.ac.uk/emdb/EMD-47956] (T), EMD-47957 [https://www.ebi.ac.uk/emdb/EMD-47957] (T: CTP/UTP/CP), EMD-47958 [https://www.ebi.ac.uk/emdb/EMD-47958] (R: CTP/UTP/CP/SIN), EMD-47959 [https://www.ebi.ac.uk/emdb/EMD-47959] (T: CTP/CP), EMD-47960 [https://www.ebi.ac.uk/emdb/EMD-47960] (R: CTP/CP/SIN), EMD-47961 [https://www.ebi.ac.uk/emdb/EMD-47961] (R: CP/SIN), EMD-47963 [https://www.ebi.ac.uk/emdb/EMD-47963] (R: ATP/CP/SIN), EMD-47964 [https://www.ebi.ac.uk/emdb/EMD-47964] (R: ATP/GTP/CP/SIN), EMD-47965 [https://www.ebi.ac.uk/emdb/EMD-47965] (R: ATP/GTP/CP), EMD-47966 [https://www.ebi.ac.uk/emdb/EMD-47966] (T: ATP/CP). The associated atomic models have been deposited to the Protein Data Bank with accession codes: 9EEK [http://doi.org/10.2210/pdb9EEK/pdb] (T), 9EEL [http://doi.org/10.2210/pdb9EEL/pdb] (T: CTP/UTP/CP), 9EEM [http://doi.org/10.2210/pdb9EEM/pdb] (R: CTP/UTP/CP/SIN), 9EEN [http://doi.org/10.2210/pdb9EEN/pdb] (T: CTP/CP), 9EEO [http://doi.org/10.2210/pdb9EEO/pdb] (R: CTP/CP/SIN), 9EEP [http://doi.org/10.2210/pdb9EEP/pdb] (R: CP/SIN), 9EEQ [http://doi.org/10.2210/pdb9EEQ/pdb] (R: ATP/CP/SIN), 9EER [http://doi.org/10.2210/pdb9EER/pdb] (R: ATP/GTP/CP/SIN), 9EES [http://doi.org/10.2210/pdb9EES/pdb] (R: ATP/GTP/CP), 9EEU [http://doi.org/10.2210/pdb9EEU/pdb] (T: ATP/CP). SEC-SAXS datasets are available on Zenodo with record numbers: 17684630 [https://doi.org/10.5281/zenodo.17684630] (T), 17684632 [https://doi.org/10.5281/zenodo.17684632] (R), 17684634 [https://doi.org/10.5281/zenodo.17684634] (R: CTP), 17684636 [https://doi.org/10.5281/zenodo.17684636] (R: CTP/UTP), 17684638 [https://doi.org/10.5281/zenodo.17684638] (R: ATP), 17684640 [https://doi.org/10.5281/zenodo.17684640] (R: ATP/GTP). Source data are provided as a Source Data file.

## Acknowledgements

The authors thank S. Meisburger, J. Bacik, D. Xu, J. Kaelber, and the Phenix team for technical discussions and B. Barstow, H. Wang, J. Dalal, and R. Schweinfurth for critical reading of this manuscript. We are grateful to the staff and resources at CHESS, CCMR, APS, NCCAT, and the Cornell NMR and Chemistry Mass Spectrometry Facilities. We thank C. Aplin, M. Lynch, R. Schweinfurth for assistance with data collection. SAXS/crystallography were conducted at the Center for High-Energy X-ray Sciences (CHEXS), which is supported by the National Science Foundation award DMR-2342336, and the MacCHESS facility, which is supported by award 1-P30-GM124166 from the National Institute of General Medical Sciences (NIGMS) and the National Institutes of Health (NIH). Cryo-crystallography was conducted at NE-CAT, which is funded by the NIGMS from the NIH (P30 GM124165), using resources of APS, a U.S. Department of Energy (DOE) Office of Science User Facility operated for the DOE Office of Science by Argonne National Laboratory under DE-AC02-06CH11357. Cryo-EM performed at NCCAT was supported by the NIH Common Fund Transformative High Resolution Cryo-Electron Microscopy program grant U24 GM129539. This work was supported by a half-time student appointment at CHEXS (to R.C.M.), Cornell Provost Diversity Fellowship (to. M.G.P.), NIH grant GM124847 (to. N.A.), and startup funds from Cornell University (to N.A.).

## Author Contributions

R.C.M. and M.G.P. purified proteins and performed SAXS experiments. R.C.M. performed SAXS data analysis, kinetic modeling of assay data, cryo-EM experiments, and cryo-EM data processing, model building, and refinement. M.G.P. performed activity assays, kinetic modeling, and crystallographic and cryo-EM model building and refinement. M.G.P. and N.B. grew crystals and performed crystallography experiments. X.P. synthesized PALA. N.A. supervised the project, acquired funding, and performed data analysis. R.C.M., M.G.P., and N.A. made figures and wrote the manuscript.

