## Supplemental Information for "Cooperativity in *E. coli* Aspartate Transcarbamoylase is Tuned by Allosteric Breathing"

**This PDF file includes:**

Supplementary Methods

Supplementary Figures 1 to 34

Supplementary Tables 1 to 12

Supplementary Movies 1 to 6

Supplementary References

### Supplementary Methods

#### Protein Expression and Purification

Tagless *E. coli* ATCase was expressed and purified using established protocols[1] with modifications. The native *E. coli pyrBI* coding sequence (RefSeq accession NC_000913.3; 1421 bp) was commercially synthesized by GenScript and subcloned into the pET-24a(+) expression vector using NdeI and NotI restriction sites via GenScript Express Cloning. Protein expression was performed in T7 Express BL21 (DE3) *E. coli* cells (New England Biolabs). A 250-mL starter culture was grown overnight at 200 rpm and 37 °C in LB (Miller) medium with 40 µg/mL kanamycin. Large-scale cultures were inoculated with 10 mL starter culture per 1 L of LB supplemented with 40 µg/mL kanamycin and grown at 200 rpm, 37 °C. Expression was induced at $OD_{600}$ = 0.4 with 1 mM isopropyl β-D-1-thiogalactopyranoside IPTG (final concentration) and allowed to proceed for 5 hours at 200 rpm and 37 °C. Cells were harvested by centrifugation (4000 × *g*, 4 °C, 30 min), flash frozen in liquid nitrogen, and stored at –80 °C.

Cell lysis and purification were performed in a cold room kept at 4 °C. Cells were resuspended in ~5 mL purification buffer (50 mM Tris-acetate pH 8.3, 1 mM tris(2-carboxyethyl)phosphine (TCEP)) per g cell paste with one protease inhibitor tablet (cOmplete Mini, EDTA-free, Roche) per ~10 mL of suspension. Cells were lysed using an Avestin Emulsiflex C3 homogenizer for 10–15 minutes at ~15000 psi, and the lysate was centrifuged at 25000 × *g* and 4 °C for 40 min. Clarified lysate was passed through a 0.2-µm filter and purified in three chromatographic steps using an ÄKTA pure fast protein liquid chromatography (FPLC) system. Lysate was loaded onto an anion-exchange column (HiPrep Q Sepharose Fast Flow 16/10, Cytiva) pre-equilibrated with purification buffer. ATCase was eluted at 1 mL/min over a 250-mL linear gradient (0 to 0.5 M NaCl) in purification buffer. Fractions containing ATCase were pooled and dialyzed overnight (16–18 hours) against fresh purification buffer at 4 °C. Finely ground ammonium sulfate ($AmSO_4$) powder was gradually added to dialyzed protein solution with stirring over 1 hour until 20% (w/v) saturation. The solution was filtered (0.2-µm) and loaded onto a hydrophobic interaction column (HiPrep Phenyl Sepharose Fast Flow 16/10, Cytiva) pre-equilibrated with purification buffer containing 20% $AmSO_4$. ATCase was eluted at 1 mL/min over a 15-mL linear gradient (20 to 0% $AmSO_4$) in purification buffer. Fractions containing ATCase were pooled and loaded onto a size-exclusion column (HiLoad Superdex 200 pg 16/600, Cytiva) pre-equilibrated in ATCase buffer (40 mM Tris-HCl pH 7.5, 15 mM $MgCl_2$, 1 mM TCEP). Fractions containing pure ATCase were pooled, and the concentration was determined with a theoretical extinction coefficient of $\varepsilon_{280nm}$ = 166215 $M^{-1}cm^{-1}$. All ATCase concentrations in this work are given as molar concentrations of the holoenzyme (308.4 kDa dodecamer). Previously, it was estimated that 5 g of ATCase can be obtained from 700 g of wet-packed *E. coli* cells.[2] Assuming an average cell volume of 1 $\mu m^3$ and a cell mass of $10^{-12}$ g, and further assuming that the cell paste consists only of cells (no water), this places an upper limit of approximately 23 µM for the cellular concentration of ATCase in *E. coli*.

#### Activity Assays

Enzyme kinetics were measured under solution conditions mimicking those used for SAXS and cryo-EM using a previously described colorimetric assay for carbamoyl aspartate (CAsp) production.[3] Fresh 10 mM carbamoyl aspartate (CP) solution was prepared in water at 25 °C, and freshly prepared color-reagent solutions (8 g/L 2,3-butanedionemonoxime (BDM) in 5% acetic acid and 5 g/L antipyrine (AP) in 50% sulfuric acid) were kept on ice. Five nucleotide conditions were examined: 1) no nucleotides, 2) 1.5 mM CTP, 3) 1.5 mM CTP, 1.5 mM UTP, 4) 5 mM ATP, and 5) 5 mM ATP, 1 mM GTP. Enzyme solutions (260 µL) and standard solutions (0.5 mL) were freshly prepared in assay buffer (40 mM Tris-HCl pH 7.5, 15 mM $MgCl_2$) in 96-deep-well plates. Enzyme solutions contained 1 nM ATCase, variable concentrations of aspartate (Asp), and nucleotide(s) as specified. Standards were prepared with 0, 5, 40, 80 and 120 µM CAsp. The plate was briefly centrifuged (1000 × *g*, 1 min), shaken on an orbital shaker for 10 minutes at ambient temperature, and equilibrated in a 25 °C water bath for 10 min. The color reagent was prepared by mixing the AP and BDM solutions in a 1:2 (v/v) ratio and kept on ice.

Reactions were initiated by addition of 240 µL 10 mM CP, yielding 0.5-mL reaction mixtures with 1 nM ATCase, 40 mM Tris-HCl pH 7.5, 15 mM $MgCl_2$, 4.8 mM CP, 0–150 mM Asp, and nucleotide(s) as described. Initial reaction rates were obtained by running reactions for exactly 5 minutes before quenching with 0.5 mL color reagent. The same volume of color reagent was added to standard solutions. The plate was centrifuged, wrapped in foil, and incubated in the dark at ambient temperature for ≥16 hours. The plate was placed in a 47 °C water bath with the foil removed to illuminate samples uniformly with an LED light pad for 15 minutes. The plate was re-wrapped in foil and stored on ice. Absorbance spectra (300–700 nm) were measured with an Agilent Cary 3500 Compact UV-Vis Spectrophotometer. A calibration curve (absorbance at 466 nm vs. CAsp concentration) was produced with the standards. Assay solutions were diluted 4-fold to fall within the detection range of the standard solutions. Saturation curves are reported as specific activity in µmol product $hr^{-1}$(mg protein)$^{-1}\times10^{-3}$ (equivalent to mmol product $hr^{-1}$(mg protein)$^{-1}$) for comparison with previous studies.

**Kinetic Modeling**

Plots of initial rates as a function of Asp concentration were obtained by averaging two sets of duplicate measurements (four total), and nonlinear curve fitting was performed in Python. The transition region of each curve was fit to the Hill equation (Supplementary Fig. 18, Supplementary Table 9),

$$v = \frac{V_{max}[S]^n}{\left(K_{1/2}\right)^n + [S]^n} \tag{1}$$

where the exponent *n* is the Hill coefficient (shown as $n_H$ in the text), $V_{max}$ is the maximal rate, [S] is the free substrate concentration, and $K_{1/2}$ is the substrate concentration at $V_{max}/2$. The Hill fits were initialized with parameters obtained from fits to the Michaelis-Menten (MM) equation,

$$v = \frac{V_{max}[S]}{K_m + [S]} \tag{2}$$

where $K_m$ is the Michaelis constant. Stable solutions for the Hill fits were obtained using one half of the $V_{max}$ and $K_m$ values from the MM fits as initial estimates for $V_{max}$ and $K_{1/2}$. As Eq. (1) reduces to Eq. (2) in the limit of $n_H = 1$, the curve obtained for ATCase in the presence of ATP and GTP were also modeled with the MM equation (Supplementary Fig. 18, Supplementary Table 9).

We note that high-activity states of ATCase exhibit signs of substrate inhibition at high Asp concentrations. This well-known behavior has previously been described by Pastra-Landis *et al.* using a modification of the Hill equation that accounts for substrate inhibition.[4] Although this model provides useful visual guides for the full range of our kinetics data (shown in Figure 2D), we did not use it for quantitative analysis as the parameters are underdetermined without independent measurement and modeling of the inhibition component of the kinetics curve at unphysiologically high Asp concentrations (hundreds of millimolar).

**PALA Synthesis and Characterization**
*N*-phosphonacetyl-L-aspartatic acid (PALA) was synthesized following a previously reported protocol (Supplementary Fig. 29).[5] The intermediate product **3** and final product (PALA, **4**) were characterized by nuclear magnetic resonance (NMR) and high-resolution electrospray ionization (HR-ESI) mass spectrometry (Supplementary Figs. 30-34). Solution $^{1}H$ and $^{31}P$ NMR spectra were collected on either a 400 MHz Bruker AVIII HD spectrometer with a BBFO probe, or a 500 MHz Bruker AVIII HD spectrometer with a BBO Prodigy cryoprobe. For $^{1}H$ NMR spectra, deuterated solvents were used as references of chemical shifts ($d_6$-DMSO: 2.50 ppm; $D_2O$: 4.79 ppm). Chemical shifts of $^{31}P$ spectra are given in ppm relative to 85% phosphoric acid (sealed in a glass capillary and used as an internal reference). Mass spectra were acquired on an Exactive Orbitrap mass spectrometer equipped with an Ion Max source and HESI II probe (Thermo Scientific).

Integration analyses were completed in MestReNova (v15.0.0-34764) under default settings. During multiplet analyses, the following $^{1}H$ NMR spectra were subjected to apodization before the Fourier transform step: **3**, -1.0 Hz exponential function and 1.3 Hz Gaussian function; **4** in $D_2O$, -0.3 Hz exponential function and 1.0 Hz Gaussian function. Results are summarized as follows.

$^{1}H$ NMR of intermediate product **3** ($d_6$-DMSO, 400 MHz): $\delta$ 8.32 (d, $J$ = 8.0 Hz, 1H), 4.46 (dd, $J$ = 14.0, 6.3 Hz, 1H), 4.09 – 3.95 (m, 4H), 2.96 (dd, $J$ = 21.4, 14.6 Hz, 1H), 2.91 (dd, $J$ = 21.5, 14.8 Hz, 1H), 2.61 (dd, $J$ = 17.9, 7.5 Hz, 1H), 2.55 (dd, $J$ = 17.9, 7.6 Hz, 1H), 1.40 (d, $J$ = 2.4 Hz, 18H), 1.22 (t, $J$ = 7.0 Hz, 6H). HRMS (ESI/Orbitrap) m/z: $[M + H]^+$ calcd for $^{12}C_{18}H_{35}{}^{16}O_8{}^{14}NP$, 424.209; found, 424.209.

The final product (PALA, **4**) after lyophilization at < 6.11 mbar vacuum for 2 days is a fluffy white crystalline powder. $^{1}H$ NMR in $d_6$-DMSO (500 MHz): $\delta$ 8.14 (d, J = 8.0 Hz, 1H), 4.54 (dd, J = 13.3, 6.5 Hz, 1H), 2.73 – 2.56 (m, 4H; partly overlap with DMSO peak; see spectra in Supplementary Figs. 30-34). $^{31}P$ NMR in $d_6$-DMSO (500 MHz): $\delta$ 17.87 (s). $^{1}H$ NMR in $D_2O$ (500 MHz): $\delta$ 3.01 (dd, $J$ = 17.3, 5.9 Hz, 1H), 2.96 (dd, $J$ = 17.2, 5.5 Hz, 1H), 2.83 (dd, $J$ = 20.6, 14.3 Hz, 1H), 2.79 (dd, $J$ = 20.4, 14.1 Hz, 1H). $^{31}P$ NMR in $D_2O$ (500 MHz): $\delta$ 15.33 (s). HRMS (ESI/Orbitrap) m/z: $[M - H]^-$ calcd for $^{12}C_6H_9{}^{16}O_8{}^{14}NP$, 254.006; found, 254.006.

**SAXS**
*Beamline Setup and Data Processing.* Samples were screened on a laboratory source (Xenocs BioXolver) before data collection at the Cornell High-Energy Synchrotron Source (CHESS) ID7A station using a 250-µm square X-ray beam with a flux of $\sim 10^{12}$ photons $s^{-1}$ $mm^{-1}$ at the sample position. SAXS was performed on 7 different occasions with X-ray energies of 9.8–11.3 keV (Supplementary Tables 1-6). Scattering images were collected on an Eiger 4M detector covering a range of $q \approx 0.01$–$0.55$ Å$^{-1}$, where the momentum transfer variable is defined as $q = 4\pi/\lambda \sin\theta$, $\lambda$ is the X-ray wavelength, and $2\theta$ is the scattering angle. Data integration was performed in BioXTAS RAW,[6] and processing was performed in custom Python scripts. Integrated scattering profiles were normalized by transmitted intensities, and scattering from a matched buffer was used to yield the background-subtracted protein scattering profile, $I(q)$. The radius of gyration $R_g$ and forward scattering intensity $I(0)$ were determined by Guinier analysis, and curve-fitting uncertainties were used to estimate associated errors.

*SAXS Titrations.* A concentration series first showed that an ATCase concentration of 8 μM provides sufficient signal-to-noise and minimal interparticle effects in the mid-$q$ and low-$q$ regions, respectively. Titration experiments were performed at 4 ºC with 40-μL protein solutions freshly prepared in ATCase buffer with variable concentrations of ligands (e.g., substrate, substrate analog, nucleotides, Supplementary Tables 1-3). Samples were centrifuged (15,000 × $g$, 4 °C, 10 min) immediately before injection into an in-vacuum flow-cell. Background scattering was measured with matched buffer solutions. Ten to twenty 1-s exposures were collected with sample oscillation. Individual frames without detectable radiation-induced changes and averaged. Singular value decomposition (SVD) was performed in Python, and the position of the second peak in Kratky curves ($Iq^2$ vs $q$) was determined by a Gaussian fit.

*Chromatography-Coupled SAXS.* To obtain high-quality scattering profiles for direct comparison with high-resolution models, SEC-SAXS was performed under conditions identified by titration experiments that saturate structural transitions. Samples (50 μL of 56 μM ATCase in ATCase buffer with ligand concentrations adjusted for each condition of interest) were centrifuged (15,000 × $g$, 10 min, 4 °C) and loaded onto a Superdex 200 10/300 GL column (Cytiva) operated by a GE Akta Purifier at 4 °C that was pre-equilibrated in a matched buffer. The elution flowed directly into an in-vacuum X-ray sample cell held at 4 °C at 0.2–0.6 mL min$^{-1}$. Two to three second exposures were collected throughout elution, and scattering profiles of the elution buffer were averaged to produce a background-subtracted SEC-SAXS chromatogram. Six conditions were examined (Supplementary Tables 4-6): 1) no ligands, 2) 500 μM CP, 10 mM succinate, 3) 500 μM CP, 10 mM succinate + 1.5 mM CTP, 4) 500 μM CP, 10 mM succinate + 1.5 mM CTP, 1.5 mM UTP, 5) 500 μM CP, 10 mM succinate + 5 mM ATP, and 6) 500 μM CP, 10 mM succinate + 5 mM ATP, 1 mM GTP.

Evolving-factor analysis (EFA)[7] and regularized alternating least squares (REGALS)[8] were applied in Python to separate overlapping elution peaks in a model-independent manner. The elution window of each species was first estimated by EFA, and REGALS was used to model each species for separation. In all datasets, the dodecameric complex was the dominant species (Supplementary Figs. 4-6, Supplementary Tables 4-6), and the corresponding REGALS-extracted scattering profile was used for further structural analysis. Fits to high-resolution models were performed in CRYSOL3[9] with 50 spherical harmonics, 500 points between 0 and 0.25 Å$^{-1}$, and the default electron density of water. CRYSOL3 overcomes a key limitation in previous versions of CRYSOL in evaluating the hydration shell of proteins such as ATCase, which are highly non-globular in shape and contain a large central opening.

**Cryo-EM**

*Sample Preparation and Data Collection*. Cryo-EM was performed on ATCase in identical buffer conditions used for SAXS (Supplementary Tables 7-8) with enzyme concentration of 2 μM to limit particle density on the grid. Seven ligand conditions were prepared in ATCase buffer: 1) no ligands, 2) 500 μM CP, 10 mM succinate, 3) 500 μM CP, 10 mM succinate + 1.5 mM CTP, 4) 500 μM CP, 10 mM succinate + 1.5 mM CTP, 1.5 mM UTP, 5) 500 μM CP, 10 mM succinate + 5 mM ATP, 6) 500 μM CP, 10 mM succinate + 5 mM ATP, 1 mM GTP, and 7) 500 μM CP + 5 mM ATP, 1 mM GTP. For each condition, 20-40 μL solutions were prepared and centrifuged (15,000 × $g$, 4 °C, 15 min), kept on ice, and applied to grids within 1 hour of preparation.

Grids were prepared using an FEI Vitrobot Mark IV (100% humidity, 4 °C). QuantiFoil R 1.2/1.3 300-mesh copper grids were glow-discharged on a PELCO easiGlow system for 45 s with 20-mA current, functionalized with graphene oxide (GO) as previously described,[10] and used within 2 hours. We found that ATCase is prone to dissociation and structural perturbation in the cryo-EM grid environment, and the use of GO grids was essential for minimizing these issues. Four µL of the sample was applied onto the grid, incubated for 90 s, blotted for 3.5 s, and plunge-frozen in liquid ethane cooled by liquid nitrogen.

Initial grid screening was performed on a Talos Arctica (Thermo Fisher) at the Cornell Center for Materials Research (CCMR), operating at 200 keV with a Gatan K3 detector and BioQuantum energy filter (20 eV slit) at a nominal magnification of 63,000× (nominal pixel size of 1.31 Å $pix^{-1}$). Movies were collected with a –0.8 to –2.0 µm defocus range and a total dose of 50 $e^{-}$ $Å^{-2}$ over 50 frames, with a 3 × 3 multishot scheme in SerialEM[11]. For the unliganded condition, the final dataset consisting of 2,136 movies (Supplementary Table 7) was collected with these parameters at an exposure time, frame time, and dose rate of 2.036 s, 0.04072 s, and 24.56 $e^{-}$ $Å^{-2}$ $s^{-1}$, respectively.

Final datasets for other conditions were collected on a Krios G4 (Thermo Fisher) at the National Center for CryoEM Access and Training (NCCAT) operating at 300 keV with a Gatan K3 detector, BioQuantum energy filter (20 eV slit), and 2.7 mm spherical aberration. Image acquisition parameters are summarized in Supplementary Tables 7-8. Movies were collected with a defocus range of –0.8 to –2.3 µm using a Leginon multishot scheme.[12]

*Data Processing.* Datasets for all seven conditions were processed in cryoSPARC 4[13] and RELION 4[14] (Supplementary Figs. 8-13, 26). Patch motion correction and contrast transfer function (CTF) estimation were first performed in cryoSPARC. Micrographs were curated based upon statistical image parameters (e.g., CTF fit resolution, ice thickness estimates) and visual inspection, retaining ~80% of the movies for each dataset. Particles were picked by template and blob particle picking, extracted with box sizes of 480 × 480 pixels (super-resolution) or 360 × 360 pixels (real pixel size), and Fourier cropped to 240 × 240 and 180 × 180 pixels, respectively. Particles were filtered via iterative 2D classification with a class probability cutoff of 0.95 followed by *ab initio* reconstruction with two classes to remove 'junk' particles. Duplicated particles with separation distance below 25 Å were removed. The retained particles were re-extracted with box sizes of 480 × 480 pixels (super-resolution) or 360 × 360 pixels (real pixel size) with an average particle density of ~ 200 good particles per micrograph for all datasets.

Micrographs initially motion-corrected in cryoSPARC were motion-corrected again in RELION 4 using its own implementation of MotionCorr2 with 7 by 5 patches. Particle coordinates from cryoSPARC were converted to RELION star format with pyem,[15] and particles were re-extracted from the newly motion-corrected micrographs in RELION 4 with box sizes of 480 × 480 pixels (super-resolution) or 360 × 360 pixels (real pixel size). 3D classification was performed on the particles, using refined volumes from initial processing in cryoSPARC for alignment, using 3, 5, and 8 classes. Particles in classes without secondary structure features were discarded, and the remaining classes were sub-classified based upon quaternary structure. 3D classification yielded a mixture of T-state-like and R-state-like classes for three datasets: 1) CTP/UTP/CP/succinate, 2) CTP/CP/succinate, and 3) ATP/GTP/CP. Particles in like classes were combined for consensus refinement. For data collected with SerialEM, this was followed by CTF parameter refinement (anisotropic magnification, beam tilt, trefoil, spherical aberration, per-micrograph astigmatism, and per-particle defocus), with particles split into 9 optics groups based on the 3 x 3 multishot position of their originating micrographs. For data collected at NCCAT with Leginon, particles were split into 140 optics groups based on the measured image shift for each acquisition. Bayesian polishing parameters were trained on 10,000 particles of the resulting refinement before Bayesian polishing was performed to obtain shiny particles. This process of 3D refinement, CTF parameter refinement, and Bayesian polishing was repeated twice to obtain the final shiny particle stack and volume, with C3 symmetry relaxation applied during the initial alignment phase of the final 3D refinement to obtain final low-noise consensus maps without imposed symmetry. C3 symmetry was only imposed for the following cases where it led to an improvement in the map quality: all T-state maps (ligand-free, CTP/CP, CTP/UTP/CP, ATP/CP) and one R-state map (ATP/GTP/CP). The resulting consensus maps for all states were sharpened using the RELION post-processing tool (masked and unmasked half-map FSC curves are shown in Supplementary Figs. 8-13, 26). To provide an independent assessment of resolution, including model-map FSCs, we also used phenix.mtriage to compute FSC curves from the full maps, half-maps, and corresponding atomic models (Supplementary Fig. 14).

**3D Variability Analysis**

3D variability analysis (3DVA)[16] was performed in cryoSPARC 4[13] to examine the continuous heterogeneity within each structural state. Shiny particles from the final refinement in RELION 4 performed in the absence of imposed symmetry were imported into cryoSPARC for non-uniform 3D refinement into the highest possible symmetry of D3. The resulting particle stack was symmetry-expanded, removing the imposed symmetry and multiplying the size of the particle stack by the order of the symmetry operator, to assist in identifying potential sources of asymmetry, before being subjected to 3DVA with 1, 3, and 5 classes and a resolution filter of 4.0 Å. The resolution cutoff was determined empirically by iteratively increasing the filter resolution cutoff until portions of the volume were lost throughout the 3DVA trajectory. The resulting trajectories were visualized by discretizing each mode into 20 uniformly spaced sub-volumes using the 3DVA display function in cryoSPARC. Each structural state displayed a single primary 3DVA mode (Supplementary Movies 1-6), and all other modes represented noise. The 3DVA performed with a single class was therefore used for subsequent analyses.

The dominant 3DVA mode primarily describes variation in the center-of-mass distance between the catalytic trimers, which can be captured effectively by rigid-body fitting. Thus, each of the 12 chains in the corresponding consensus models were rigid-body fit into every sub-volume in Chimera[17]. The quality of the docking models was inspected visually using secondary structure features. For each rigid-body model, the Cα center-of-mass (COM) distance between the top and bottom catalytic trimers was calculated across the trajectory and shown to have a linear relationship with the 3DVA latent variable (Supplementary Figs. 8-13, 26). The linear relationship was then used to convert the latent variable coordinate to the inter-trimer distance. Each distribution of trimer distances could be described as normal, and the mean and standard deviation obtained from a Gaussian fit were used as the average trimer distance and estimated uncertainty reported in Figure 3G. A similar calculation was performed using crystal structures of the T-state (4fyy, 4fyx, 4fyv, 4fyw, 2fzc, 2fzk, 2fzg, 5at1, 1za1, 6at1, 3at1, 1raa, 1za2, 4at1), which have an average trimer distance of 45.4 ± 0.2 Å, and the R-state (4kgx, 2h3e, 4kh1, 4kgz, 4kh0, 2ipo, 4kgv, 8at1, 1d09, 1at1, 2at1, 7at1), which have an average trimer distance of 55.9 ± 0.3 Å (ranges are shown as shaded bars in Figure 2C).

**Cryo-EM Model Building and Refinement**

Initial models were generated using crystal structures from this work or others as described below. For the ligand-free T-state and all open conformations, components of crystal structures were docked into sharpened maps in Chimera[18] followed by iterative real-space refinement and model building in Phenix[19] and Coot.[20] In the first refinement iteration, the catalytic subunits, Zn-domains (regulatory subunit residues 101–153 and $Zn^{2+}$), and the nucleotide-binding domains (regulatory subunit residues 1–100, nucleotides, and $Mg^{2+}$) were refined as rigid bodies. Subsequent iterations incorporated global minimization and local-grid searches, with the addition of atomic-displacement parameters in the final iteration. In all conditions, the map resolution was highest at the active sites and lowest at the nucleotide-binding domains (which includes the regulatory N-termini). In substrate-bound conditions, map quality surrounding the succinate molecule varied from active site to active site, consistent with the fact that it is a substrate analog (Supplementary Figs. 17, 27). Nucleotides were copied from crystal structures as described below, and CP and succinate (generated by removing the amine group from Asp) were copied from a crystal structure of *Trypanosoma cruzi* ATCase (PDB: 6jks).[21] All ligands and regulatory N-termini were restrained to their initial geometries at 0.1 sigma when present. To capture asymmetry, domains/motifs/ligands were manually divided into non-crystallographic symmetry (NCS) groups that yielded good model-to-map fits. $Mg^{2+}$ and $Zn^{2+}$ coordination bonds were restrained based upon the starting model. Ligand restraints were generated using the Grade Web Server.[22] Sidechains and NCS groups were adjusted manually to remove clashes. Model geometry was validated with the wwPDB[23] server via Phenix.

For the ligand-free T-state, the starting model was generated using chains A and B of a T-state structure of *E. coli* ATCase (PDB: 6at1).[24] Refinement was performed with 2 NCS groups: (1) the catalytic subunits, and (2) regulatory subunits. Residues 1–11 were not included in the regulatory chains due to limited map coverage.

For the ATP/GTP-bound R-state, the starting model was generated using chains A, B, and D of the ATP/GTP-bound crystal structure obtained in this work (PDB: 9eeh), and refinement was performed with 4 NCS groups: (1) the CP domains (catalytic residues 1–135, 292–310, and CP), (2) Asp domains (catalytic residues 136–291 and succinate), (3) nucleotide-binding domains (regulatory residues 4-100, ATP, GTP, and $Mg^{2+}$), and (4) Zn domains (regulatory residues 101–153 and $Zn^{2+}$). The ATP/GTP-bound crystal structure was an appropriate starting model for the nucleotide-binding region because ATP/GTP binding is sufficiently potent to preserve the bent regulatory-dimer conformation, even as the crystal lattice compresses the R-state through the flexible inter-domain linkers (a behavior not seen for ATP alone or for any other nucleotide combination). For the nucleotide-free R-state and ATP-bound R-state, the starting models were generated using chains B and L of the ATP-bound crystal structure obtained in this work (PDB: 9eej), with residues 1–11 and 1–9 of the regulatory chains excluded in the respective models due to limited map coverage. Refinement was then performed with the same NCS groups as the ATP/GTP-bound R-state.

For the CTP-bound R-state and CTP/UTP-bound R-state, the starting models were generated using catalytic subunits (chain B) and Zn domains (chain L residues 101-153 and $Zn^{2+}$) of the ATP-bound crystal structure obtained in this work (PDB: 9eej). The starting model for the CTP/UTP-bound R-state was then completed with the nucleotide-binding domain (residues 7-100, CTP, UTP, and $Mg^{2+}$ of chains B and D) of the CTP/UTP/PALA-bound structure of *E. coli* ATCase (PDB: 4kh1),[25] and refinement was performed with 6 NCS groups: (1) the catalytic subunits, (2) Zn domains (regulatory residues 101–143 and $Zn^{2+}$), (3, 4) nucleotide-binding domains (regulatory residues 7–100, CTP, UTP, and $Mg^{2+}$) with two unique H2-helix conformations, and (5, 6) two distinct C-terminal conformations of the regulatory chains (regulatory residues 144–153). The starting model for the CTP-bound R-state was completed using the nucleotide-binding domain (chain D residues 12–100) of a CTP/PALA-bound structure of *E. coli* ATCase (PDB: 4kgx),[25] wherein a second CTP molecule and a $Mg^{2+}$ ion was first real-space refined into a large patch of unmodeled difference density ($F_O - F_C$) for a site-2 nucleotide in the deposited electron-density maps. Refinement was performed with 10 NCS groups: (1) the CP domains (catalytic residues 1–135, 292–310, and CP), (2) Asp domains (catalytic residues 136–291 and succinate), (3) Zn domains (regulatory residues 101–143 and $Zn^{2+}$), (4) 50s loops of nucleotide-binding domains (regulatory residues 46-55), (5-7) the remaining regions of the nucleotide-binding domains (regulatory residues 12–45, 56–100, CTP, and $Mg^{2+}$) with three unique H2-helix conformations, and (8-10) three distinct C-terminal conformations of the regulatory chains (regulatory residues 144–153).

The starting model for the ATP/GTP/CP-bound state was generated using catalytic subunits of a CTP/CP-bound structure of *E. coli* ATCase (PDB-REDO: 1za2, chain A)[26,27], and regulatory subunits from the ATP/GTP-bound structure obtained in this work (PDB: 9eeh, chains B and D, residues 4–153). The model was refined with 2 NCS groups matching those of the T-state. Classification of this dataset also yielded a T-state-like class, but the densities for the nucleotides and regulatory N-termini were more consistent with ATP bound in both sites 1 and 2.

Starting models for the T-states obtained from pyrimidine-containing conditions were generated using catalytic subunits from a crystal structure of *E. coli* ATCase (PDB: 1za2, chain A). For the CTP-bound T-state, the initial model also contained the nucleotide-bound regulatory subunits from the CTP-bound R-state EM model (chains J and L), whereas for the CTP/UTP-bound T-state, the nucleotide-binding domains (regulatory residues 1–100) were swapped with those of the CTP/UTP-bound R-state EM model (chains J and L). For both models, the sharpened and unsharpened maps displayed density in the active site that supported the placement of CP but not succinate. The geometry and relative orientation of CP was obtained from PDB: 6jks[21], consistent with other EM models, and was placed based upon the active site density. Each initial model was relaxed into its corresponding sharpened map in ISOLDE[28] and further refined via local grid searching and global minimization over a limited number refinement cycles in Phenix (<20). Ligands, nucleotide effectors, and metals were removed from the resulting models and prepared for energy minimization by inserting missing hydrogens reflecting a pH of 7.0. Energy minimization was performed in OpenMM[29] with the Amber force field[30] and the TIP3P water model[31]. Position restraints were applied to maintain structural integrity during minimization, with a force constant of 1000 $kJ/mol/nm^2$ for the backbone atoms (N, Cα, C, O) and 500 $kJ/mol/nm^2$ for the side chains, allowing controlled flexibility. A Langevin integrator at 300 K with a friction coefficient of 1 $ps^{-1}$ and a time step of 2 fs was used for the energy minimization, which proceeded until the system's energy was reduced to a tolerance of 10 kJ/mol/nm, with a maximum of 5000 iterations. The minimized structure reflected reduced steric clashes and optimized geometries. The removed ligands, nucleotide effectors, and metals from the starting models were docked into the minimized structure followed by a final round of refinement in Phenix using local grid search and global minimization over 3 macrocycles. NCS refinement was not used.

**Crystallography**

*Co-crystallization of ATCase with PALA, ATP, and GTP.* Crystallization conditions were designed to mimic SAXS solution conditions. The protein solution was prepared with 16 μM ATCase in ATCase buffer with 10 mM ATP, 2 mM GTP, and 2 mM PALA. High-throughput crystallization screens were performed using a Mosquito (TTP Labtech) robot using a sitting-well tray, where 160 nL each of the precipitant and protein solutions were mixed before sealing and storing at ambient temperature. Small rod-shaped crystals appeared within 24 hours in condition F4 of the PEG/Ion HT screen (Hampton Research) containing 20% (w/v) PEG 3350 and 8% (v/v) Tacsimate pH 7.0. Optimization around the hit condition (17-22% PEG 3350) was performed in hanging-drop trays (VDX, Hampton Research) using 500-μL reservoir solutions and 2-μL hanging drops consisting of a 1:1 mixture of protein and precipitant solutions. Similar rod-shaped crystals appeared in all conditions within 24 hours and grew to 10-20 μm in the longest dimension. A 500-μL seed stock was prepared from two crystallization drops containing 21% (w/v) PEG 3350 using the Seed Bead Kit (Hampton Research), and 10- and 100-fold dilutions were prepared in fresh precipitant solution. Hanging-drop trays were prepared with 4-μL drops containing 1 μL precipitant solution (8% (v/v) Tacsimate pH 7.0 and 9-14% (w/v) PEG 3350), 1 μL seed solution, and 2 μL protein solution. After 5–7 days at ambient temperature, large hexagonal rod-shaped crystals (~500×300×300 μm) appeared.

*Data Collection.* Room-temperature diffraction data were collected at the CHESS ID7B2 station on an Eiger2 16M detector with an X-ray wavelength of 0.9185 Å, 10-ms exposures, and 0.1° oscillation steps. The condition described above produced two visually indistinguishable crystal forms, often in the same drop, both indexing in *P*321 but differing in unit cell parameters along the *c* axis: contracted ($c$ = 153.0 Å) or expanded ($c$ = 166.8 Å) (Supplementary Fig. 21B-C). The contracted form yielded better data quality and higher resolution, yielding the 2.19-Å resolution structure we report in this study. Diffraction data were integrated, scaled, and merged in DIALS[32] (Supplementary Tables 11-12). The rationale for the chosen resolution cutoff and additional validation is discussed in detail in Patterson (2024)[33].

*Co-crystallization of ATCase with CP, Succinate, and ATP.* High-throughput crystallization screening was performed as described above using a protein solution containing 16 μM ATCase in 40 mM Tris-HCl pH 7.0, 20 mM CP, 20 mM succinate, 30 mM $MgCl_2$, and 10 mM ATP, was mixed with precipitant solutions in a 1:1 ratio. Within 3 days, small needle-shaped (~1×10 μm) crystals appeared in condition A5 of the PEGRx HT screen (Hampton Research) containing 25% (w/v) PEG 300 and 0.1 M Bis-Tris pH 6.5. This condition was optimized in hanging-drop trays, ultimately yielding large crystals (~700×200×200 μm) at pH 6.5 and 20-25% PEG 300. Crystals were cryoprotected by soaking for 5-10 s in solutions matching the crystallization drops adjusted to 30% PEG 300 before being flash frozen in liquid nitrogen.

*Data Collection.* Data collection was performed at the Advanced Photon Source (APS) Northeastern Collaborative Access Team (NE-CAT) beamline 24-ID-C at 100 K on a Pilatus 6M detector with a wavelength of 0.9795 Å, 0.2-s exposures, and 0.2° oscillation steps. Diffraction data were integrated, scaled, and merged in $P2_12_12_1$ using DIALS (Supplementary Table 11).[32]

**Crystallographic Model Building and Refinement**

For diffraction data obtained with PALA, ATP, and GTP, initial phase estimates were obtained using molecular replacement in Phaser[34] with the ATP-bound *P*321 ATCase structure (PDB: 4kh0)[25] as the search model with residues 1-11 removed from the regulatory subunits. Iterative refinement of atomic coordinates, isotropic B-factors, ligand occupancies, and translation/libration/screw-axis (TLS) parameters was performed in Phenix[19] alongside model building in Coot.[20] The asymmetric unit was divided into six TLS groups: (1,2) the catalytic subunits, (3,4) Zn domains (regulatory residues 100-153 and $Zn^{2+}$), and (5,6) nucleotide-binding domains (regulatory residues 4-100, ATP, GTP, and $Mg^{2+}$). NCS torsion-angle restraints were imposed throughout refinement and grouped by subunit. Between the NCS-related chains, several local differences were automatically identified and allowed to refine independently. The electron-density map provided clear evidence of PALA in both active sites and ATP at the first allosteric site in both regulatory subunits, similar to the initial search model. In contrast to the search model, a contiguous patch of difference density ($F_O - F_C$) adjoined the second allosteric binding sites across the dimer interface, which was consistent with a non-canonical GTP-GTP base-pairing interaction. Additional difference density suggested that an enclosed binding pocket was formed above the plane of the nucleobases by a domain-swapped N-terminal antiparallel β-sheet. To model the base-pairing interaction, gentle bond-length restraints of 2.7 ± 0.5 Å were imposed on the non-H atoms involved in the base pair. This ensured the interaction was not forced and overrode any potential modeling bias resulting from clashes between the protons at the N1 positions of the opposing nucleobases (hydrogens were not modeled explicitly). The interatomic distances ultimately refined to ~2.5 Å each. Attempts to model an ATP-GTP interaction between the second allosteric binding sites resulted in major clashes between the opposing amino groups and large patches of difference density (Supplementary Fig. 22). Similar to the starting model, the electron-density suggested the ATP/GTP α- and γ-phosphates formed a 6-coordinate octahedral $Mg^{2+}$ center including 2 axial waters. All $Zn^{2+}$ and $Mg^{2+}$ coordination-bond lengths and angles were restrained to within 0.1-0.5 Å and 5-10° of the idealized geometries. Waters were iteratively refined and manually curated. Model geometry was validated with the wwPDB[23] server via Phenix. Polder[35] analysis was performed in Phenix. The final model contains residues 1-310 in the catalytic chains (A and C), residues 4-153 in the regulatory chains (B and D), 2 PALA molecules, 2 ATP molecules, 2 GTP molecules, 2 $Mg^{2+}$ ions, 2 $Zn^{2+}$ ions, and 321 water molecules.

For diffraction data obtained with CP, Succinate, and ATP, molecular replacement was performed with search models generated by symmetry expansion of an ATP/PALA-bound structure in *P*321 (PDB: 4kh0),[25] which was then separated into halves, each comprised of 1 catalytic trimer and 3 regulatory monomers. Atomic coordinates, isotropic B-factors, and TLS parameters were refined as described for our other crystal structure. The asymmetric unit was divided into 18 TLS groups: (1-6) the catalytic subunits, (7-12) Zn domains, and (13-18) nucleotide-binding domains, with domains as defined above. NCS and metal-coordination restraints were implemented similarly to our other structure. The electron density clearly supported the presence of CP and succinate in each active site (Supplementary Fig. 19). Similar to the previous ATP/PALA-bound structure (PDB: 4kh0),[25] electron density supported the modeling of two nucleotides coordinating a $Mg^{2+}$, although density for the nucleobases displayed varying levels of disorder (Supplementary Fig. 20). Likewise, electron density for the N-terminal residues of the regulatory subunits varied across the structure, and residues were added or removed from each chain accordingly. The final model contains residues 10-153 (D), 6-153 (E), 9-153 (H), 6-153 (I), 6-153 (J), and 4-153 (L) in the regulatory chains, residues 1-310 in the catalytic chains (A, B, C, F, G, and K), 12 CP molecules, 12 succinate molecules, 12 ATP molecules, 6 $Mg^{2+}$ ions, and 6 $Zn^{2+}$ ions.

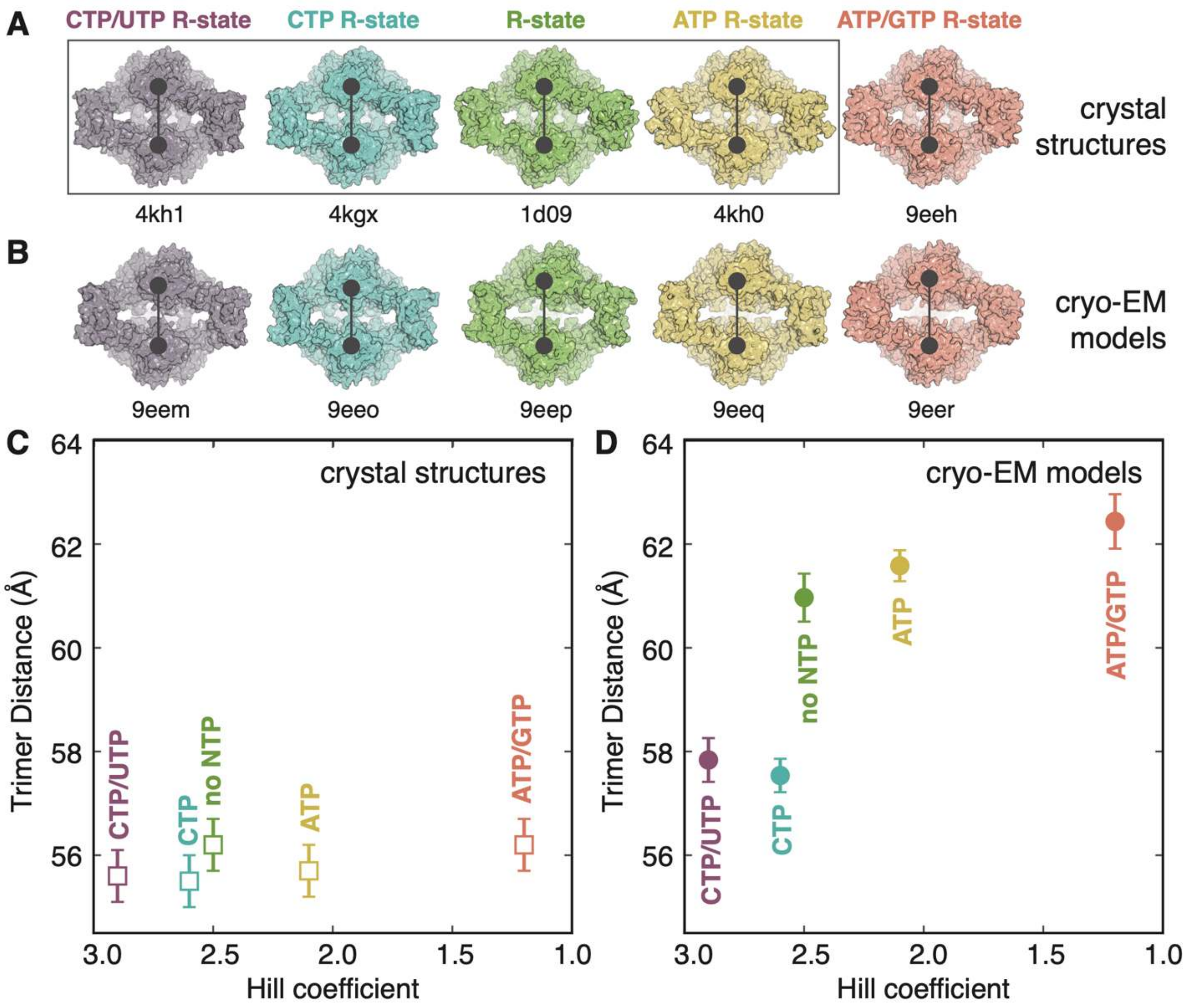


**Supplementary Figure 1. Nucleotide-dependent regulation of *E. coli* ATCase involves flexible motions that are prevented in the crystal lattice.**

Comparison of **(A)** crystal structures and **(B)** cryo-EM models of the R-state (PDB accession codes shown below). The "trimer distance" is defined as the distance between the centers of mass each catalytic trimer, a metric that utilizes the coordinates of many atoms. Structures from previous studies are boxed; all others were determined in this study. **(C)** Crystal structures of the R-state consistently display trimer distances of ~56 Å, regardless of ligand identity. The small, sub-angstrom variations (± 0.5 Å) differences in trimer separation are difficult to interpret; the error bars shown are the estimated coordinate errors.[24,36] Moreover, no correlation is observed between trimer distance and enzyme cooperativity, as represented by the Hill coefficient (values from this study). **(D)** In contrast, cryo-EM (this study) reveals that, in solution and free from crystal contacts, the R-state is highly flexible. Nucleotides induce a gradient of conformations differing by up to ~5 Å in trimer separation. Pyrimidines compress the R-state, while purines expand it, and these structural changes that correlate with changes in cooperativity. Shown are the mean and full-width-at-half-maximum values of the trimer distance distributions from 3D variability analysis (Figure 2C).

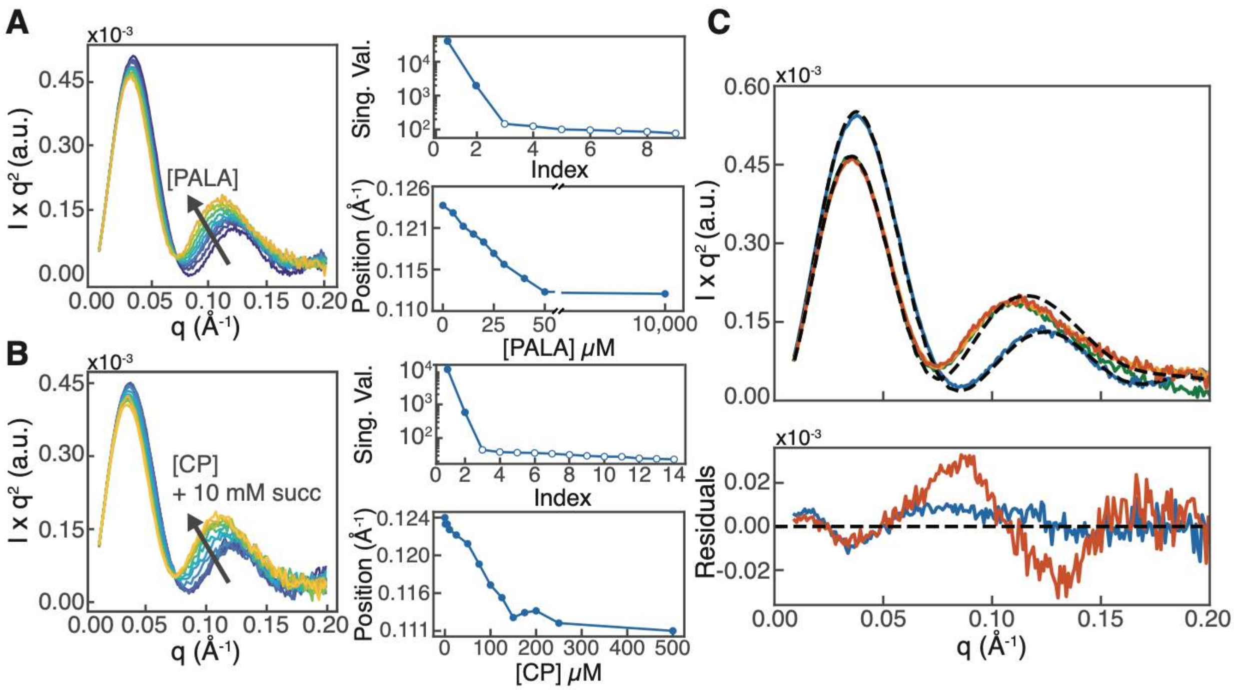


**Supplementary Figure 2. SAXS analysis of *E. coli* ATCase reveals a substrate-induced two-state transition to an R-state conformation, which is more expanded in solution than in the crystal.**

All measurements were made in 40 mM Tris-HCl pH 7.5, 15 mM $MgCl_2$, 1 mM TCEP with ligand concentrations adjusted as described. **(A)** Scattering profiles of 8 μM ATCase were collected at increasing concentrations of PALA (0 - 10 mM), a tight-binding bisubstrate analog (*left*). Singular value decomposition (SVD) yielded two significant singular values, indicating that the structural change is consistent with a two-state transition (*top right*). The second Kratky peak shifted to lower *q*, consistent with structural expansion, and saturated around 50 μM PALA (*bottom right*). **(B)** Scattering profiles of 8 μM ATCase at increasing concentrations of substrate CP (0 - 500 μM) with a fixed concentration of succinate (10 mM), an analog of the second substrate Asp (*left*). As in panel A, SVD of the dataset yielded two significant singular values (*top right*), and the second Kratky peak shifted to lower *q*, saturating around 250 μM CP (*bottom right*). **(C)** The scattering profile of ligand-free ATCase (*blue*) agrees well with the crystal structure of the ligand-free T-state[24] (PDB: 6at1, *dashed*). In contrast, the PALA-bound R-state crystal structure[37] (PDB: 1d09, *dashed*) fits poorly to the scattering profile obtained with 50 μM PALA (*orange*). This profile overlays with those obtained at 10 mM PALA (*green*) and at 500 μM CP, 10 mM succinate (*red*), indicating that the both PALA and CP/succinate induce transitions to the same R-state. Data parameters detailed in Supplementary Table 1.

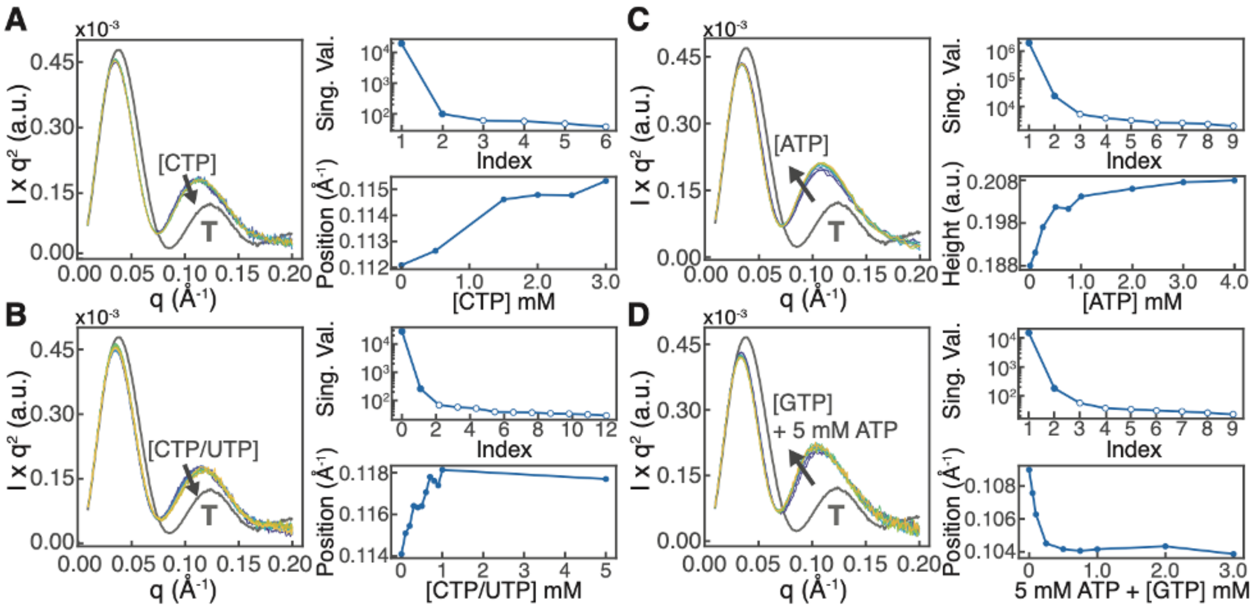


**Supplementary Figure 3. SAXS analysis of *E. coli* ATCase reveals that nucleotides perturb the R-state, with pyrimidines inducing compaction and purines inducing expansion.**

Nucleotides were added to ATCase in the R-state (8 µM ATCase with 500 µM CP and 10 mM succinate). For reference, the scattering of the ligand-free T-state is shown in all Kratky plots (*gray*, labeled "T"). All measurements were made in 40 mM Tris-HCl pH 7.5, 15 mM $MgCl_2$, 1 mM TCEP with ligand concentrations adjusted as described. **(A)** Addition of 0–3 mM CTP to ATCase in the R-state results in subtle changes in scattering (arrow) (*left*). SVD revealed up to two significant singular values (*top right*), consistent with a small but detectable shift of the second Kratky peak toward higher *q* (*bottom right*). This shift saturated above ~1.5 mM CTP without reaching the T-state value of 0.122 Å$^{-1}$, indicating that CTP induces modest compaction of the R-state. **(B)** Equimolar addition of CTP and UTP from 0–5 mM produced a more pronounced change in scattering than CTP alone (*left*). Two significant singular values were observed (*top right*), and the second Kratky peak shifted clearly to ~0.118 Å$^{-1}$ above ~1 mM CTP/UTP (*bottom right*), indicating that this pyrimidine pair induces compaction of the R-state. **(C)** Addition of 0–4 mM ATP led to a subtle but distinct change in scattering (*left*). Two significant singular values were observed (*top right*), and the second Kratky peak increased in intensity (*bottom right*). This transition saturated above ~3 mM ATP. **(D)** In contrast, addition of 0–3 mM GTP together with 5 mM ATP produced a more pronounced change in scattering than ATP alone (*left*). Two significant singular values were observed (*top right*), and the second Kratky peak shifted clearly to lower *q*, with the transition complete above ~250 µM GTP (*bottom right*), indicating that this purine pair drives a clear expansion of the R-state. Data parameters detailed in Supplementary Tables 2-3.

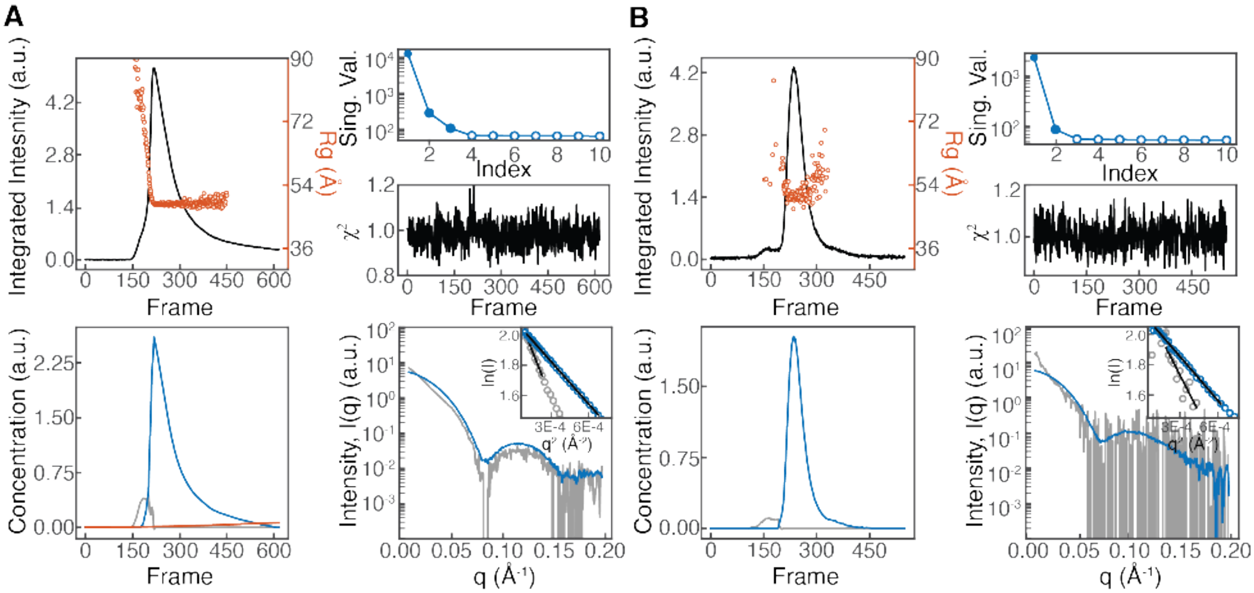


**Supplementary Figure 4. SAXS with in-line size exclusion chromatography yields high-quality scattering profiles of *E. coli* ATCase in the ligand-free T-state and nucleotide-free R-state.**

Both the sample buffer and elution buffer were 40 mM Tris-HCl pH 7.5, 15 mM $MgCl_2$, 1 mM TCEP with ligand concentrations adjusted as described. **(A)** SAXS data were collected on ligand-free ATCase as it eluted from an in-line size-exclusion column, shown here as integrated intensity per frame (*top left*). Consistent with changes in radius of gyration ($R_g$) across the elution, SVD identified three significant singular values (*top right*). Concentration profiles (*bottom left*) and corresponding scattering profiles (*bottom right*) were extracted via REGALS,[8] a model-free decomposition method, for three components: a minor aggregate species (*gray*), a dominant dodecameric species (*blue*), and a changing background (*red*). $\chi^2$ values were ~1 across all frames, indicating excellent agreement between the data and REGALS model (*middle right*). *Bottom-right inset:* Guinier analysis yielded an $R_g$ of 48.2 ± 0.2 Å for the dominant species, corresponding to the ligand-free T-state. **(B)** The same experiment as in panel A was performed for ATCase in the presence of 500 µM CP, 10 mM succinate (*top left*). SVD identified two significant singular values (*top right*). Concentration (*bottom left*) and scattering profiles (*bottom right*) were extracted for a minor aggregate species (*gray*) and a dominant dodecameric species (*blue*). $\chi^2$ values were ~1 across all frames (*middle right*). *Bottom-right inset:* Guinier analysis yielded an $R_g$ of 49.7 ± 2.6 Å for the dominant species, corresponding to the nucleotide-free R-state. Data parameters detailed in Supplementary Table 4.

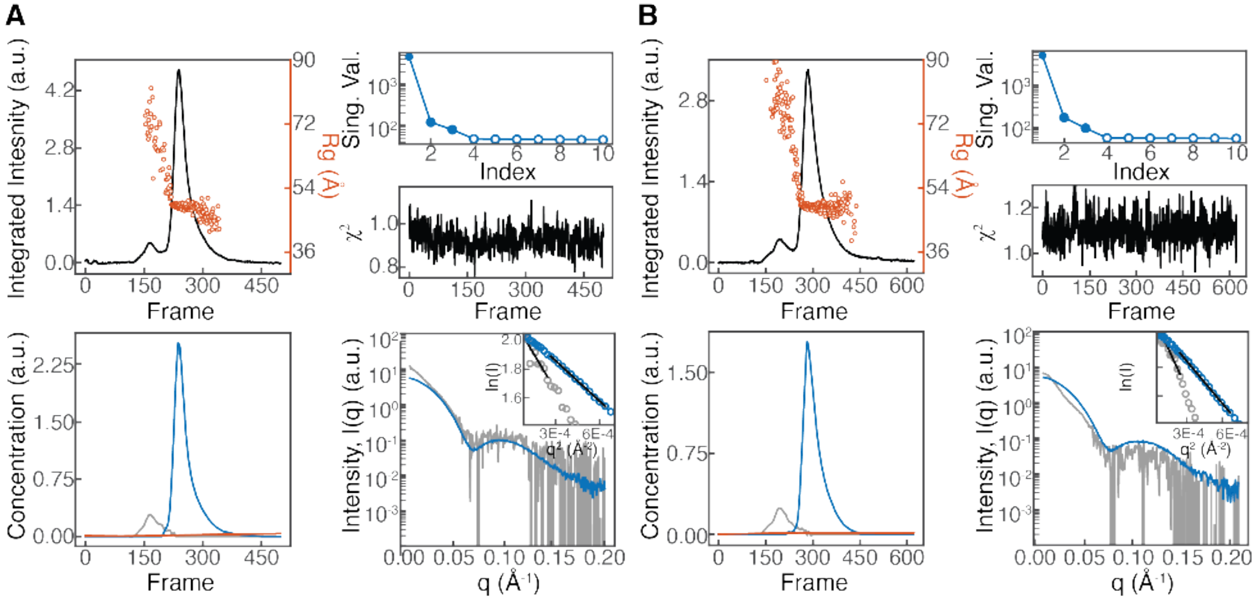


**Supplementary Figure 5. SAXS with in-line size exclusion chromatography yields high-quality scattering profiles of *E. coli* ATCase in the CTP- and CTP/UTP-bound R-states.**

Both the sample buffer and elution buffer were 40 mM Tris-HCl pH 7.5, 15 mM $MgCl_2$, 1 mM TCEP with ligand concentrations adjusted as described. **(A)** SEC-SAXS data collected on ATCase in the presence of 1.5 mM CTP, 500 µM CP, 10 mM succinate. Consistent with changes in $R_g$ across the elution, SVD identified three significant singular values (*top right*). Concentration profiles (*bottom left*) and corresponding scattering profiles (*bottom right*) were extracted via REGALS,[8] a model-free decomposition method, for three components: a minor aggregate species (*gray*), a dominant dodecameric species (*blue*), and a changing background (*red*) via REGALS. $\chi^2$ values were ~1 across all frames, indicating excellent agreement between the data and REGALS model (*middle right*). *Bottom-right inset:* Guinier analysis yielded an $R_g$ of 49.1 ± 0.6 Å for the dominant species, corresponding to the CTP-bound R-state. **(B)** SEC-SAXS data collected on ATCase in the presence of 1.5 mM CTP, 1.5 mM UTP, 500 µM CP, 10 mM succinate (*top left*). Again, SVD yielded three significant singular values (*top right*), and concentration profiles (*bottom left*) and scattering profiles (*bottom right*) were extracted for a minor aggregate species (*gray*), a dominant dodecameric species (*blue*), and a changing background (*red*). $\chi^2$ values were ~1 across all frames (*middle right*). *Bottom-right inset:* Guinier analysis yielded an $R_g$ of 49.0 ± 0.5 Å for the dominant species, corresponding to the CTP/UTP-bound R-state. Data parameters detailed in Supplementary Table 5.

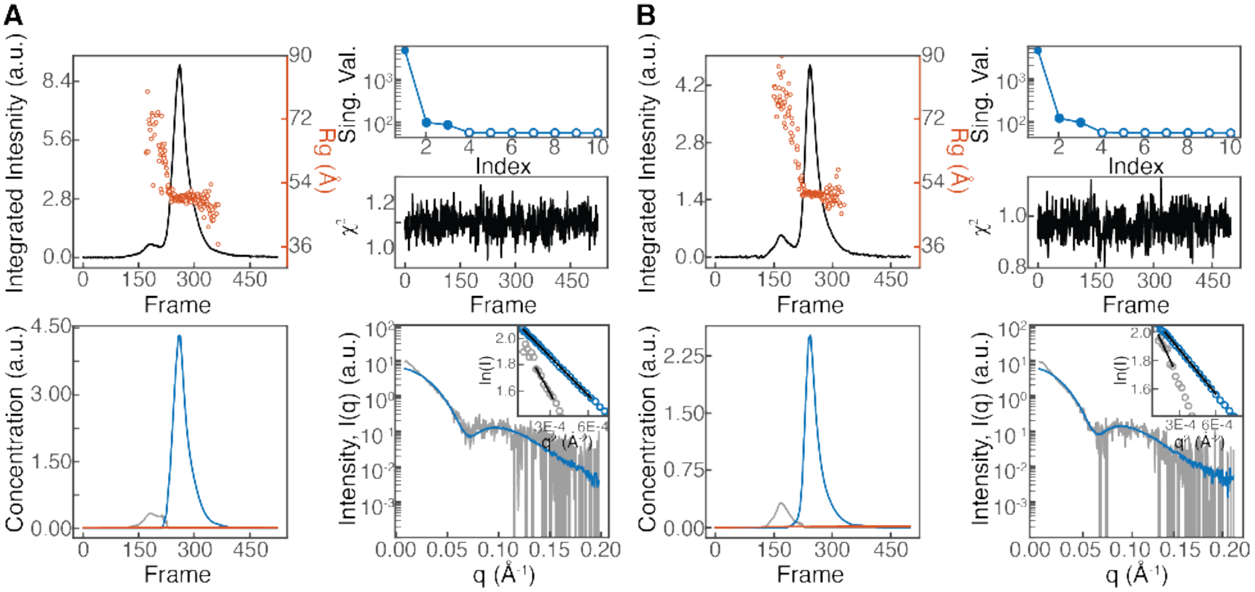


**Supplementary Figure 6. SAXS with in-line size exclusion chromatography yields high-quality scattering profiles of *E. coli* ATCase in the ATP- and ATP/GTP-bound R-states.**

Both the sample buffer and elution buffer were 40 mM Tris-HCl pH 7.5, 15 mM $MgCl_2$, 1 mM TCEP with ligand concentrations adjusted as described. **(A)** SEC-SAXS data collected on ATCase in the presence of 5 mM ATP, 500 µM CP, 10 mM succinate (*top left*). Consistent with changes in $R_g$ across the elution, SVD identified three significant singular values (*top right*). Concentration profiles (*bottom left*) and corresponding scattering profiles (*bottom right*) were extracted via REGALS,[8] a model-free decomposition method, for three components: a minor aggregate species (*gray*), a dominant dodecameric species (*blue*), and a changing background (*red*) via REGALS. $\chi^2$ values were ~1 across all frames, indicating good agreement between the data and REGALS model (*middle right*). *Bottom-right inset:* Guinier analysis yielded an $R_g$ of 49.9 ± 0.7 Å for the dominant species, corresponding to the ATP-bound R-state. **(B)** SEC-SAXS data collected on ATCase in the presence of 5 mM ATP, 1 mM GTP, 500 µM CP, 10 mM succinate (*top left*). Again, SVD yielded three significant singular values (*top right*), and concentration profiles (*bottom left*) and scattering profiles (*bottom right*) were extracted for a minor aggregate species (*gray*), a dominant dodecameric species (*blue*), and a changing background (*red*). $\chi^2$ values were ~1 across all frames (*middle right*). *Bottom-right inset:* Guinier analysis yielded an $R_g$ of 51.2 ± 0.7 Å for the dominant species, corresponding to the ATP/GTP-bound R-state. Data parameters detailed in Supplementary Table 6.

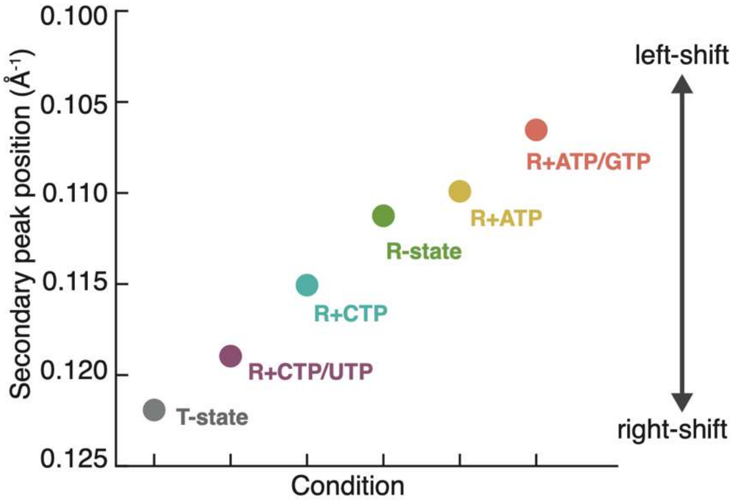


**Supplementary Figure 7. SAXS profiles show peak shifts consistent with a flexible R-state that expands in the presence of purines and contracts with pyrimidines.**

Shown are the positions of the secondary Kratky peaks in Figure 2A. A decrease in the peak position in reciprocal space corresponds to an expansion in real space. The T-state was obtained from ligand-free ATCase (*gray*). R-state conditions included 500 µM CP, 10 mM succinate and the following nucleotide compositions: 1.5 mM CTP/UTP (*purple*), 1.5 mM CTP alone (*blue*), no nucleotides (*green*), 5 mM ATP (*yellow*), or 5 mM ATP + 1 mM GTP (*salmon*). The pyrimidine pair CTP/UTP and purine pair ATP/GTP exert the strongest effects.

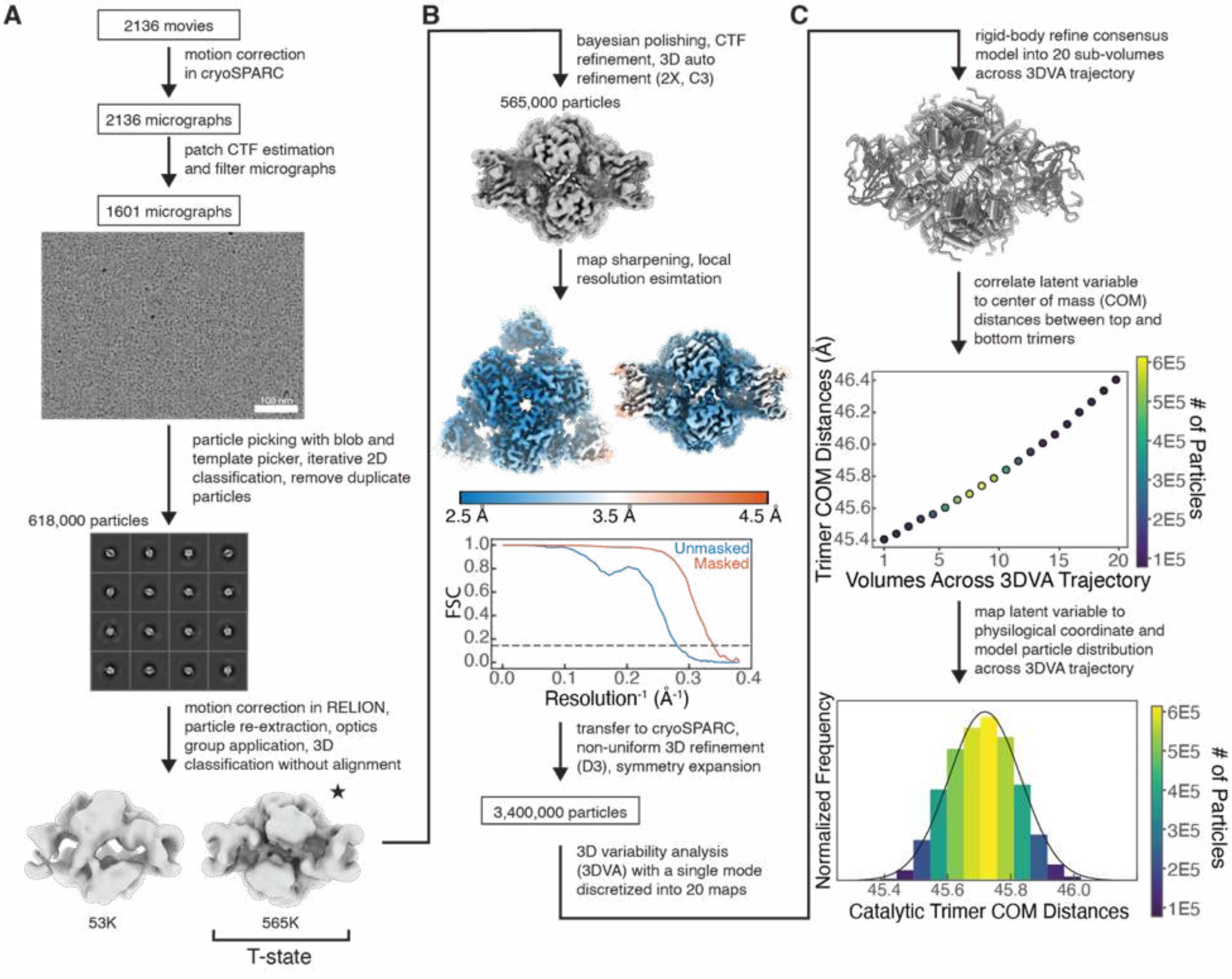


**Supplementary Figure 8. Cryo-EM imaging and processing pipeline of *E. coli* ATCase frozen with $Mg^{2+}$ but without ligands yields a map in the closed T-state conformation.**

**(A)** 2136 movies were imaged from ATCase frozen in buffer (40 mM Tris-HCl pH 7.5, 15 mM $MgCl_2$, 1 mM TCEP) with no ligands. Initial processing in cryoSPARC yielded a 618K particle stack, which was transferred to RELION. 3D classification separated a T-state-like class of 565K intact particles from a junk class. **(B)** Iterative 3D refinement with C3 symmetry, CTF parameter refinement, Bayesian polishing, and map sharpening yielded a 2.62-Å resolution map, as estimated at Fourier shell correlation (FSC) = 0.143. **(C)** The final polished particle stack was transferred to cryoSPARC and subjected to non-uniform refinement with D3 symmetry, followed by symmetry expansion. 3D Variability Analysis (3DVA) of the resulting particle stack indicated that the heterogeneity could be described by a single mode of breathing-like continuous motion. The 3DVA trajectory was discretized into 20 sub-volumes, and individual chains of the consensus model were rigid-body fit into each volume to convert the 3DVA trajectory coordinate to the center-of-mass (COM) distance between catalytic trimers. The resulting particle distribution was modeled as a normal distribution. Data processing and refinement statistics in Supplementary Table 7.

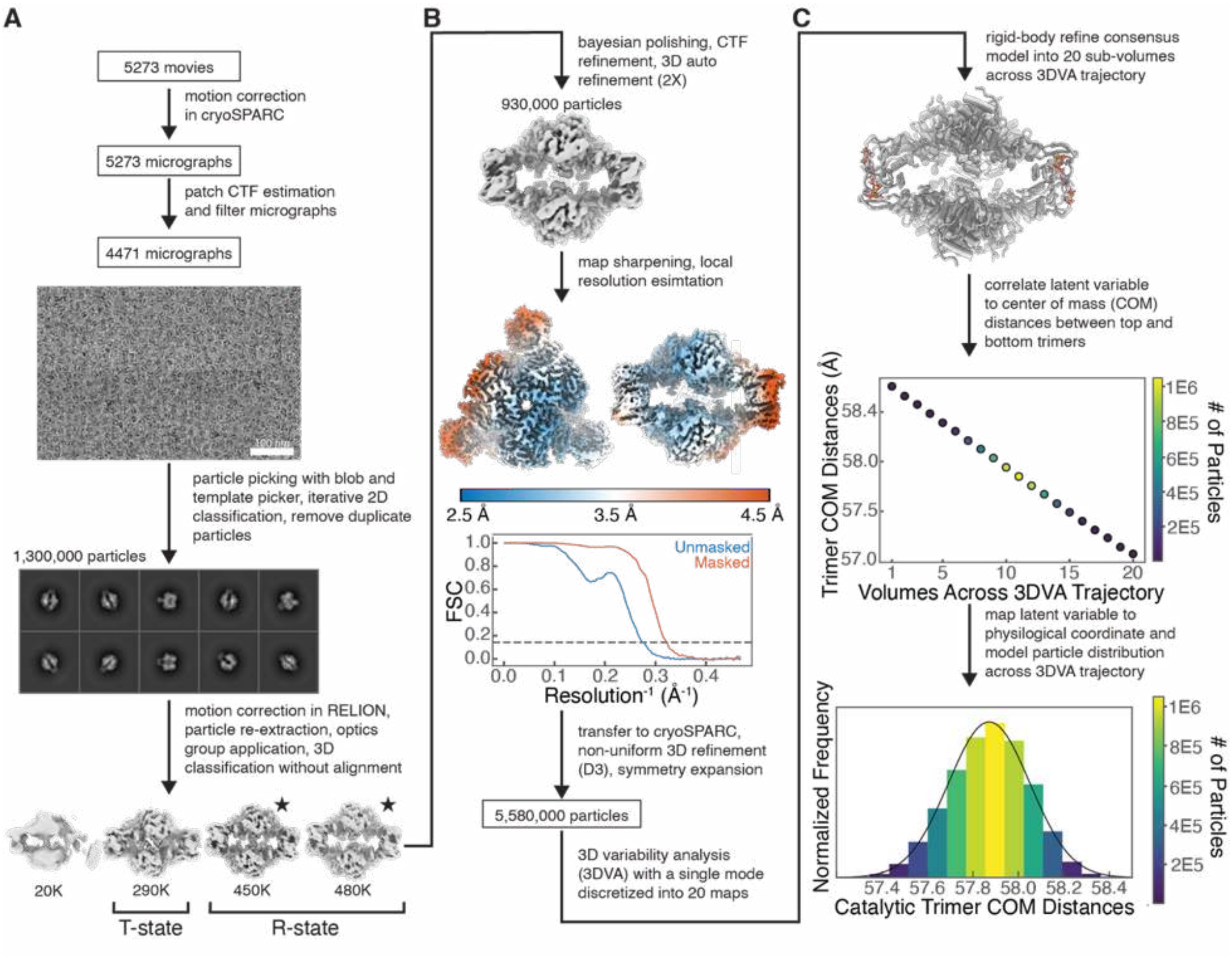


**Supplementary Figure 9. Cryo-EM imaging and processing pipeline of *E. coli* ATCase frozen with CP, succinate, CTP, UTP, and $Mg^{2+}$ yields maps representing the CTP/UTP-bound T- and R-states.**

**(A)** 5273 movies were imaged from ATCase frozen in buffer (40 mM Tris-HCl pH 7.5, 15 mM $MgCl_2$, 1 mM TCEP) supplemented with 1.5 mM CTP, 1.5 mM UTP, 500 µM CP, and 10 mM succinate. Initial processing in cryoSPARC yielded a 1.3M particle stack, which was transferred to RELION. 3D classification separated a T-state-like class of 290K intact particles and two R-state-like classes of 930K intact particles (*starred*) from a junk class. **(B)** Iterative 3D refinement with C1 symmetry, CTF parameter refinement, Bayesian polishing, and map sharpening yielded a 3.05-Å resolution map of ATCase bound to CTP/UTP, CP, and succinate, as estimated at FSC = 0.143. A similar workflow was applied to the T-state-like class to produce a 3.56-Å resolution C3 map of ATCase bound to CTP/UTP and CP only. **(C)** The final polished particle stack was transferred to cryoSPARC and subjected to non-uniform refinement with D3 symmetry, followed by symmetry expansion. 3DVA of the resulting particle stack indicated that the heterogeneity could be largely described by a single mode of breathing-like continuous motion. The 3DVA trajectory was discretized into 20 sub-volumes, and individual chains of the consensus model were rigid-body fit into each volume to convert the 3DVA trajectory coordinate to the COM distance between catalytic trimers. The resulting particle distribution was modeled as a normal distribution. Data processing and refinement statistics in Supplementary Tables 7-8.

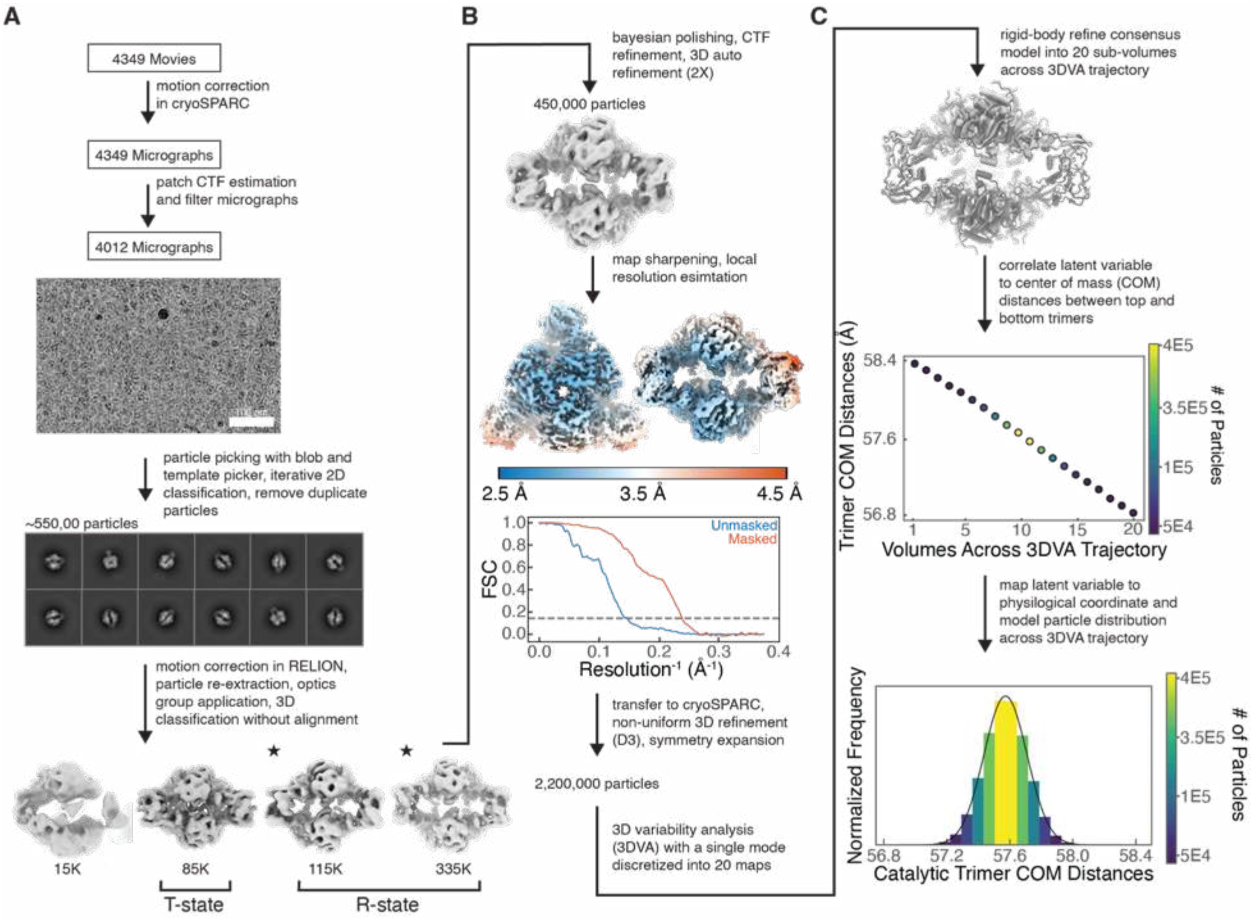


**Supplementary Figure 10. Cryo-EM imaging and processing pipeline of *E. coli* ATCase frozen with CP, succinate, CTP, and $Mg^{2+}$ yields maps representing the CTP-bound T- and R-states.**

**(A)** 4349 movies were imaged from ATCase frozen in buffer (40 mM Tris-HCl pH 7.5, 15 mM $MgCl_2$, 1 mM TCEP) supplemented with 1.5 mM CTP, 500 µM CP, and 10 mM succinate. Initial processing in cryoSPARC yielded a 550K particle stack, which was transferred to RELION. 3D classification separated a T-state-like class of 85K intact particles and two R-state-like classes of 450K intact particles (*starred*) from a junk class. **(B)** Iterative 3D refinement with C1 symmetry, CTF parameter refinement, Bayesian polishing, and map sharpening yielded a 3.53-Å resolution map of ATCase bound to CTP/CTP, CP, and succinate, as estimated at FSC = 0.143. A similar workflow was applied to the T-state-like class to produce a 3.85-Å resolution C3 map of ATCase bound to CTP/CTP and CP only. **(C)** The final polished particle stack was transferred to cryoSPARC and subjected to non-uniform refinement with D3 symmetry, followed by symmetry expansion. 3DVA of the resulting particle stack indicated that the heterogeneity could be largely described by a single mode of breathing-like continuous motion. The 3DVA trajectory was discretized into 20 sub-volumes, and individual chains of the consensus model were rigid-body fit into each volume to convert the 3DVA trajectory coordinate to the COM distance between catalytic trimers. The resulting particle distribution was modeled as a normal distribution. Data processing and refinement statistics in Supplementary Tables 7-8.

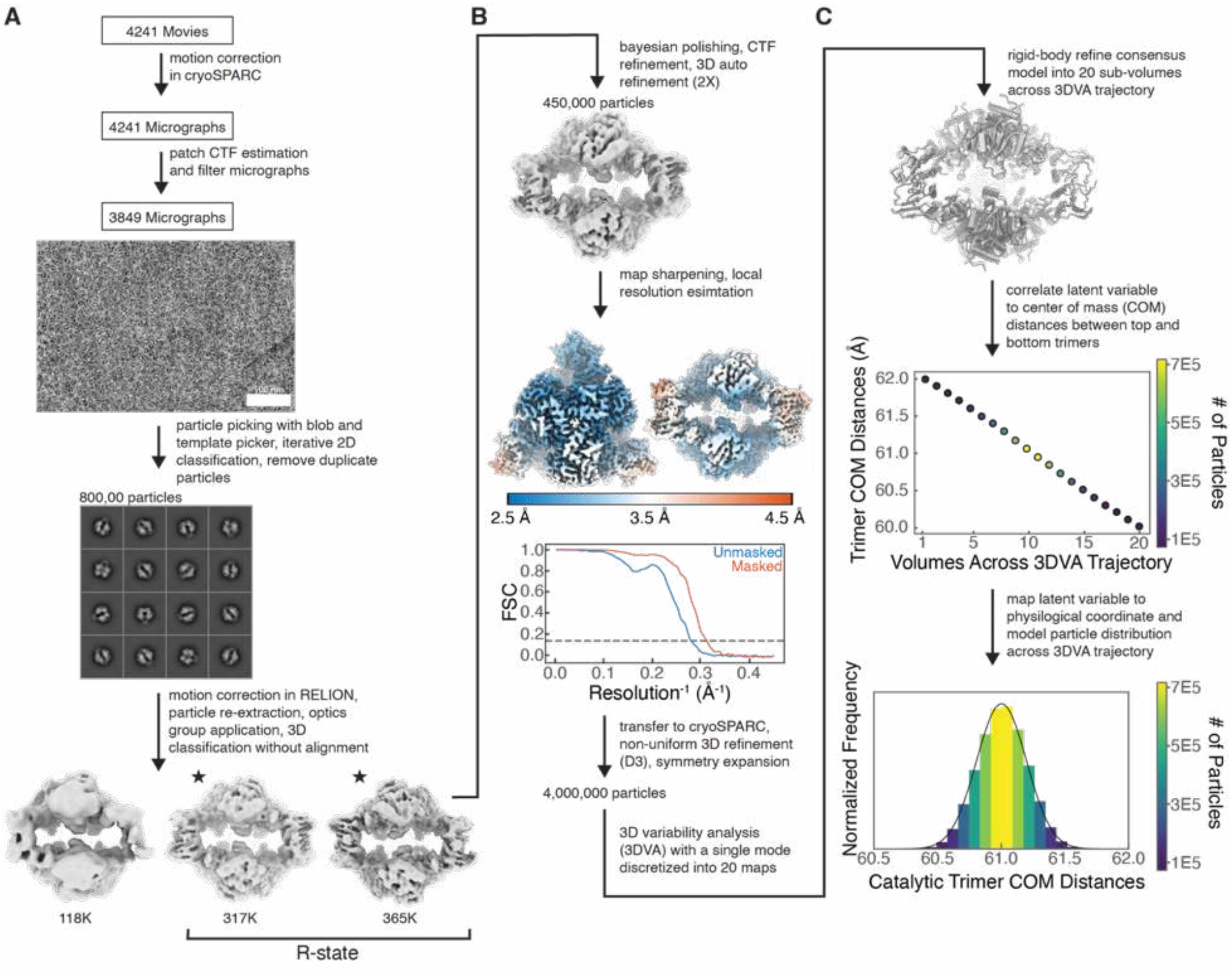


**Supplementary Figure 11. Cryo-EM imaging and processing pipeline of *E. coli* ATCase frozen with CP, succinate, and $Mg^{2+}$ yields a map representing the nucleotide-free R-state.**

**(A)** 4241 movies were imaged from ATCase frozen in buffer (40 mM Tris-HCl pH 7.5, 15 mM $MgCl_2$, 1 mM TCEP) supplemented with 500 µM CP and 10 mM succinate. Initial processing in cryoSPARC yielded an 800K particle stack, which was transferred to RELION. 3D classification separated two R-state-like classes of 682K intact particles from a junk class. **(B)** Iterative 3D refinement with C1 symmetry, CTF parameter refinement, Bayesian polishing, and map sharpening was performed to produce a 3.08-Å resolution map of ATCase bound to CP and succinate, as estimated at FSC = 0.143. **(C)** The final polished particle stack was transferred to cryoSPARC and subjected to non-uniform refinement with D3 symmetry, followed by symmetry expansion. 3DVA of the resulting particle stack indicated that the heterogeneity could be largely described by a single mode of breathing-like continuous motion. The 3DVA trajectory was discretized into 20 sub-volumes, and individual chains of the consensus model were rigid-body fit into each volume to convert the 3DVA trajectory coordinate to the COM distance between catalytic trimers. The resulting particle distribution was modeled as a normal distribution. Data processing and refinement statistics in Supplementary Table 8.

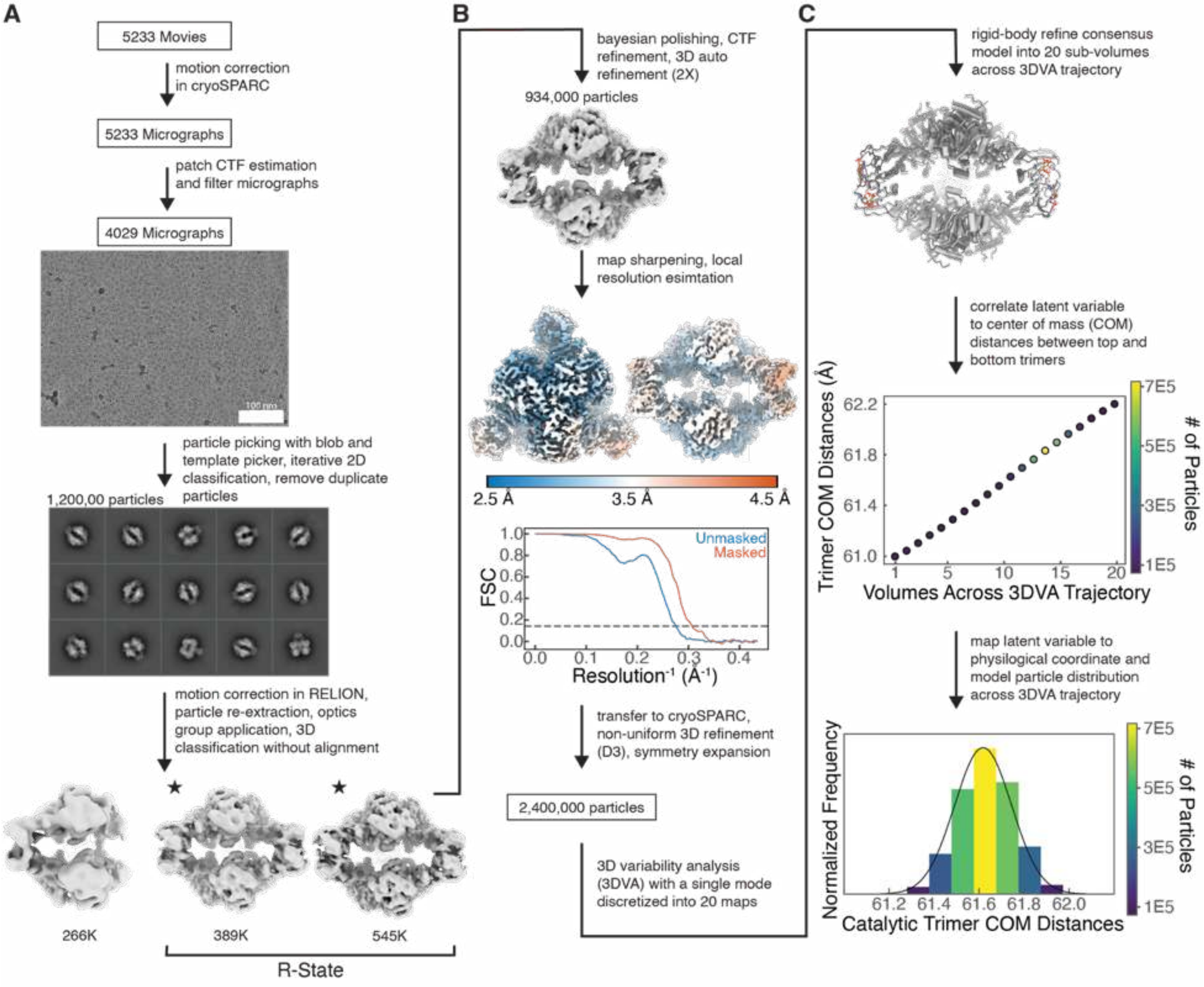


**Supplementary Figure 12. Cryo-EM imaging and processing pipeline of *E. coli* ATCase frozen with CP, succinate, ATP, and $Mg^{2+}$ yields a map representing the ATP-bound R-state.**

**(A)** 5233 movies were imaged from sample frozen in buffer (40 mM Tris-HCl pH 7.5, 15 mM $MgCl_2$, 1 mM TCEP) supplemented with 5 mM ATP, 500 µM CP, and 10 mM succinate. Initial processing in cryoSPARC yielded a 1.2M particle stack, which was transferred to RELION. 3D classification separated two R-state-like classes of 934K intact particles from a junk class. **(B)** Iterative 3D refinement with C1 symmetry, CTF parameter refinement, Bayesian polishing, and map sharpening was performed to produce a 3.27-Å resolution map of ATCase bound to ATP/ATP, CP and succinate, as estimated at FSC = 0.143. **(C)** The final polished particle stack was transferred to cryoSPARC and subjected to non-uniform refinement with D3 symmetry, followed by symmetry expansion. 3DVA of the resulting particle stack indicated that the heterogeneity could be largely described by a single mode of breathing-like continuous motion. The 3DVA trajectory was discretized into 20 sub-volumes, and individual chains of the consensus model were rigid-body fit into each volume to convert the 3DVA trajectory coordinate to the COM distance between catalytic trimers. The resulting particle distribution was modeled as a normal distribution. Data processing and refinement statistics in Supplementary Table 8.

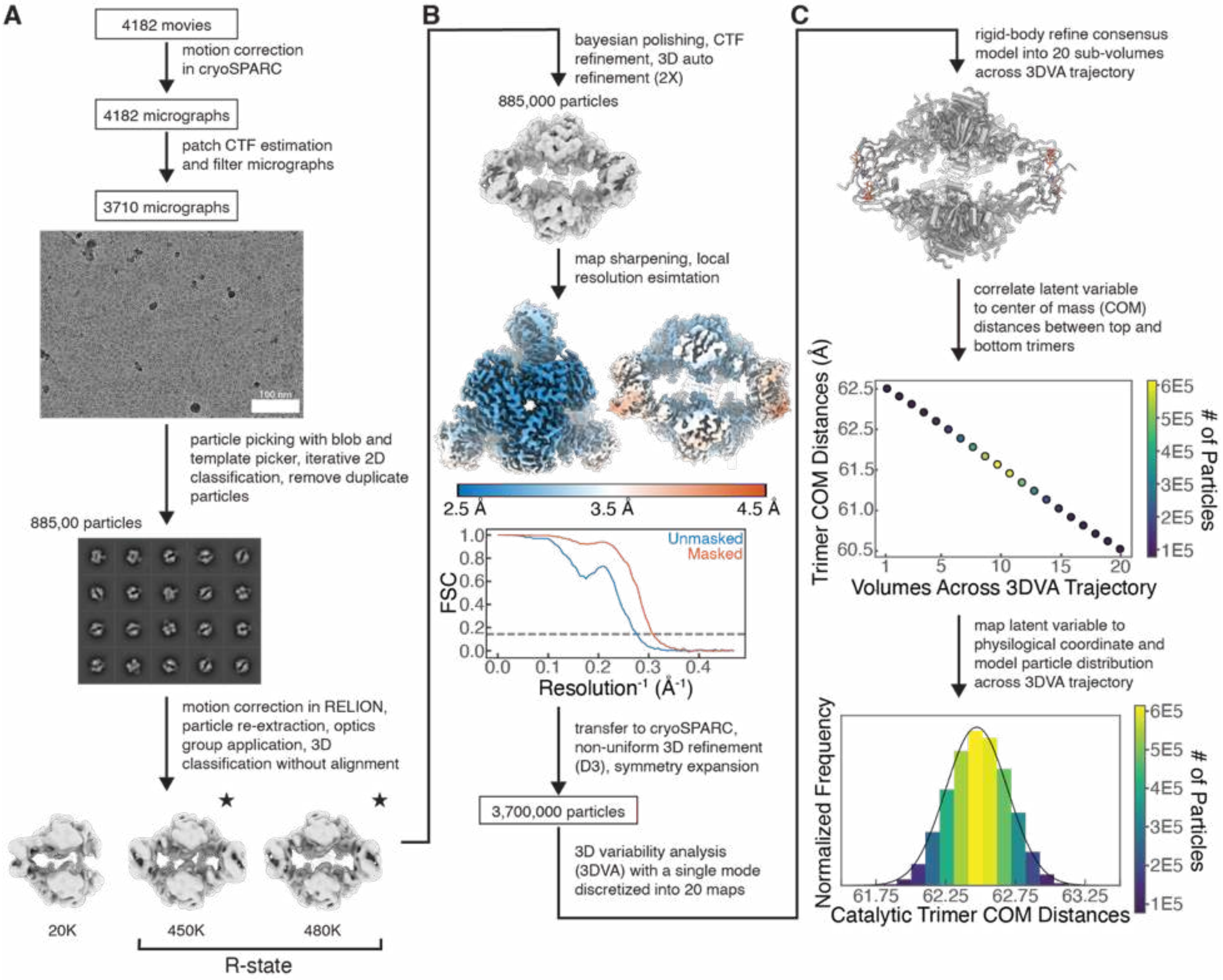


**Supplementary Figure 13. Cryo-EM imaging and processing pipeline of *E. coli* ATCase frozen with CP, succinate, ATP, GTP, and $Mg^{2+}$ yields a map representing the ATP/GTP-bound R-state.**

**(A)** 4182 movies were imaged from ATCase frozen in buffer (40 mM Tris-HCl pH 7.5, 15 mM $MgCl_2$, 1 mM TCEP) supplemented with 5 mM ATP, 1 mM GTP, 500 µM CP, and 10 mM succinate. Initial processing in cryoSPARC yielded an 885K particle stack, which was transferred to RELION. 3D classification separated two R-state-like classes of 614K intact particles from a junk class. **(B)** Iterative 3D refinement with C1 symmetry, CTF parameter refinement, Bayesian polishing, and map sharpening was performed to produce 3.27-Å resolution map of ATCase bound to ATP/GTP, CP and succinate, as estimated at FSC = 0.143. **(C)** The final polished particle stack was transferred to cryoSPARC and subjected to non-uniform refinement with D3 symmetry, followed by symmetry expansion. 3DVA of the resulting particle stack indicated that the heterogeneity could be largely described by a single mode of breathing-like continuous motion. The 3DVA trajectory was discretized into 20 sub-volumes, and individual chains of the consensus model were rigid-body fit into each volume to convert the 3DVA trajectory coordinate to the COM distance between catalytic trimers. The resulting particle distribution was modeled as a normal distribution. Data processing and refinement statistics in Supplementary Table 8.

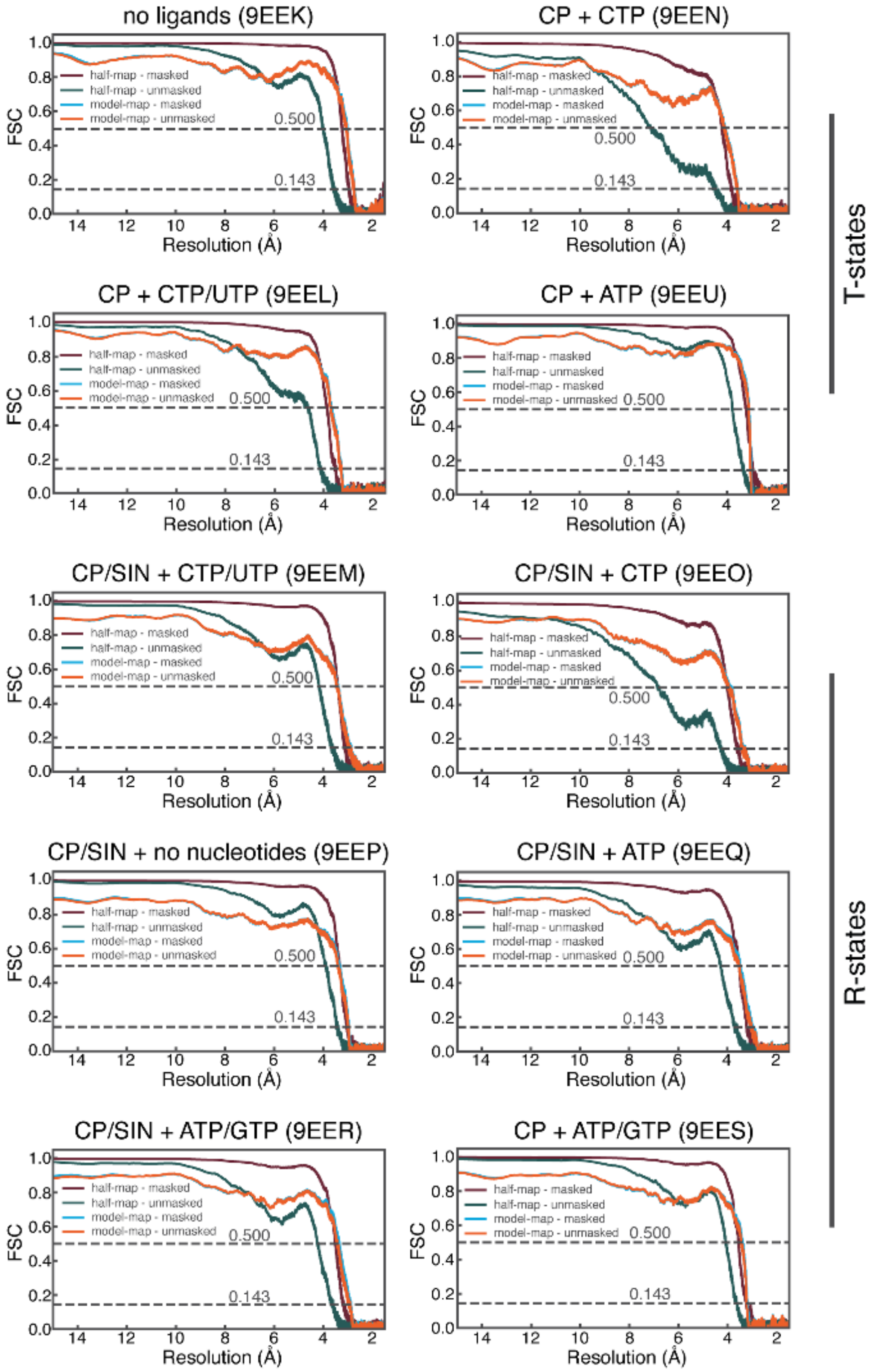


**Supplementary Figure 14. Resolution estimation by Fourier shell correlation (FSC) for all deposited cryo-EM models.**

Masked and unmasked gold-standard FSC curves (purple/green) and map–model FSC curves (light blue/orange) are shown for each cryo-EM model (9EEK–9EES, Supplementary Tables 7-8). FSCs were computed with phenix.mtriage using the sharpened full map, the corresponding half-maps, and the refined atomic model. Horizontal dashed lines indicate FSC = 0.143 and 0.5. For all reconstructions, the model-map FSC remains above 0.5 up to approximately the same resolution indicated by the masked half-map FSC, supporting the resolutions reported in Supplementary Tables 7-8.

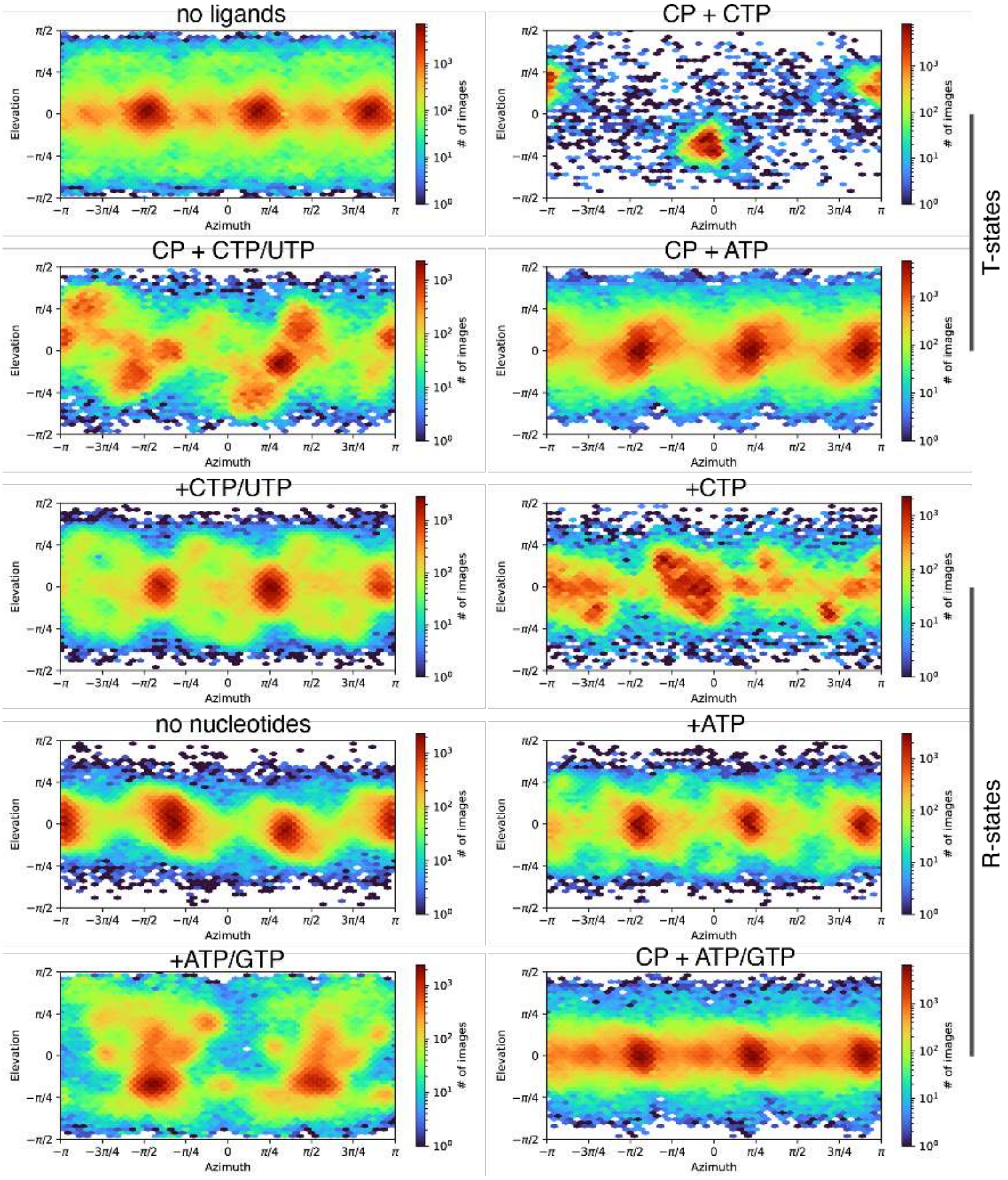


**Supplementary Figure 15. Angular distributions of particles used in the final cryo-EM reconstructions.**

Heat maps depict the distribution of particle orientations contributing to each final reconstruction, plotted by azimuth and elevation. The reconstruction of the T-state with CP + CTP was generated from a comparatively small particle stack with limited views. This conformation is not a primary focus of the present study but is included for completeness.

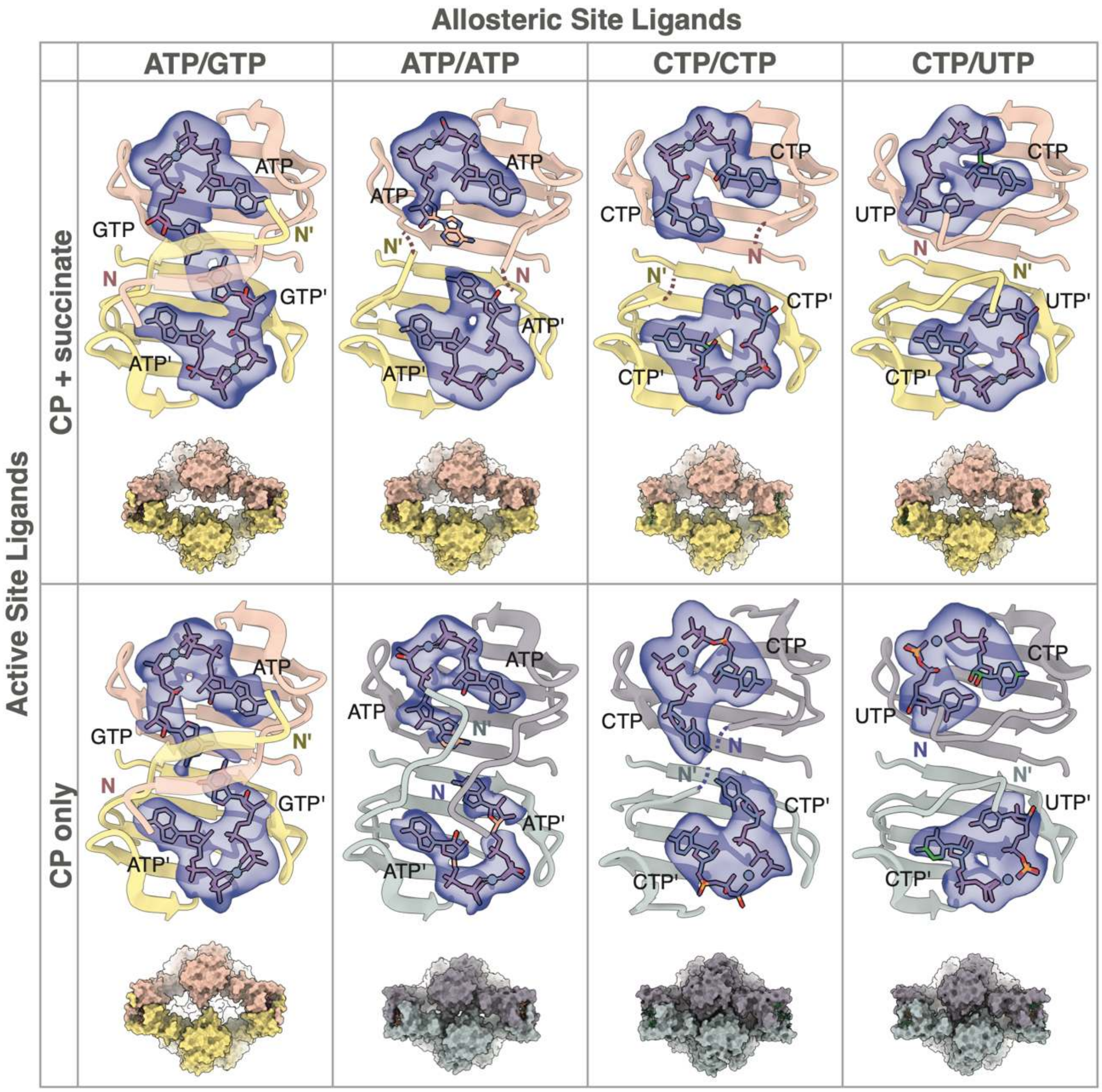


**Supplementary Figure 16. Cryo-EM density for nucleotides (*blue*) are consistent with the binding of two nucleotides per allosteric site, with the purine pair ATP/GTP and pyrimidine pair CTP/UTP producing the most ordered structures.**

T-states are shown in *blue/purple*, and R-states are shown in *yellow/pin*k. Columns from left to right: **(1)** Binding of ATP/GTP is characterized by strong density for both nucleotide-binding sites and a domain-swapped β-sheet formed by the regulatory N-termini. Two R-state maps with density for ATP/GTP were obtained, one with CP/succinate and another with CP only (Supplementary Fig. 27). **(2)** When only ATP is present, the density supports an ATP/ATP pair, but density is weak for the second site and disordered N-termini are observed reaching across the dimer interface. A cryo-EM sample with ATP, CP, and succinate yielded the R-state map, while the T-state map with CP alone was serendipitously obtained via 3D classification of another dataset (Supplementary Fig. 27B). **(3)** When only CTP is present, the density supports a CTP/CTP pair, but again the N-termini are disordered. Both T-state and R-state maps were obtained from the same dataset (Supplementary Fig. 17B). **(4)** Binding of CTP/UTP yields strong density for both nucleotide-binding sites with ordered N-termini forming specific contacts. Both T-state and R-state maps were obtained from the same dataset (Supplementary Fig. 17A).

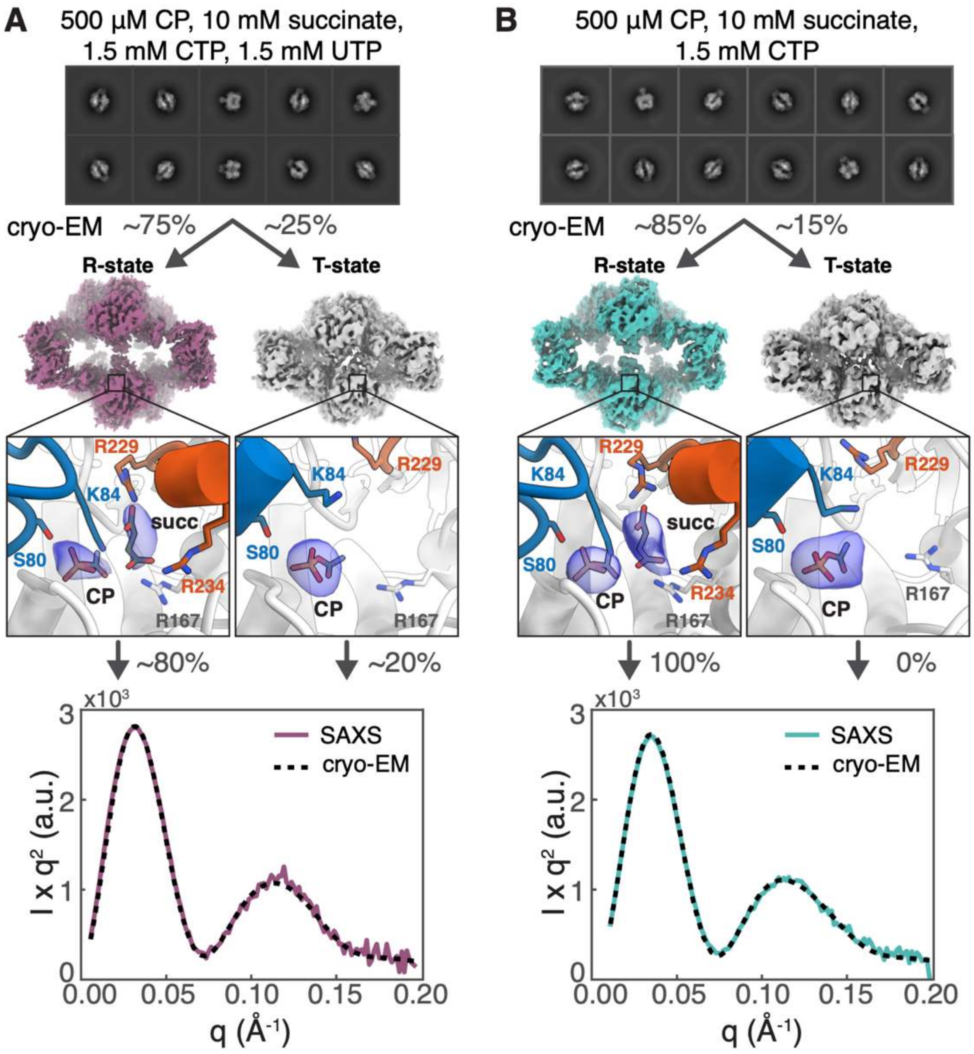


**Supplementary Figure 17. Cryo-EM combined with SAXS indicates that pyrimidines compress the R-state.**

**(A)** 3D classification of cryo-EM data obtained with 1.5 mM CTP/UTP, 500 µM CP, and 10 mM succinate yielded two maps from 1.22 million particles: ~75% refined to an open conformation with density for both CP and succinate, and the remaining ~25% to a closed conformation with density only for CP – consistent with the R-state and T-state, respectively. The SEC-SAXS profile obtained under the same condition (*solid*) is best fit by an 80:20 mixture of the refined R-state and T-state models (*dashed*). **(B)** Similarly, cryo-EM data collected with 1.5 mM CTP, 500 µM CP, and 10 mM succinate yielded two maps from 535,000 particles: ~85% refined to the R-state and the remaining ~15% to the T-state. The corresponding SEC-SAXS profile (*solid*) is best fit by the refined R-state model alone, with no contribution from the T-state model (*dashed*). Comparison with SAXS suggests that cryo-EM can overestimate the T-state population, possibly due to perturbations by the thin-layer grid environment. Nonetheless, the presence of T-state particles only in cryo-EM datasets obtained with pyrimidines and in SAXS data obtained with CTP/UTP support the interpretation that pyrimidines compress and destabilize the R-state, shifting the equilibrium toward the T-state.

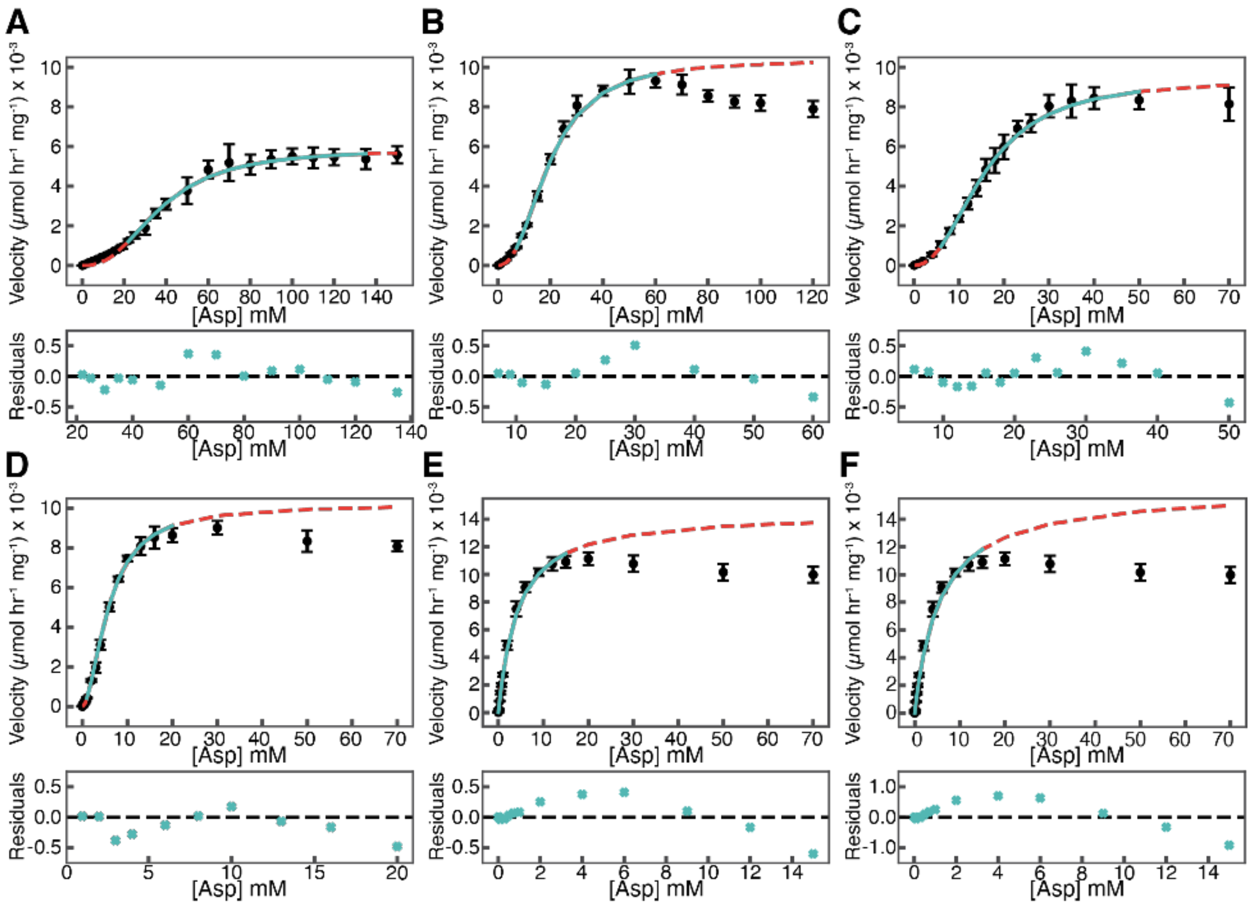


**Supplementary Figure 18. Nucleotides tune the cooperativity of *E. coli* ATCase, with the purine pair ATP/GTP and pyrimidine pair CTP/UTP producing the most pronounced effects.**

Saturation curves were obtained by measuring initial rates with 4.8 mM CP and different Asp concentrations. Model fitting (*red, dashed*) was performed within a truncated data frame (*cyan, solid*) to exclude regions affected by apparent substrate inhibition. **(A)** The saturation curve obtained with 1.5 mM CTP/UTP fit to the Hill equation yielded a Hill coefficient ($n_H$) of 2.9 and a $K_{1/2}$ of 37 mM. **(B)** With 1.5 mM CTP, the Hill fit gave $n_H \sim 2.6$ and $K_{1/2}$ ~18 mM. **(C)** With no nucleotides, the Hill fit gave $n_H \sim 2.5$ and $K_{1/2}$ ~15 mM. **(D)** With 5 mM ATP, the Hill fit gave $n_H \sim 2.1$ and $K_{1/2}$ ~5.6 mM. **(E)** With 5 mM ATP, 1 mM GTP, the Hill fit gave $n_H \sim 1.2$ and $K_{1/2}$ ~2.9 mM. **(F)** Michaelis-Menten fit to the same data shown in panel E. Notably, conditions with a single nucleotide (CTP or ATP alone) or no nucleotides produced similar maximum velocities ($V_{max}$) of ~9-10 mmol hr$^{-1}$ mg$^{-1}$. In contrast, the pyrimidine pair CTP/UTP had a strong inhibitory effect, lowering $V_{max}$ to 5.7 mmol hr$^{-1}$ mg$^{-1}$, while the purine pair ATP/GTP clearly activated the enzyme increasing $V_{max}$ to 12.6 mmol hr$^{-1}$ mg$^{-1}$. These nucleotide pairs also produced the most pronounced effects on enzyme cooperativity.

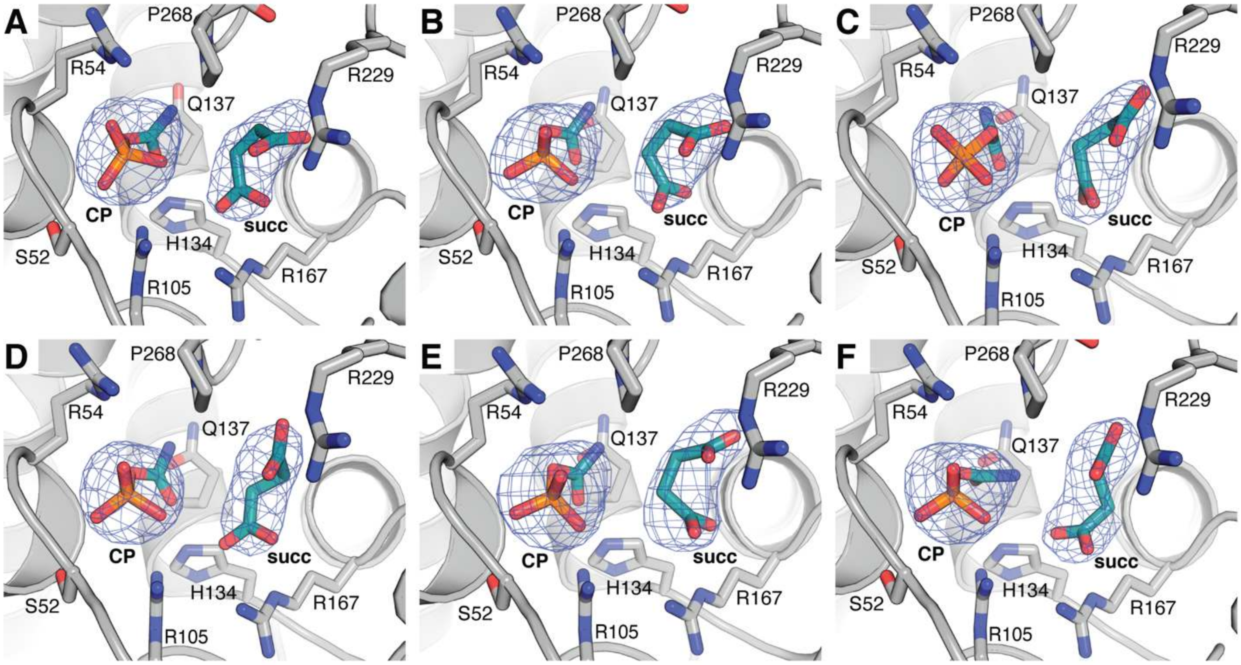


**Supplementary Figure 19. Electron density supports the presence of CP and succinate in the active sites of our $P2_12_12_1$ crystal structure of *E. coli* ATCase.**

ATCase was co-crystallized with CP, succinate, ATP, and $Mg^{2+}$ at near-neutral pH. In this crystal form, the asymmetric unit contains the entire enzyme, yielding six unique active sites. Shown in blue mesh are the $mF_O - DF_C$ Polder omit maps for CP and succinate, contoured at $6\sigma$, for **(A)** chain A, **(B)** chain B, **(C)** chain F, **(D)** chain C, **(E)** chain G, and **(F)** chain K. The conformation of the substrate CP is similar across all active sites, while that of succinate is more variable, consistent with the fact that it is an analog of the second substrate Asp.

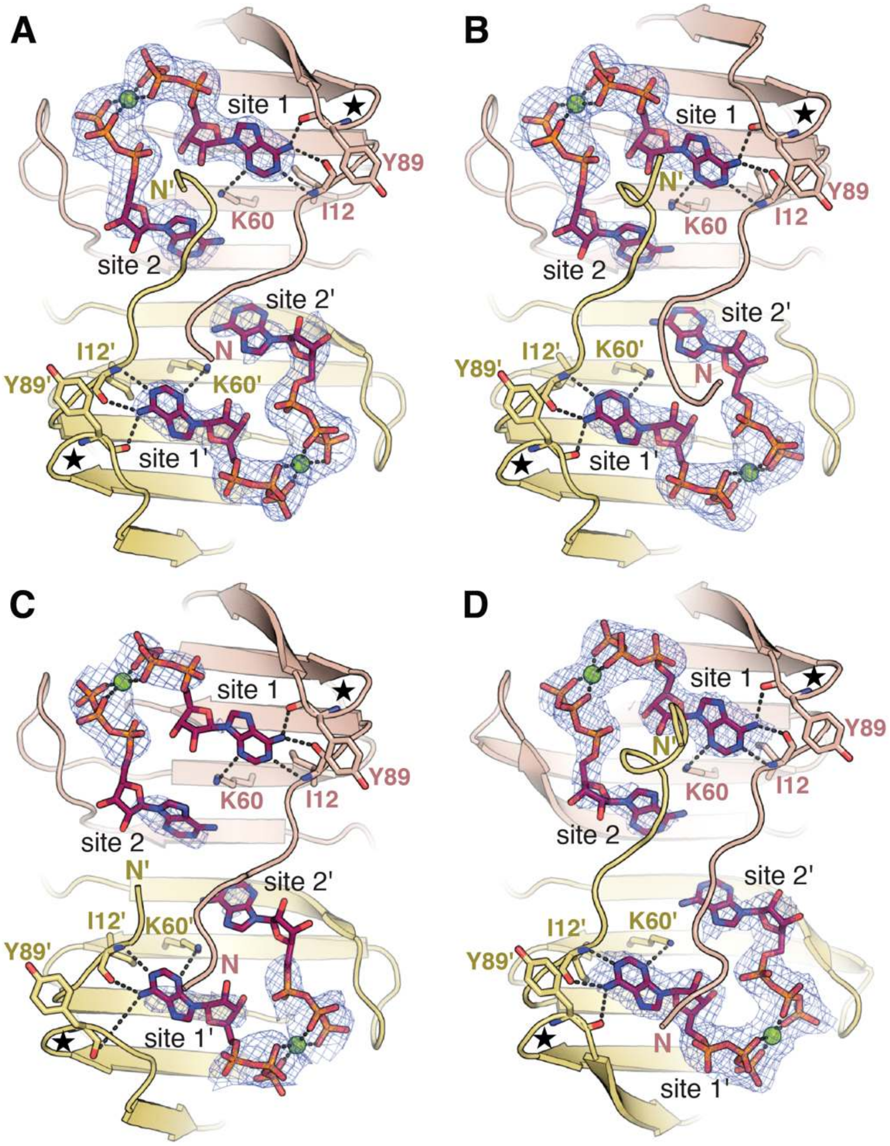


**Supplementary Figure 20. In crystal structures of *E. coli* ATCase obtained with $Mg^{2+}$, each allosteric site contains electron density for two ATP molecules, but significant disorder is observed.**

ATCase was co-crystallized with CP, succinate, ATP, and $Mg^{2+}$ at near-neutral pH. Shown in blue mesh are the $mF_O - DF_C$ Polder omit maps for the ATP molecules, contoured at $6\sigma$, in all three regulatory dimers: **(A)** chains E and I, **(B)** chains J and L, and **(C)** chains D and H. Disorder is most evident in for ATP in site 2. The quality and completeness of the electron density for the N-termini was variable across the dimers. Unlike the ATP/GTP-bound R-state structure, the N-termini reach across the dimer interface without forming any secondary structure. **(D)** Previous crystal structure of ATP-bound ATCase (PDB: 4kh0)[25] shows similar $2F_O - F_C$ electron density (contoured at $1\sigma$), indicating disorder in the N-termini and ATP molecules in site 2. In this *P*321 crystal form, there is only one unique dimer, and the N-terminus shown in yellow forms a lattice contact with a symmetry-related chain. In all panels, outer loop residues (88-89) are labeled with a star and $Mg^{2+}$ ions are shown as green spheres.

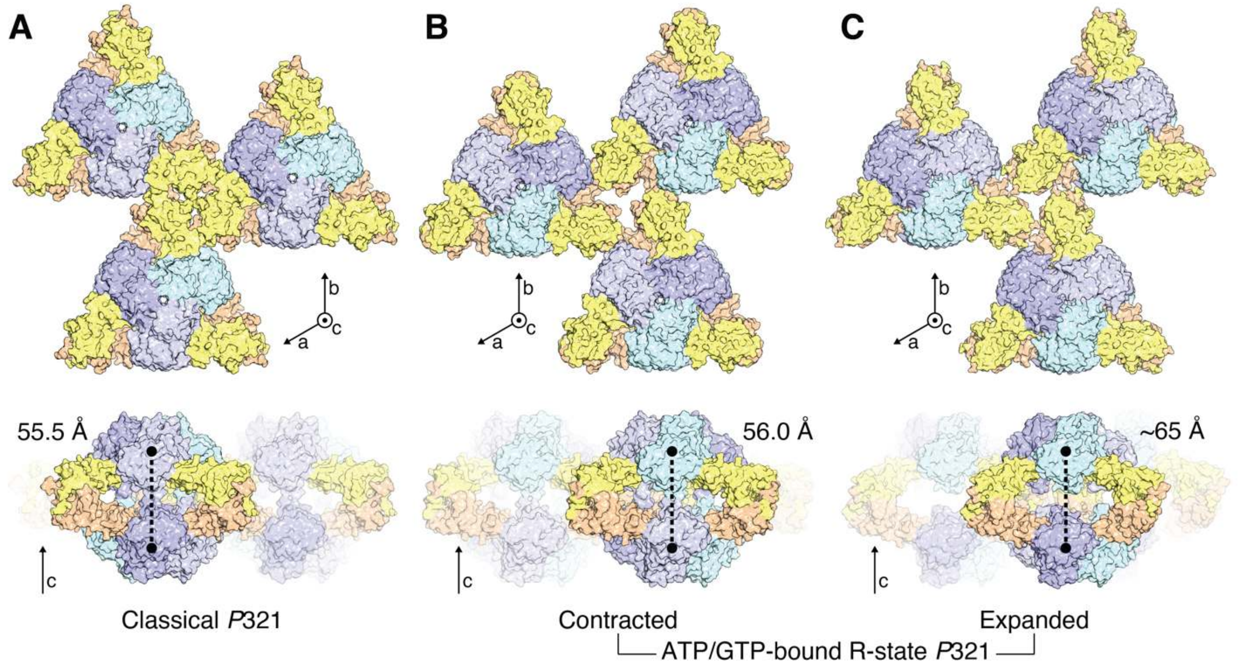


**Supplementary Figure 21: Co-crystallization of *E. coli* ATCase with ATP, GTP, PALA, and $Mg^{2+}$ at neutral pH produces a new crystal form.**

In all panels, the regulatory subunits are colored yellow and wheat, while catalytic subunits are colored in shades of blue and purple. **(A)** Classical *P*321 crystals in previous studies were grown in acidic conditions (pH 5.7-5.9), and nucleotide-bound structures with $Mg^{2+}$ were obtained by soaking crystals in nucleotide solutions following crystallization.[25] The center-of-mass distance between the trimers is 55.5 Å in the shown structure (PDB: 4kh1). **(B)** A new *P*321 crystal form was obtained at neutral pH that yielded the 2.2-Å crystal structure of ATP/GTP-bound R-state presented in this work (Supplementary Table 11). In this crystal form, the lattice is organized differently, placing the symmetry-related regulatory dimers further apart from each other. Although ATP/GTP are bound, the trimer distance is only 56.0 Å, similar to other R-state crystal structures. **(C)** The same crystallization drops that produced crystals described in panel B also produced crystals with an expanded **c**-axis dimension (Supplementary Table 12). These crystals did not diffract to high enough resolution for an unambiguous molecular replacement solution, but phasing was possible with various cryo-EM models of the R-state, reflecting the tendency for ATP/GTP to expand the enzyme.

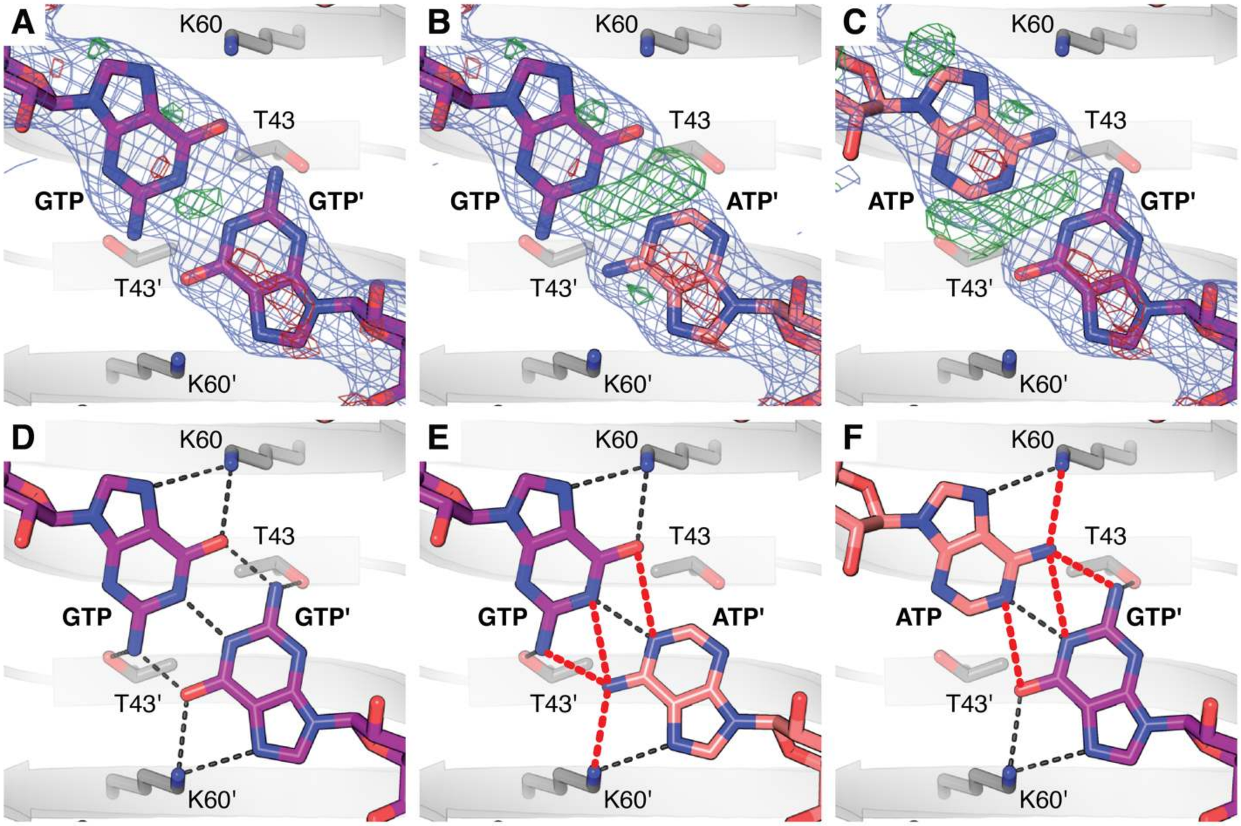


**Supplementary Figure 22. Electron density and ligand interactions observed in the crystal structure obtained with ATP, GTP, and PALA support GTP as the physiological partner for ATP.**

Zoomed-in view of the nucleotides bound near the regulatory dimer interface (see Figure 3B). $2F_O - F_C$ electron density is shown in blue at $1\sigma$, while positive and negative $F_O - F_C$ densities are shown in green and red, respectively, at ± 3 σ. **(A)** Modeling of a non-canonical GTP-GTP base-pairing interaction is supported by the $2F_O - F_C$ density and produces minimal difference density. **(B, C)** If either nucleotide is modeled as an ATP molecule, positive difference density is observed, supporting the presence of an amino group at the C2 position of the purine ring. **(D)** Assuming one of the GTP nucleobases is deprotonated at the N1 position, reasonable hydrogen-bonding interactions (*black dashed*) support the non-canonical base pairing. A domain-swapped interaction is also observed between each GTP nucleobase and T43 of the adjacent chain. **(E, F)** Modeling ATP in either site produces several clashes (*red dashed*), and the domain-swapped interaction with T43 is lost. The latter is also true with ATP modeled at both sites.

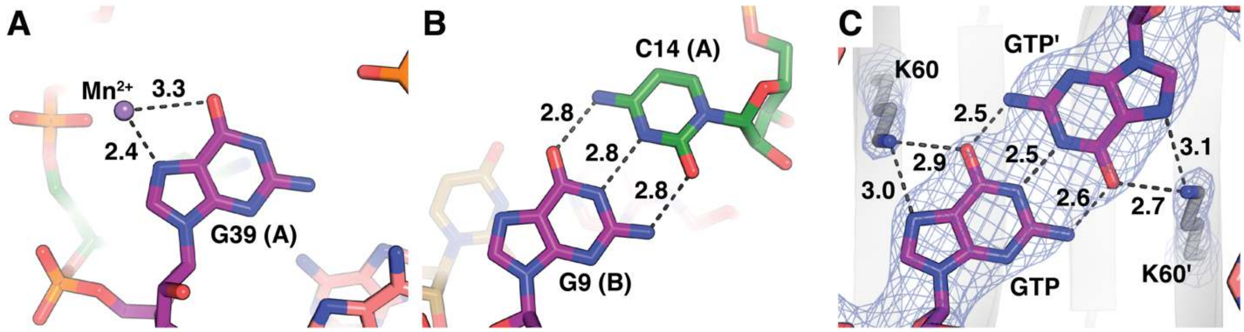


**Supplementary Figure 23. The allosteric site of *E. coli* ATCase supports non-canonical GTP base pairing by altering the p$K_a$ at the N1 position.**

**(A)** The hammerhead ribozyme provides a precedent for how a positive charge on the Hoogsteen face of a guanine can modulate the p$K_a$ at the N1 position. Shown is a crystal structure of the hammerhead ribozyme with a $Mn^{2+}$ ion bound to the Hoogsteen face of G39 (PDB: 5di2, chain A).[38] Electronic structure calculations based on this structure, with $Mg^{2+}$ in place of $Mn^{2+}$, suggests that the p$K_a$ is shifted from 9.2 (the value expected for free guanine) to 8.0, enabling G39 to act as a proton acceptor during catalysis.[39] **(B)** In a canonical C:G Watson-Crick base pair (also from PDB: 5di2), the N1 of guanine serves as a proton donor. **(C)** In our crystal structure of the ATP/GTP-bound R-state, GTP forms a non-canonical base pair, suggesting that only one GTP is protonated at the N1 position at a given time. A positive charge provided by a conserved K60 on the Hoogsteen face likely shifts the p$K_a$ to support this interaction.

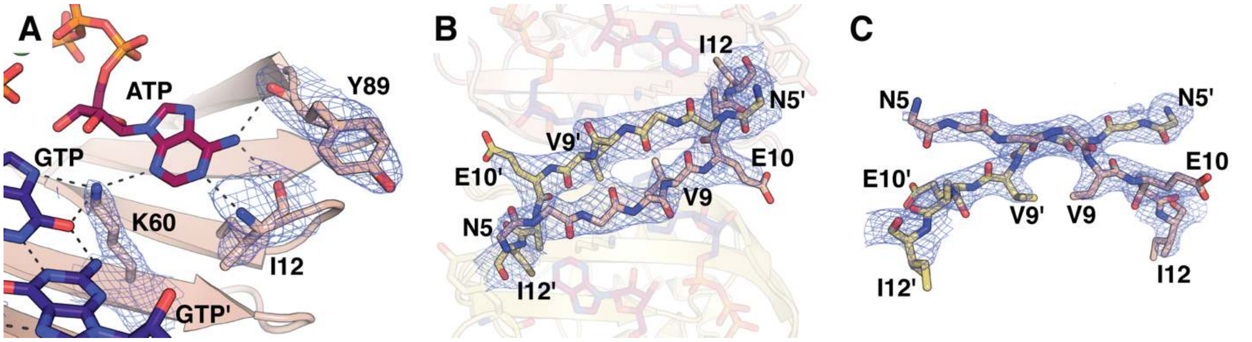


**Supplementary Figure 24. Interactions between key residues in the regulatory subunit and the ATP/GTP pair are supported by strong electron density.**

ATCase was co-crystallized with PALA, ATP, GTP and $Mg^{2+}$ at neutral pH. The $2F_O - F_C$ electron density (blue mesh, contoured at $1\sigma$) is shown for key interacting residues. **(A)** Consistent with previous crystal structures, I12 and Y89 bind ATP in site 1 (*black dashed lines*). In addition, our crystal structure reveals that K60 bridges ATP and GTP in site 2 (*black dashed lines*). **(B)** Top and **(C)** side views of the N-terminal residues 5–12 of the regulatory dimer. The electron density supports a domain-swapped N-terminal anti-parallel β-sheet not seen in previous crystal structures. Residue D4 and the sidechain atoms of residues N5, K6, L7, and Q8 are included in the model but not shown here due to limited electron density at $1\sigma$.

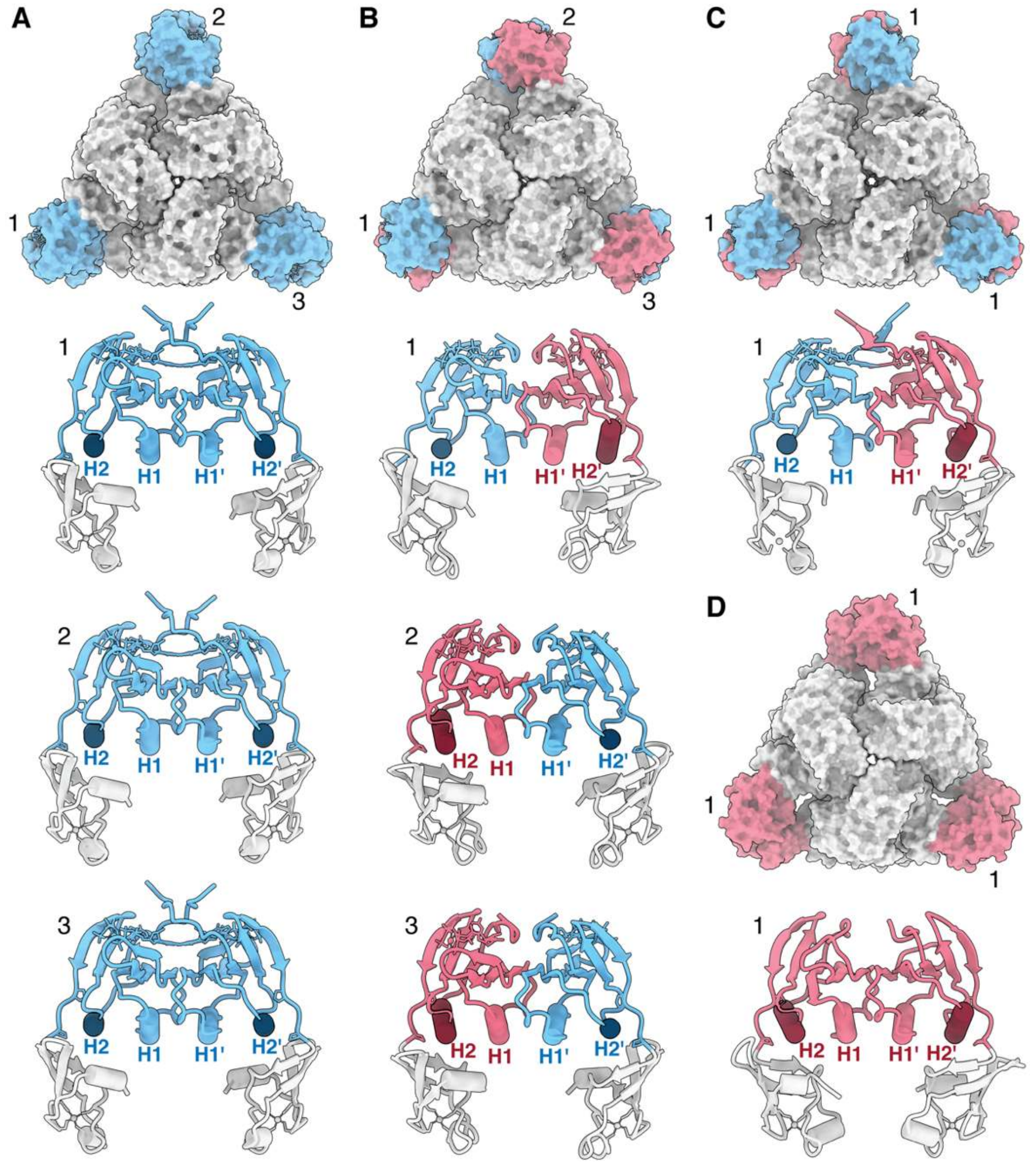


**Supplementary Figure 25. Cryo-EM reveals that pyrimidines produce a partially collapsed R-state, while purines produce a fully expanded R-state.**

Non-crystallographic symmetry (NCS) refinement was used to examine nucleotide-dependent changes in the internal structure and symmetry of *E. coli* ATCase. **(A)** For the ATP/GTP R-state, the nucleotide-binding domains in all three regulatory dimers (labeled 1-3) could be refined in a single NCS group (*blue*). In this fully expanded R-state conformation, both Zn domains (*gray cartoon*) are open, and the H2 and H2′ helices are straightened and parallel. **(B)** In contrast, the CTP/UTP R-state required two NCS groups to describe the asymmetry within each regulatory dimer. In each dimer, the Zn domain on one side is closed and the H2 helix is tucked at an angle (*red*), while the opposite side (*blue*) resembles the relaxed, open conformation seen in panel A. Inverted H2/H2′ asymmetry is observed in only one of the three dimers, resulting in an overall conformation that appears partially collapsed (like a wobbly table that leans to one side). **(C)** Crystal structures of the R-state also display H2/H2′ asymmetry like that observed in the EM model of the CTP/UTP R-state. However, in *P*321, there is only one unique regulatory dimer (labeled as 1). Shown here is PDB: 9eeh (from this study). **(D)** In both crystal structures and cryo-EM models of the T-state, both Zn domains are closed, and the H2 and H2′ helices are tucked at an angle. Shown here is PDB: 6at1,[24] a *P*321 crystal structure with one unique regulatory dimer (labeled as 1).

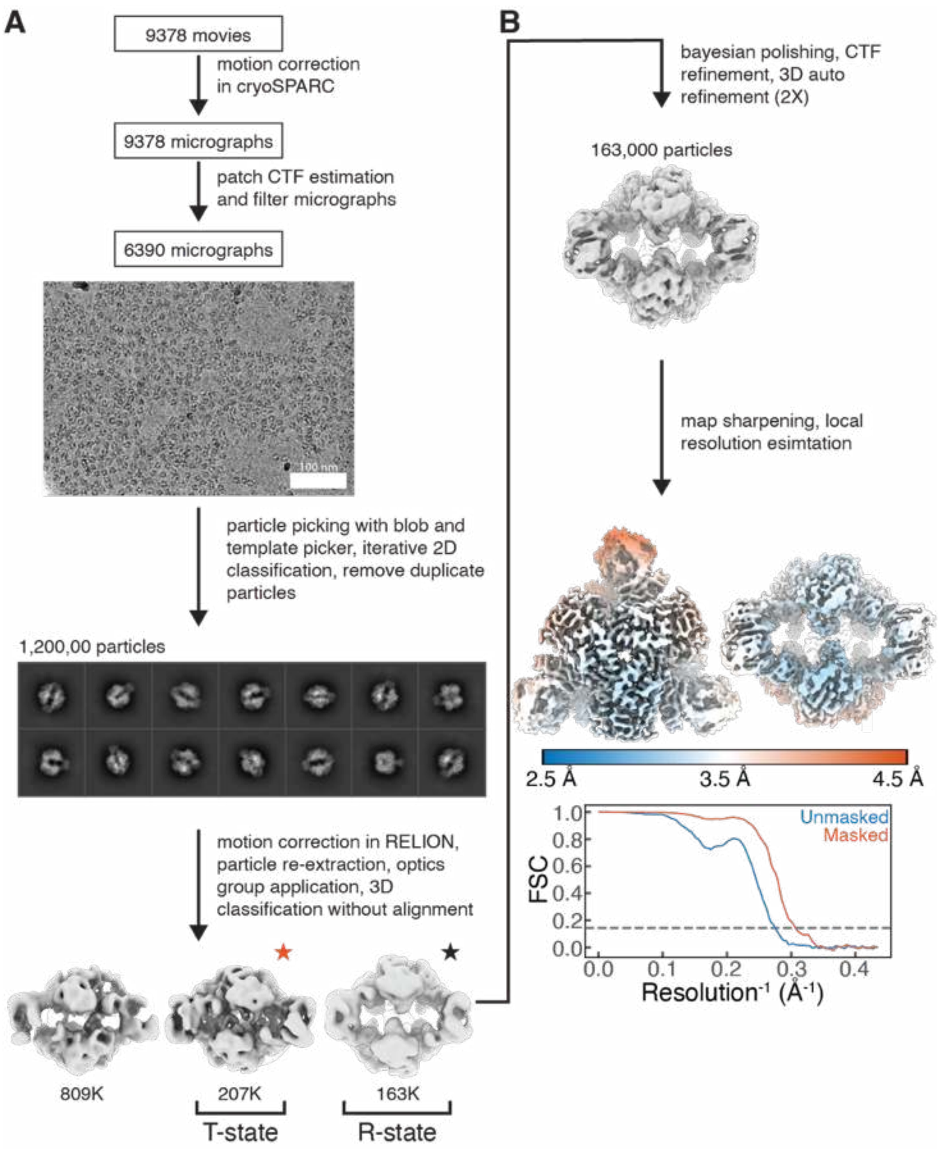


**Supplementary Figure 26. Cryo-EM imaging and processing pipeline of *E. coli* ATCase frozen with CP, ATP, GTP, and $Mg^{2+}$ yields maps representing the ATP/GTP-bound R-state and ATP/ATP-bound T-state.**

**(A)** 9378 movies were imaged from ATCase frozen in buffer (40 mM Tris-HCl pH 7.5, 15 mM $MgCl_2$, 1 mM TCEP) supplemented with 5 mM ATP, 1 mM GTP, and 500 µM CP. Initial processing in cryoSPARC yielded a 1.2M particle stack, which was transferred to RELION. 3D classification separated a T-state-like class of 207K intact particles (*red star*) and a R-state-like class of 163K intact particles (*black star*) from a junk class. **(B)** Iterative 3D refinement with C3 symmetry, CTF parameter refinement, Bayesian polishing, and map sharpening yielded a 3.27-Å resolution map of ATCase bound to ATP/GTP, and CP, as estimated at FSC = 0.143. A similar workflow was applied to the T-state-like class (*red star* in panel A) to produce a 2.97-Å resolution map of ATCase; unlike the R-state map, density at the nucleotide-binding sites was more consistent with two molecules of ATP bound (Supplementary Fig. 16). Data processing and refinement statistics in Supplementary Table 7.

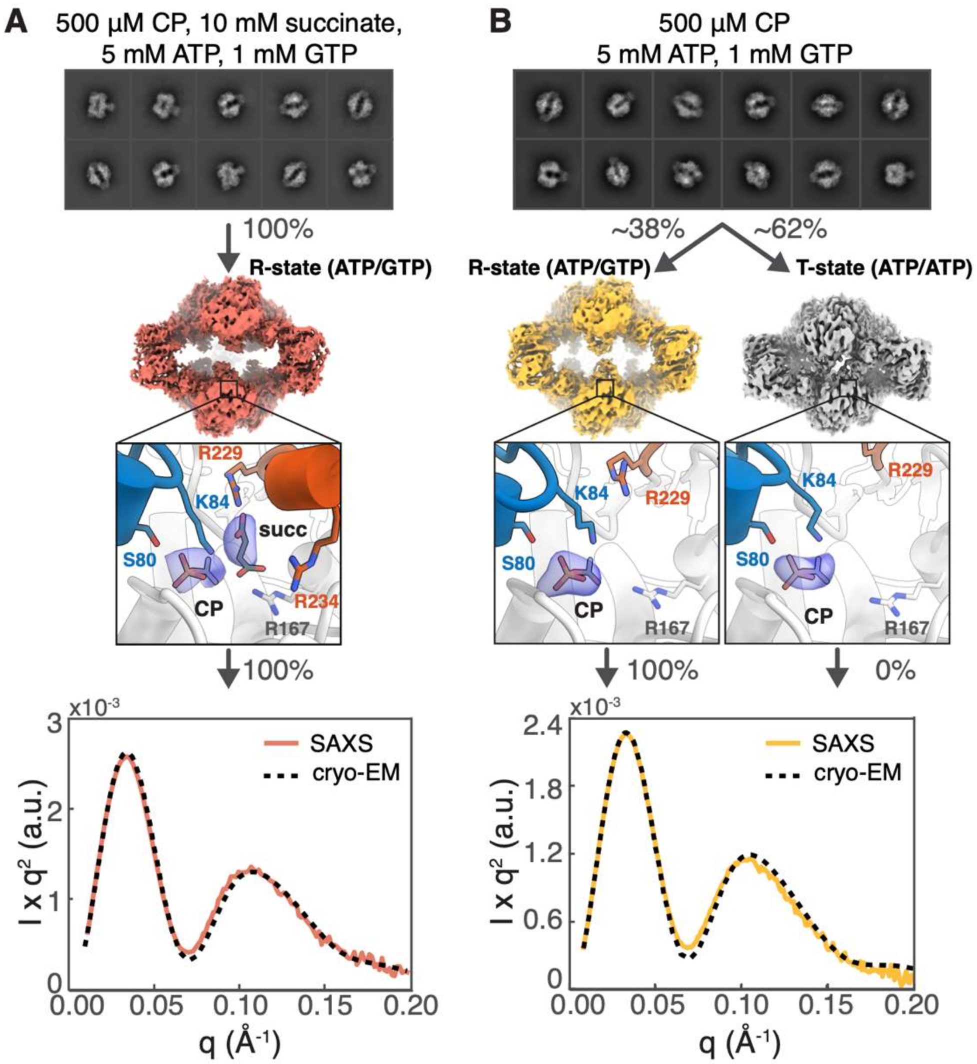


**Supplementary Figure 27. Cryo-EM combined with SAXS indicates that purines expand the R-state.**

**(A)** 3D classification of cryo-EM data obtained with 5 mM ATP, 1 mM GTP, 500 µM CP, and 10 mM succinate yielded a single map from 885,000 intact particles, which refined to an open R-state conformation with density for both CP and succinate. The SEC-SAXS profile obtained under the same condition (*solid*) aligns well with the refined R-state model (*dashed*). **(B)** In contrast, cryo-EM data collected with 5 mM ATP, 1 mM GTP, and 500 µM CP (no succinate) yielded two maps from 163,000 particles, both having density for CP only in the active sites: ~38% refined to an ATP/GTP-bound R-state conformation, while the remaining refined to an ATP/ATP-bound T-state conformation (see Supplementary Fig. 16). The corresponding SEC-SAXS profile (*solid*) is best fit by the refined R-state model alone, with no contribution from the T-state model (*dashed*). The differences in T- and R-state populations observed by cryo-EM and SAXS likely reflect differences in the sample environment: the thin layer environment of the cryo-EM grid appears to favor collapse into the T-state more than bulk solution. Nonetheless, we find that the purine pair ATP/GTP is the only nucleotide combination that can expand the ATCase from the closed T-state.

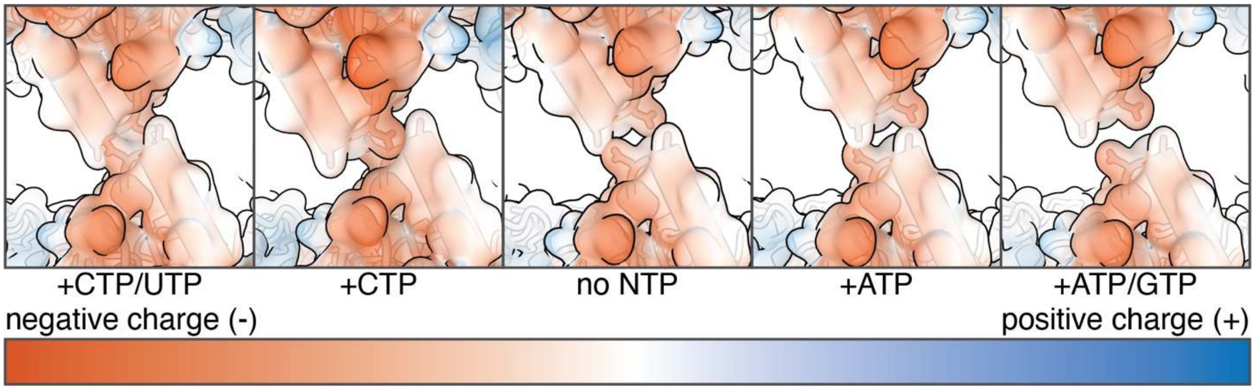


**Supplementary Figure 28. An extended 240s loop conformation is incompatible with all R-states except in the fully expanded conformation induced by the purine pair ATP/GTP.**

When only the first substrate (CP) is bound, the 240s loops of ATCase remains extended, as seen in the novel R-state conformation we observed by cryo-EM with ATP, GTP, and CP. To assess whether this extended 240s loop conformation is compatible with R-states induced by other nucleotide conditions, we rigid-body docked the catalytic trimers (chains B, A, F and C, K, G) into cryo-EM maps shown in Figure 2B. In all cases except ATP/GTP, a steric clash and an unfavorable electrostatic interaction are observed between opposing 240s loops. This finding explains why binding of both substrates (CP and Asp) is required to induce the R-state under all nucleotide conditions—except with ATP/GTP, where binding of CP alone is sufficient to transition to the R-state.

**Supplementary Figure 29. Synthetic route of PALA.**

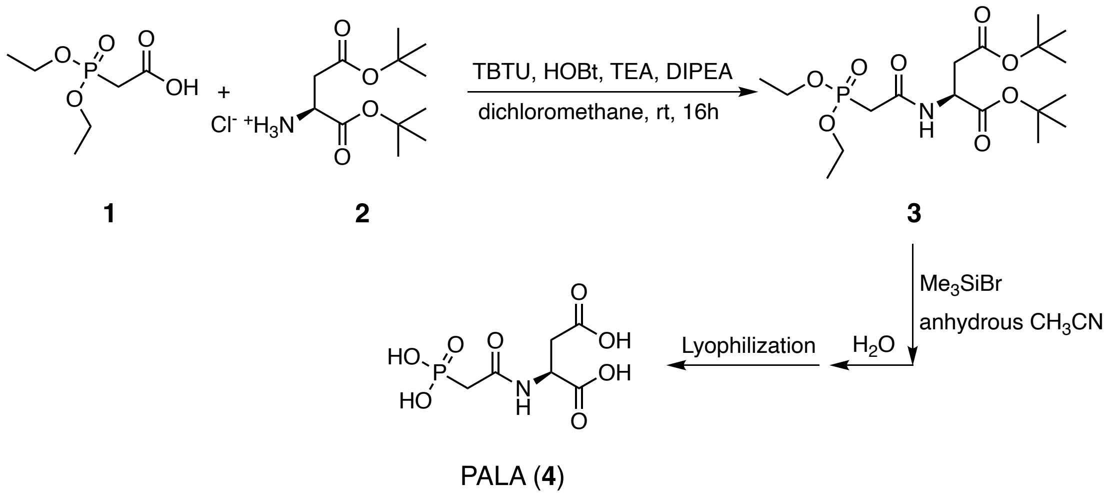

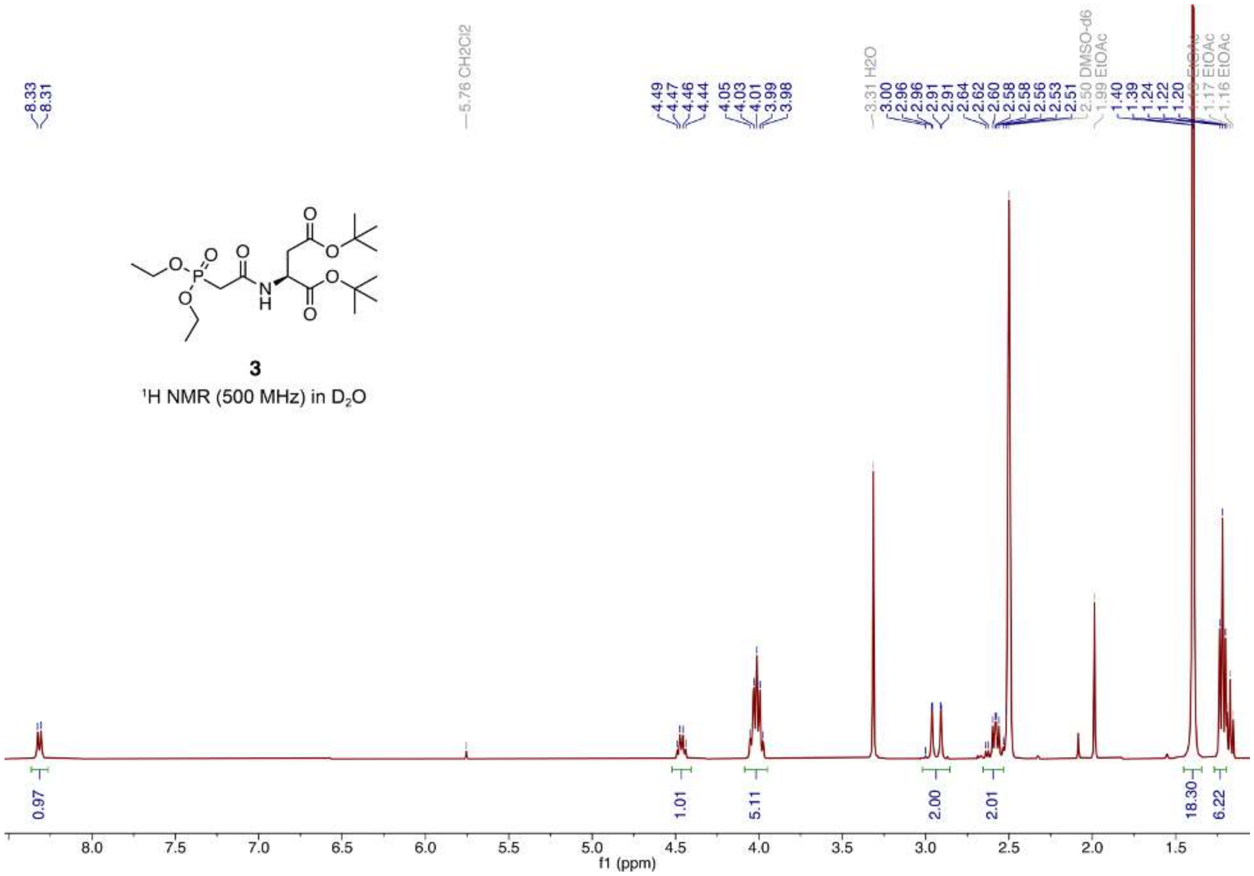


**Supplementary Figure 30. ¹H NMR spectrum of intermediate precursor to PALA in DMSO.**

¹H NMR spectrum of compound **3** in Supplementary Scheme 1. Signals at ~4.01 ppm and ~1.24 ppm fully or partially overlap with the signals from residual solvent (EtOAc, ethyl acetate).

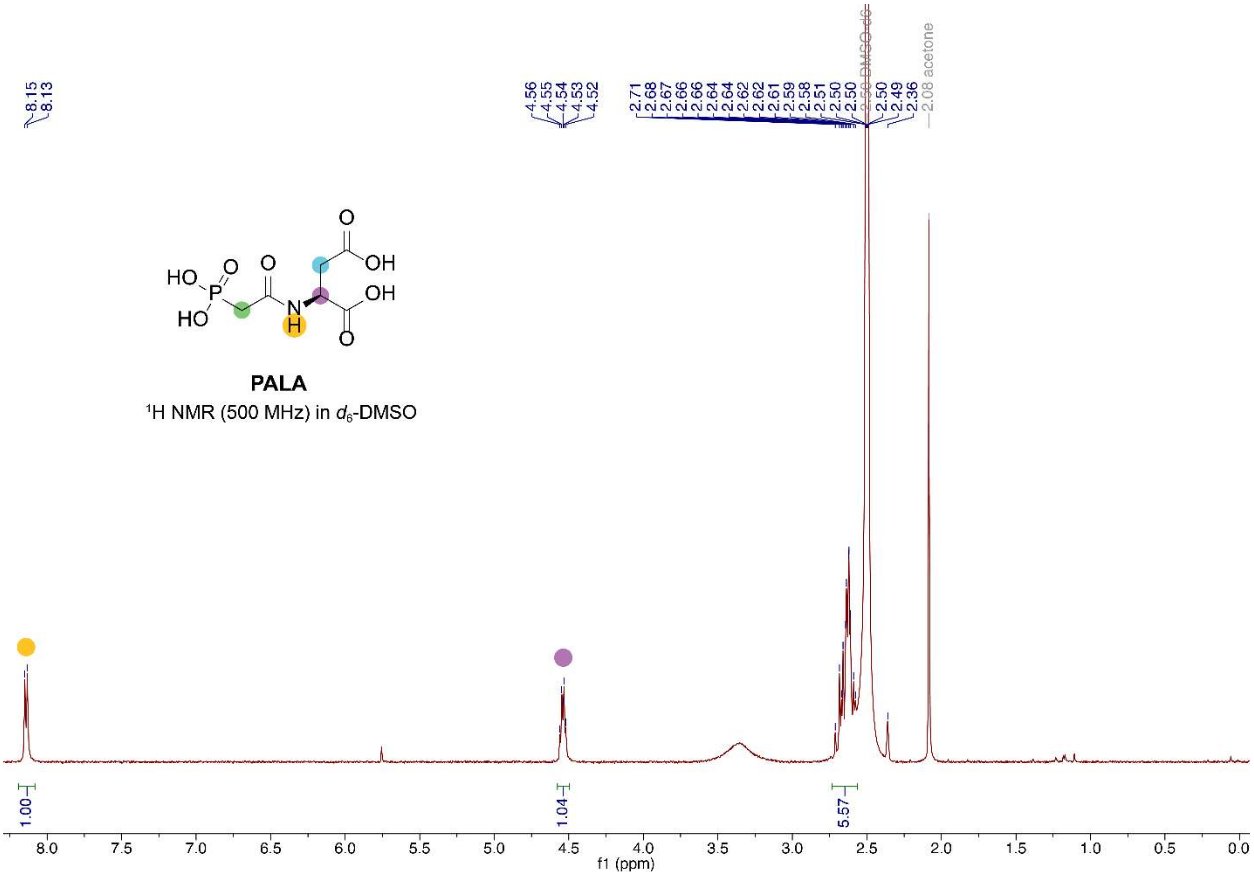


**Supplementary Figure 31. ¹H NMR spectrum of PALA in DMSO.**

Signals from the -$CH_2$- groups labeled in *green* and *blue* are highly overlapped with each other and partially submerged under the solvent peak, which are further resolved in the spectrum obtained in $D_2O$ (see Supplementary Fig. 32).

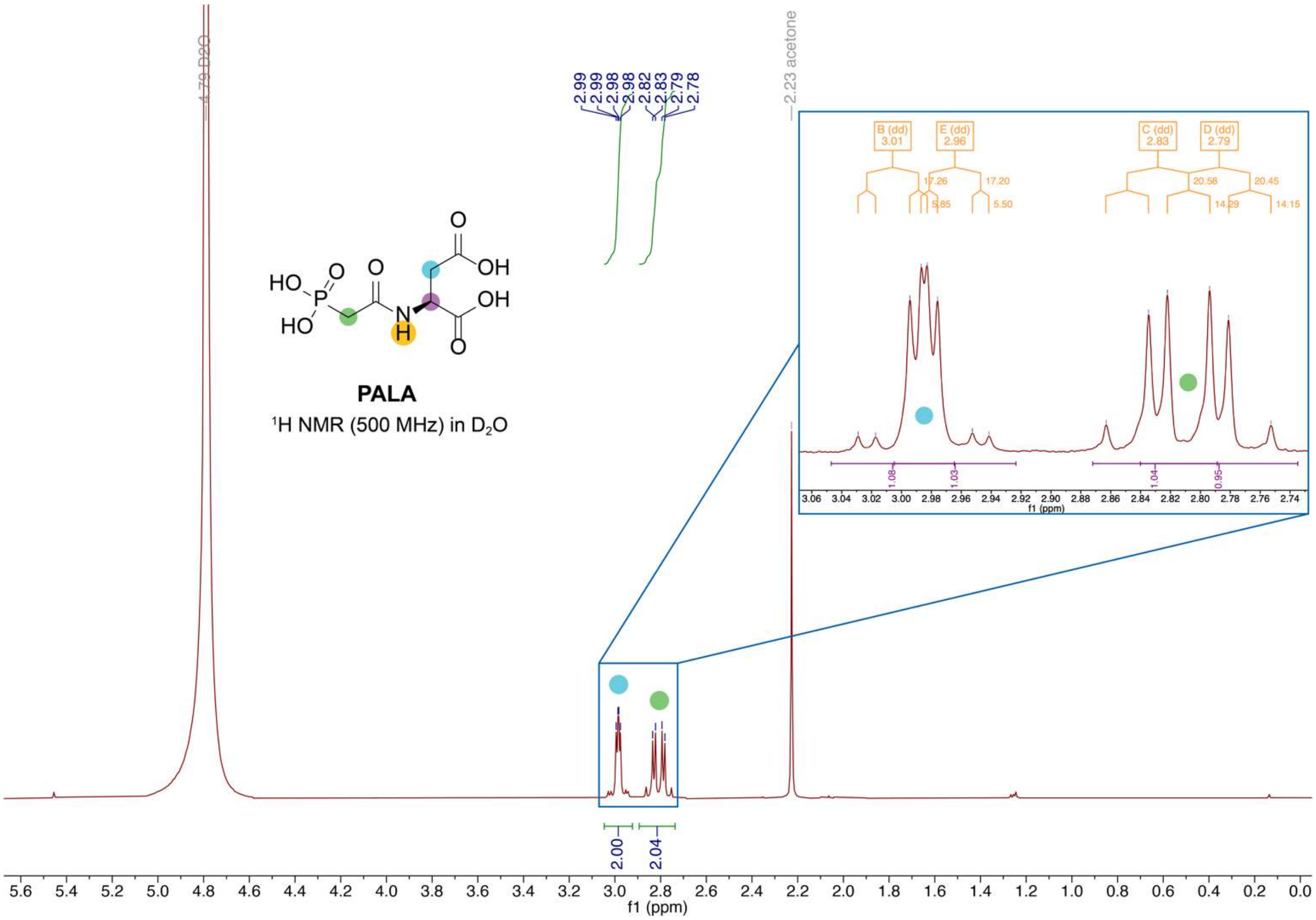


**Supplementary Figure 32. $^1$H NMR spectrum of PALA in $D_2O$.**

Signal from the hydrogen on chiral carbon (labelled in *purple*) is likely submerged under the $D_2O$ peak based on the previously reported chemical shift (4.70 ppm).[5] *Inset: J*-coupling trees and coupling constants of the two sets of peaks, showing how PALA being enantiopure makes the signals from two hydrogens on the same -$CH_2$- inequivalent.

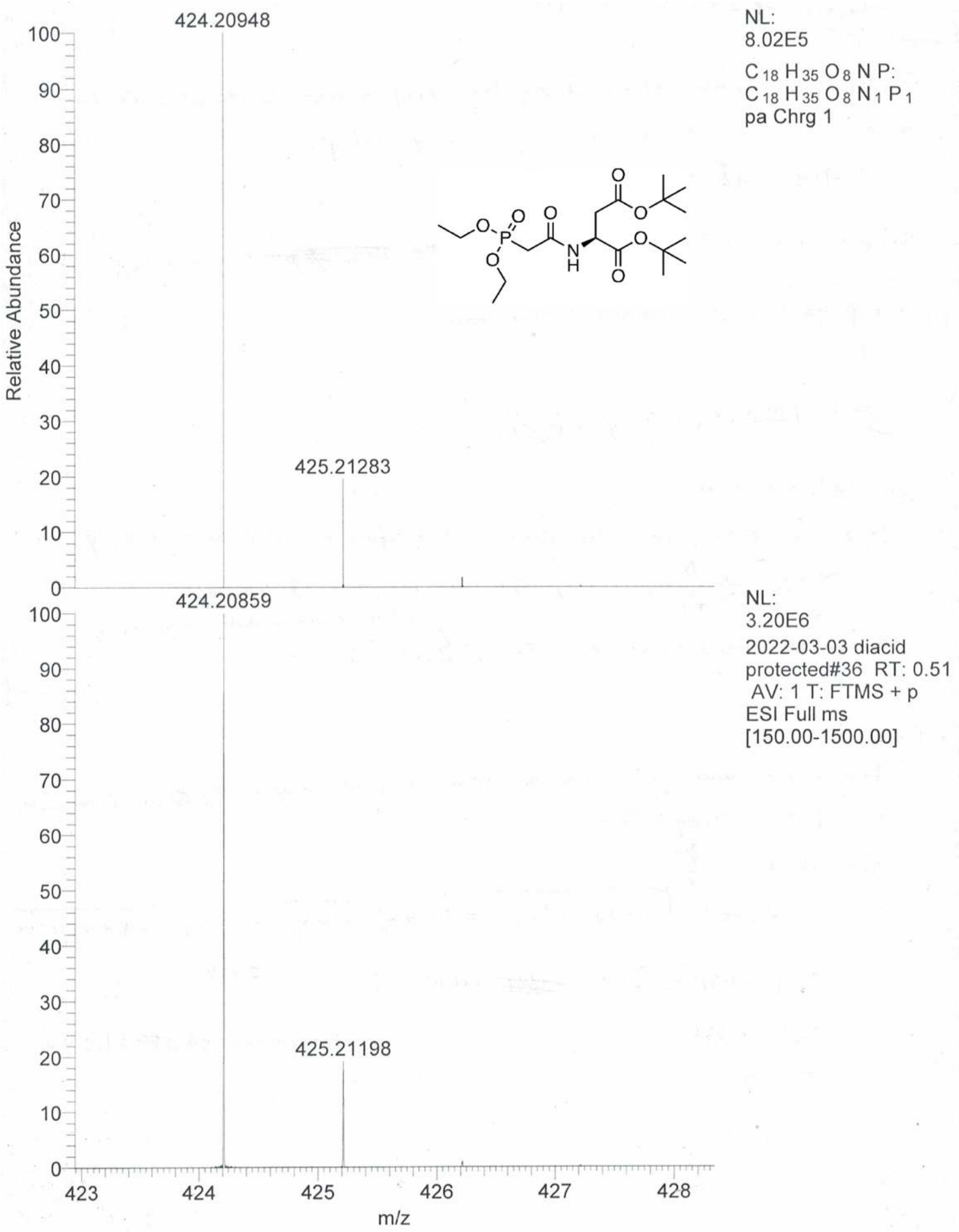


**Supplementary Figure 33. The HR-ESI mass spectra for the immediate precursor to PALA (compound 3 in Supplementary Figure 29).**

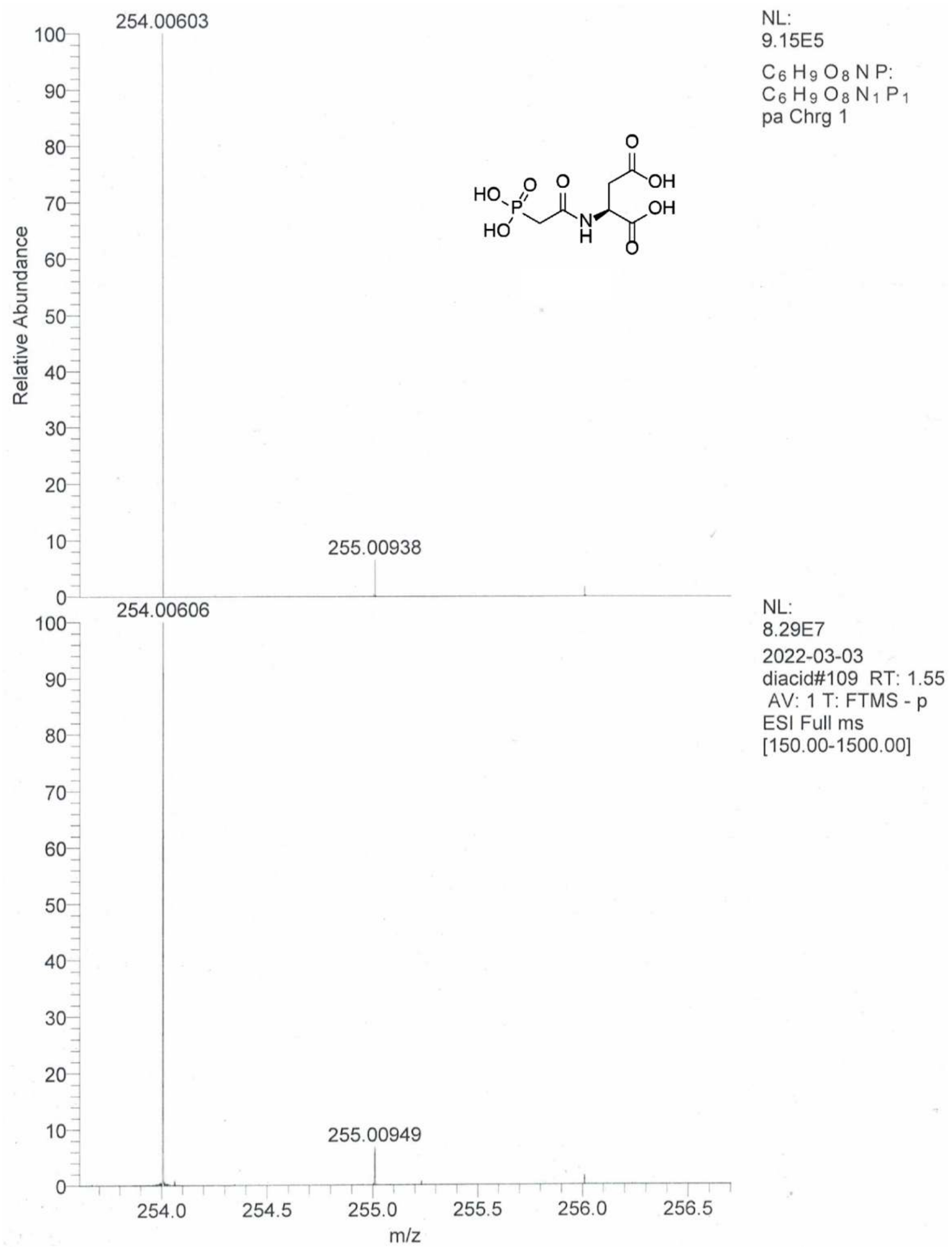


**Supplementary Figure 34. The HR-ESI mass spectra for PALA.**

#### Supplementary Table 1: SAXS data collection parameters for substrate and substrate analog titrations (Supp. Fig. 2).

| **SAXS data collection parameters** | | |
|---|---|---|
| *E. coli* ATCase concentration | 8 µM | 8 µM |
| Titrated ligand | 0 – 500 µM CP | 0 – 50 µM PALA |
| Buffer | 40 mM Tris-HCl pH 7.5, 15 mM $MgCl_2$, 1 mM TCEP, 10 mM succinate | 40 mM Tris-HCl pH 7.5, 15 mM $MgCl_2$, 1 mM TCEP |
| Beamline/detector | CHESS ID7A1/Eiger4M | CHESS ID7A1/Eiger4M |
| Energy (keV) | 9.81 | 9.93 |
| Beam size (µm) | 250 × 250 | 250 × 250 |
| Detector distance (mm) | 1602 | 1588 |
| $q$-measurement range ($Å^{-1}$) | 0.0108 – 0.524 | 0.0089 – 0.549 |
| Normalization | Transmitted intensity | Transmitted intensity |
| Exposures | 10 × 1 s | 10 × 1 s |
| Experimental temperature (°C) | 4 | 4 |

#### Supplementary Table 2: SAXS data collection parameters for pyrimidine titrations (Supp. Fig. 3).

| **SAXS data collection parameters** | | |
|---|---|---|
| *E. coli* ATCase concentration | 8 µM | 8 µM |
| Titrated ligand | 0 – 3 mM CTP | 0 – 5 mM CTP/UTP |
| Buffer | 40 mM Tris-HCl pH 7.5, 15 mM $MgCl_2$, 1 mM TCEP, 10 mM succinate, 0.5 mM CP | 40 mM Tris-HCl pH 7.5, 15 mM $MgCl_2$, 1 mM TCEP, 10 mM succinate, 0.5 mM CP |
| Beamline/detector | CHESS ID7A1/Eiger4M | CHESS ID7A1/Eiger4M |
| Energy (keV) | 10.08 | 10.08 |
| Beam size (µm) | 250 × 250 | 250 × 250 |
| Detector distance (mm) | 1606 | 1606 |
| $q$-measurement range ($Å^{-1}$) | 0.0087 – 0.547 | 0.0092 – 0.547 |
| Normalization | Transmitted intensity | Transmitted intensity |
| Exposures | 10 × 1 s | 10 × 1 s |
| Experimental temperature (°C) | 4 | 4 |

#### Supplementary Table 3: SAXS data collection parameters for purine titrations (Supp. Fig. 3).

| **SAXS data collection parameters** | | |
|---|---|---|
| *E. coli* ATCase concentration | 8 µM | 8 µM |
| Titrated ligand | 0 – 4 mM ATP | 0 – 3 mM GTP |
| Buffer | 40 mM Tris-HCl pH 7.5, 15 mM $MgCl_2$, 1 mM TCEP, 10 mM succinate, 0.5 mM CP | 40 mM Tris-HCl pH 7.5, 15 mM $MgCl_2$, 1 mM TCEP, 10 mM succinate, 0.5 mM CP, 5 mM ATP |
| Beamline/detector | CHESS ID7A1/Eiger4M | CHESS ID7A1/Eiger4M |
| Energy (keV) | 9.81 | 10.08 |
| Beam size (µm) | 250 × 250 | 250 × 250 |
| Detector distance (mm) | 1602 | 1606 |
| $q$-measurement range ($Å^{-1}$) | 0.0108 – 0.524 | 0.0092 – 0.547 |
| Normalization | Transmitted intensity | Transmitted intensity |
| Exposures | 10 × 1 s | 10 × 1 s |
| Experimental temperature (°C) | 4 | 4 |

**Supplementary Table 4: SEC-SAXS data collection parameters for ligand-free T-state and nucleotide-free R-state (Supp. Fig. 4)**

| | | |
|---|---|---|
| **SAXS data collection parameters** | | |
| Loading *E. coli* ATCase concentration | 56 µM | 56 µM |
| Running buffer | 40 mM Tris-HCl pH 7.5, 15 mM $MgCl_2$, 1 mM TCEP | 40 mM Tris-HCl pH 7.5, 15 mM $MgCl_2$, 1 mM TCEP, 10 mM succinate, 0.5 mM CP |
| Beamline/detector | CHESS ID7A1/Eiger4M | CHESS ID7A1/Eiger4M |
| Energy (keV) | 10.00 | 9.95 |
| Beam size (µm) | 250 × 250 | 250 × 250 |
| Detector distance (mm) | 1648 | 1623 |
| $q$-measurement range (Å $^{-1}$) | 0.0086 – 0.466 | 0.0080 – 0.525 |
| Normalization | Transmitted intensity | Transmitted intensity |
| Exposures | 1393 × 2 s | 1500 × 2 s |
| Column (flow rate, ml/min) | Superdex 200 10/300 GL (0.5) | Superdex 200 10/300 GL (0.6) |
| Experimental temperature (°C) | 4 | 4 |
| **REGALS analysis** | | |
| Buffer range (frames) | 109 – 187 | -24 – 57 |
| Main component range (frames modeled) | 174 – 600 | 185 – 450 |
| **Guinier analysis of main component** | | |
| $I(0)$ | 5.76 | 6.12 |
| $R_g$ (Å) | 48.2 ± 0.2 | 49.7 ± 2.6 |
| $qR_g$ range | 0.49 – 1.29 | 0.51 – 1.27 |

*Frames were renumbered to reflect the data range used for REGALS analysis.

#### Supplementary Table 5: SEC-SAXS data collection parameters for pyrimidine-bound R-states (Supp. Fig. 5)

| **SAXS data collection parameters** | | |
|---|---|---|
| Loading *E. coli* ATCase concentration | 56 µM | 56 µM |
| Running buffer | 40 mM Tris-HCl pH 7.5, 15 mM $MgCl_2$, 1 mM TCEP, 10 mM succinate, 0.5 mM CP, 1.5 mM CTP | 40 mM Tris-HCl pH 7.5, 15 mM $MgCl_2$, 1 mM TCEP, 10 mM succinate, 0.5 mM CP, 1.5 mM CTP/UTP |
| Beamline/detector | CHESS ID7A1/Eiger4M | CHESS ID7A1/Eiger4M |
| Energy (keV) | 9.95 | 11.34 |
| Beam size (µm) | 250 × 250 | 250 × 250 |
| Detector distance (mm) | 1717 | 1755 |
| $q$-measurement range (Å $^{-1}$) | 0.0095 – 0.481 | 0.0079 – 0.453 |
| Normalization | Transmitted intensity | Transmitted intensity |
| Exposures | 1018 × 2 s | 973 × 3 s |
| Column (flow rate, ml/min) | Superdex 200 10/300 GL (0.5) | Superdex 200 10/300 GL (0.3) |
| Experimental temperature (°C) | 4 | 4 |
| **REGALS analysis** | | |
| Buffer range (frames) | 60 – 101 | -14 – 38 |
| Main component range (frames modeled) | 195 – 450 | 215 – 450 |
| **Guinier analysis of main component** | | |
| $I(0)$ | 6.16 | 5.68 |
| $R_g$ (Å) | 49.1 ± 0.6 | 49.0 ± 0.5 |
| $qR_g$ range | 0.66 – 1.30 | 0.62 – 1.30 |

*Frames were renumbered to reflect the data range used for REGALS analysis.

### Supplementary Table 6: SEC-SAXS data collection parameters for purine-bound R-states (Supp. Fig. 6)

| **SAXS data collection parameters** | | |
|---|---|---|
| Loading *E. coli* ATCase concentration | 56 µM | 56 µM |
| Running buffer | 40 mM Tris-HCl pH 7.5, 15 mM $MgCl_2$, 1 mM TCEP, 10 mM succinate, 0.5 mM CP, 5 mM ATP | 40 mM Tris-HCl pH 7.5, 15 mM $MgCl_2$, 1 mM TCEP, 10 mM succinate, 0.5 mM CP, 5 mM ATP, 1 mM GTP |
| Beamline/detector | CHESS ID7A1/Eiger4M | CHESS ID7A1/Eiger4M |
| Energy (keV) | 9.95 | 9.95 |
| Beam size (µm) | 250 × 250 | 250 × 250 |
| Detector distance (mm) | 1623 | 1717 |
| $q$-measurement range (Å $^{-1}$) | 0.0080 – 0.525 | 0.0095 – 0.481 |
| Normalization | Transmitted intensity | Transmitted intensity |
| Exposures | 1200 × 2 s | 1006 × 2 s |
| Column (flow rate, ml/min) | Superdex 200 10/300 GL (0.2) | Superdex 200 10/300 GL (0.2) |
| Experimental temperature (°C) | 4 | 4 |
| **REGALS analysis** | | |
| Buffer range (frames) | -2 – 33 | -8 – 33 |
| Main component range (frames modeled) | 215 – 450 | 185 – 450 |
| **Guinier analysis of main component** | | |
| $I(0)$ | 6.21 | 6.37 |
| $R_g$ (Å) | 49.9 ± 0.7 | 51.2 ± 0.7 |
| $qR_g$ range | 0.48 – 1.28 | 0.69 – 1.29 |

*Frames were renumbered to reflect the data range used for REGALS analysis.

### Supplementary Table 7: EM data processing and refinement statistics for ligand-free T-state and states with only CP bound

| | **No ligands** | **CP, CTP** | **CP, CTP/UTP** | **CP, ATP** | **CP, ATP/GTP** |
|---|---|---|---|---|---|
| EMDB ID | EMD-47956 | EMD-47959 | EMD-47957 | EMD-47966 | EMD-47965 |
| PDB ID | 9EEK | 9EEN | 9EEL | 9EEU | 9EES |
| **Data collection** | | | | | |
| Software | | | | | |
| Collection | Serial-EM | Leginon | Leginon | Leginon | Leginon |
| Processing | Relion | Relion | Relion | Relion | Relion |
| Microscope | Talos Arctica | Titan Krios | Titan Krios | Titan Krios | Titan Krios |
| Camera | K3 | K3 | K3 | K3 | K3 |
| Nominal Magnification | 63,000 | 81,000 | 81,000 | 105,000 | 105,000 |
| Voltage (keV) | 200 | 300 | 300 | 300 | 300 |
| Electron exposure ($e^-/Å^2$) | 50.00 | 51.75 | 51.75 | 63.45 | 63.45 |
| Defocus range (μm) | -0.6 to -2.0 | -0.6 to -2.3 | -0.6 to -2.3 | -0.6 to -2.3 | -0.6 to -2.3 |
| Pixel size (Å) | 1.310 | 1.069 | 1.069 | 0.826 | 0.826 |
| Micrographs used (no.) | 1,601 | 4,012 | 4,471 | 6,390 | 6,390 |
| Final particles (no.) | 565,000 | 85,000 | 290,000 | 207,000 | 163,000 |
| Symmetry imposed | C3 | C3 | C3 | C3 | C3 |
| Map | | | | | |
| Resolution§ (Å) | 2.95 | 3.84 | 3.55 | 2.97 | 3.25 |
| Resolution range (Å) | 2.43 – 34.51 | 3.47 – 12.89 | 3.22 – 8.61 | 2.63 - 30.42 | 2.76 – 17.18 |
| Sharpening B-factor ($Å^2$) | 288.6 | 123.7 | 154.7 | 93.1 | 117.3 |
| **Model refinement** | | | | | |
| Resolution† (Å) | 3.0 | 4.1 | 3.6 | 3.0 | 3.4 |
| Model composition | | | | | |
| Non-H atoms | 21,174 | 21,606 | 21,804 | 21,858 | 21,984 |
| Protein residues | 2,712 | 2,718 | 2,726 | 2,748 | 2,760 |
| Ligands | 6 | 30 | 30 | 30 | 30 |
| B-factors ($Å^2$) | | | | | |
| Protein | 86.08 | 164.13 | 170.00 | 140.61 | 104.93 |
| Ligands | 86.82 | 199.53 | 143.99 | 123.78 | 131.63 |
| R.M.S deviations | | | | | |
| Bond lengths (Å) | 0.001 | 0.003 | 0.003 | 0.002 | 0.003 |
| Bond angles (°) | 0.373 | 0.561 | 0.577 | 0.434 | 0.517 |
| Validation | | | | | |
| *CC* (mask, volume) | 0.82, 0.81 | 0.74, 0.74 | 0.81, 0.80 | 0.86, 0.84 | 0.76, 0.74 |
| Overall Q-score | 0.48 | 0.30 | 0.36 | 0.51 | 0.42 |
| MolProbity score | 1.40 | 2.07 | 1.90 | 1.38 | 1.66 |
| Clashscore | 7.25 | 10.93 | 9.06 | 6.93 | 9.26 |
| Rotamer outliers (%) | 0.77 | 2.53 | 1.70 | 0.64 | 0.76 |
| Ramachandran plot | | | | | |
| Disallowed (%) | 0.00 | 0.15 | 0.11 | 0.00 | 0.00 |
| Allowed (%) | 1.97 | 3.16 | 3.53 | 1.80 | 2.96 |
| Favored (%) | 98.03 | 96.70 | 96.36 | 98.20 | 97.04 |
| Ramachandran Z-scores | | | | | |
| Overall (r.m.s.d.) | 1.32 (0.17) | -0.21 (0.16) | 0.07 (0.16) | 0.62 (0.16) | 0.29 (0.17) |
| Helix (r.m.s.d.) | 1.41 (0.17) | 0.80 (0.17) | 0.93 (0.17) | 1.16 (0.17) | 1.21 (0.18) |
| Sheet (r.m.s.d.) | 1.40 (0.21) | -0.18 (0.29) | -0.58 (0.23) | 0.44 (0.23) | 0.25 (0.23) |
| Loop (r.m.s.d.) | 0.22 (0.21) | -0.69 (0.17) | -0.19 (0.18) | -0.02 (0.17) | -0.43 (0.18) |

All entries are in a closed T-state conformation except for the last column, which corresponds to the ATP/GTP-bound pre-Asp R-state. FSC thresholds for the reported resolutions are 0.143 (§), and 0.500 (†).

### Supplementary Table 8: EM data processing and refinement statistics for R-states with both CP and succinate bound

| | No nucleotide | CTP | CTP/UTP | ATP | ATP/GTP |
|---|---|---|---|---|---|
| EMDB ID | EMD-47961 | EMD-47960 | EMD-47958 | EMD-47963 | EMD-47964 |
| PDB ID | 9EEP | 9EEO | 9EEM | 9EEQ | 9EER |
| **Data collection** | | | | | |
| Software | | | | | |
| Collection | Leginon | Leginon | Leginon | Leginon | Leginon |
| Processing | Relion | Relion | Relion | Relion | Relion |
| Microscope | Titan Krios | Titan Krios | Titan Krios | Titan Krios | Titan Krios |
| Camera | K3 | K3 | K3 | K3 | K3 |
| Nominal Magnification | 81,000 | 81,000 | 81,000 | 81,000 | 81,000 |
| Voltage (keV) | 300 | 300 | 300 | 300 | 300 |
| Electron exposure ($e^-/Å^2$) | 50.47 | 51.75 | 51.75 | 52.47 | 52.47 |
| Defocus range (µm) | -0.6 to -2.3 | -0.6 to -2.3 | -0.6 to -2.3 | -0.6 to -2.3 | -0.6 to -2.3 |
| Pixel size (Å) | 0.534 | 1.069 | 1.069 | 1.068 | 1.068 |
| Micrographs used (no.) | 3849 | 4,012 | 4,471 | 4029 | 4471 |
| Final particles (no.) | 682,000 | 450,000 | 830,000 | 934,000 | 865,000 |
| Symmetry imposed | C1 | C1 | C1 | C1 | C1 |
| Map | | | | | |
| Resolution§ (Å) | 3.08 | 3.53 | 3.05 | 3.27 | 3.27 |
| Resolution range (Å) | 2.43 – 34.51 | 3.17 – 19.34 | 2.67 – 11.65 | 2.82 – 9.81 | 2.74 – 11.08 |
| Sharpening B-factor ($Å^2$) | 99.9 | 98.5 | 109.25 | 104.8 | 102.3 |
| **Model refinement** | | | | | |
| Resolution† (Å) | 3.3 | 3.9 | 3.4 | 3.5 | 3.4 |
| Model composition | | | | | |
| Non-H atoms | 21270 | 21,624 | 21,852 | 21,732 | 22,032 |
| Protein residues | 2712 | 2,712 | 2,742 | 2,724 | 2,760 |
| Ligands | 18 | 36 | 36 | 36 | 36 |
| B-factors ($Å^2$) | | | | | |
| Protein | 71.59 | 100.61 | 70.05 | 62.82 | 74.51 |
| Ligands | 62.49 | 119.35 | 60.87 | 85.81 | 89.36 |
| R.M.S deviations | | | | | |
| Bond lengths (Å) | 0.002 | 0.002 | 0.002 | 0.002 | 0.002 |
| Bond angles (°) | 0.440 | 0.469 | 0.476 | 0.500 | 0.475 |
| Validation | | | | | |
| *CC* (mask, volume) | 0.71, 0.70 | 0.70, 0.69 | 0.70, 0.69 | 0.70, 0.68 | 0.74, 0.73 |
| Overall Q-score | 0.42 | 0.32 | 0.42 | 0.39 | 0.42 |
| MolProbity score | 1.68 | 1.90 | 1.64 | 1.57 | 1.49 |
| Clashscore | 8.84 | 11.55 | 9.20 | 10.53 | 9.25 |
| Rotamer outliers (%) | 0.99 | 0.60 | 0.98 | 0.43 | 0.76 |
| Ramachandran plot | | | | | |
| Disallowed (%) | 0.0 | 0.00 | 0.00 | 0.00 | 0.15 |
| Allowed (%) | 3.24 | 4.58 | 2.83 | 2.11 | 1.75 |
| Favored (%) | 96.76 | 95.42 | 97.17 | 97.89 | 98.25 |
| Ramachandran Z-scores | | | | | |
| Overall (r.m.s.d.) | 0.37 (0.17) | -0.46 (0.16) | 0.37 (0.16) | 0.68 (0.17) | 0.66 (0.17) |
| Helix (r.m.s.d.) | 0.85 (0.18) | 0.47 (0.17) | 0.92 (0.18) | 1.24 (0.18) | 1.36 (0.18) |
| Sheet (r.m.s.d.) | 0.12 (0.23) | -0.81 (0.21) | -0.32 (0.21) | 0.49 (0.23) | 0.29 (0.23) |
| Loop (r.m.s.d.) | -0.03 (0.19) | -0.52 (0.18) | 0.21 (0.18) | -0.03 (0.18) | -0.05 (0.18) |

All entries correspond to an open R-state. FSC thresholds for the reported resolutions are 0.143 (§), and 0.500 (†).

#### Supplementary Table 9: Kinetic parameters from Hill and Michaelis-Menten models

| Condition | Hill fit | | | Michaelis-Menton | |
|---|---|---|---|---|---|
| | $n_H$ | $K_{1/2}$ (mM) | $V_{max}$ | $K_m$ (mM) | $V_{max}$ |
| CTP/UTP | 2.9 ± 0.2 | 37 ± 1 | 5.7 ± 0.1 | ---- | ---- |
| CTP | 2.6 ± 0.1 | 18.3 ± 0.5 | 9.9 ± 0.2 | ---- | ---- |
| No NTP | 2.5 ± 0.2 | 15.4 ± 0.5 | 9.2 ± 0.2 | ---- | ---- |
| ATP | 2.1 ± 0.1 | 5.6 ± 0.2 | 9.4 ± 0.2 | ---- | ---- |
| ATP/GTP | 1.21 ± 0.03 | 2.89 ± 0.09 | 12.6 ± 0.2 | 4.5 ± 0.2 | 13.6 ± 0.3 |

Saturation curves of *E. coli* ATCase have historically been modeled by the Hill model (Eq. 1). Under the assumption that no cooperativity ($n_H = 1$), the Hill model reduces to the Michaelis-Menton model (Eq. 2). Maximum velocity ($V_{max}$) values are reported in units of mmol $hr^{-1}$ $mg^{-1}$.

#### Supplementary Table 10: Nucleotide saturation in *E. coli* ATCase and estimates of physiological nucleotide concentrations

| | Saturation [NTP] (mM) | Physiological [NTP] (mM) | |
|---|---|---|---|
| CTP | ~1.5 | 0.325 | 1.2 – 2.7 |
| UTP | ~1 | 0.667 | 2.4 – 8.3 |
| ATP | ~3 | 3.56 | 4.1 – 9.6 |
| GTP | ~0.25 | 1.66 | 1.3 – 4.9 |

Nucleotide saturation values were obtained from singular value decomposition of SAXS nucleotide titrations (refer to Supplementary Fig. 3). Estimated cellular concentrations of nucleotides from Buckstein, *et al.*[40] (left) and Bennett, *et al.*[41] (right).

### Supplementary Table 11: Crystallographic data collection and refinement statistics

| | ATP/GTP-bound R-state | ATP-bound R-state |
|---|---|---|
| **Data collection** | | |
| PDB ID | 9EEH | 9EEJ |
| Space group | $P\,3\,2\,1$ | $P\,2_1\,2_1\,2_1$ |
| Cell dimensions | | |
| a, b, c (Å) | 122.4, 122.4, 153.0 | 129.0, 148.8, 207.9 |
| α, β, γ (°) | 90, 90, 120 | 90, 90, 90 |
| Wavelength (Å) | 0.9185 | 0.9795 |
| Resolution (Å) | 62.05 – 2.19 (2.27 – 2.19) | 103.90 – 3.00 (3.11 – 3.00) |
| Reflections (total) | 352,397 (35,932) | 466,501 (48,201) |
| Reflections (unique) | 67,983 (6,691) | 80,502 (7,971) |
| $R_{merge}$ | 0.086 (0.808) | 0.2676 (1.027) |
| $R_{meas}$ | 0.095 (0.894) | 0.2947 (1.125) |
| $R_{pim}$ | 0.041 (0.378) | 0.1213 (0.454) |
| Mean ($I/\sigma I$) | 13.9 (1.3) | 4.2 (0.7) |
| $CC_{1/2}$ | 0.994 (0.705) | 0.966 (0.642) |
| Completeness (%) | 98.9 (98.3) | 99.3 (96.7) |
| Multiplicity | 5.2 (5.4) | 5.8 (6.0) |
| Wilson B-factor (Å$^2$) | 44.2 | 62.3 |
| **Refinement statistics** | | |
| Resolution (Å) | 62.05 – 2.19 | 120.99 – 3.00 |
| Reflections (total) | 67,947 | 80,134 |
| Reflections (test) | 1,364 | 1,992 |
| $R_{work}$, $R_{free}$ | 0.1460, 0.1686 | 0.2182, 0.2494 |
| Number of atoms | | |
| Protein | 7,182 | 21,457 |
| Ligands | 162 | 480 |
| Solvent | 321 | 0 |
| B-factors (Å$^2$) | | |
| Protein | 59.7 | 70.7 |
| Ligands | 82.1 | 95.0 |
| Solvent | 55.6 | - |
| R.M.S. deviations | | |
| Bond lengths (Å) | 0.008 | 0.002 |
| Bond angles (°) | 0.920 | 0.520 |
| MolProbity statistics | | |
| Ramachandran plot | | |
| favored, allowed, outliers (%) | 97.59, 2.41, 0.00 | 95.60, 4.11, 0.29 |
| Rotamer outliers | 2.03 | 3.22 |
| Clashscore | 2.05 | 6.11 |
| Ramachandran Z-scores | | |
| Overall (r.m.s.d.) | -0.50 (0.26) | -1.77 (0.14) |
| Helix (r.m.s.d.) | -0.56 (0.27) | -1.77 (0.13) |
| Sheet (r.m.s.d.) | 0.04 (0.34) | -0.59 (0.20) |
| Loop (r.m.s.d.) | -0.15 (0.31) | -0.84 (0.15) |

Statistics for the highest-resolution shell are shown in parentheses.

**Supplementary Table 12: Crystallographic data collection and phasing statistics**

| | ATP/GTP-bound R-state (expanded) |
|---|---|
| **Data collection** | |
| Space group | *P* 3 2 1 |
| Cell dimensions | |
| a, b, c (Å) | 126.2, 126.2, 166.8 |
| α, β, γ (°) | 90, 90, 120 |
| Wavelength (Å) | 0.9185 |
| Resolution (Å) | 166.80 – 3.50 (3.56 – 3.50) |
| Reflections (total) | 201,840 (10,006) |
| Reflections (unique) | 19,922 (970) |
| $R_{merge}$ | 0.116 (1.529) |
| $R_{meas}$ | 0.122 (1.609) |
| $R_{pim}$ | 0.039 (0.497) |
| Mean ($I/\sigma I$) | 9.1 (0.6) |
| $CC_{1/2}$ | 0.999 (0.783) |
| Completeness (%) | 100.0 (100.0) |
| Multiplicity | 10.1 (10.3) |
| Wilson B-factor (Å$^2$) | 119.10 |

Statistics for the highest-resolution shell are shown in parentheses. This crystal did not diffract to sufficiently high resolution for an unambiguous molecular replacement solution. Phasing was possible with multiple models, including all R-state EM models as well as the contracted ATP/GTP-bound R-state crystal structure (PDB: 9EEH), but the expanded ATP/GTP-bound R-state cryo-EM model (PDB: 9EER) yielded the best phasing statistics. The most definitive indicator is the increase in the unit cell *c*-dimension, which is parallel to the 3-fold axis of the holocomplex in the P321 structures: 153 Å for the contracted R-state versus 167 Å for the expanded R-state.
